# BundleWarp: Enhancing White Matter Tractometry and Morphometry with Precise Neuronal Mapping using Streamline-based Nonlinear Registration

**DOI:** 10.1101/2023.01.04.522802

**Authors:** Bramsh Qamar Chandio, Emanuele Olivetti, David Romero-Bascones, Sophia I. Thomopoulos, Julio E. Villalon-Reina, Talia M. Nir, Jaroslaw Harezlak, Paul M. Thompson, Eleftherios Garyfallidis

## Abstract

Tractometry analysis represents a significant advancement in neuroimaging, offering a detailed examination of the brain’s white matter at a micro level. Unlike traditional ROI or voxel-based methods, tractometry precisely reconstructs and characterizes white matter tracts. Using advanced diffusion MRI and tractography algorithms, it maps the trajectory, shape, and connectivity patterns of individual white matter bundles. Accurate alignment of these tracts across different groups is crucial for reliable and reproducible results. Nonlinear registration techniques are essential for achieving this alignment, harmonizing bundle shapes, and improving sensitivity to disease-related changes. However, nonlinear registration is complex, especially with tractography data, which digitally represents the brain’s white matter anatomy. Potential structural changes in the bundle’s shape during registration can lead to artifacts that obscure critical anatomical details needed for disease identification. We introduce BundleWarp, a streamline-based nonlinear deformable registration method designed specifically for white matter tracts. BundleWarp employs a sophisticated approach to align two white matter bundles while preserving their topological and anatomical features. It is formulated as a probability density estimation problem with motion coherence penalties, ensuring coherent movement of points along streamlines and maintaining the anatomical integrity of tracts through displacement field regularization. Additionally, we introduce a tract morphometry framework utilizing the displacement field generated by BundleWarp to analyze white matter tract shape differences. Our results show that BundleWarp effectively quantifies bundle shape differences and enhances structural harmonization in tractometry analysis for diverse subjects, including those with Alzheimer’s and Parkinson’s disease. Test–retest experiments further demonstrate that BundleWarp substantially improves subject finger-printing by increasing within-subject reproducibility of both bundle shape and microstructural profiles (FA, MD, RD, AD). It precisely maps the brain’s neuronal pathways, offering a robust tractometry framework with enhanced sensitivity for detecting disease-related structural and microstructural changes in white matter tracts associated with Mild Cognitive Impairment (MCI), dementia, and early-stage Alzheimer’s biomarkers, including amyloid-beta plaques and tau neurofibrillary tangles.

## 1. Introduction

Over the past few decades, diffusion-weighted MRI (dMRI) has steadily gained recognition as a valuable and widely researched MRI technique Basser, Mattiello and LeBihan (1994); Le Bihan, Mangin, Poupon, Clark, Pappata, Molko and Chabriat (2001). dMRI facilitates the reconstruction of white matter fibers and their connectivity within the brain, making it an essential tool for in vivo studies. The digital reconstruction of 3D white matter pathways enables researchers to detect subtle abnormalities or disruptions that may indicate certain neurological conditions. In group analysis, these digital pathways serve as quantitative measures that can be compared between healthy controls and individuals with diseases or disorders. By analyzing differences in white matter coherence, connectivity, or specific neural pathways across these groups, researchers can identify biomarkers or signatures associated with particular diseases. These biomarkers may include alterations in diffusion metrics such as fractional anisotropy (FA) or mean diffusivity (MD), which reflect changes in tissue microstructure and organization Smith, Jenkinson, Johansen-Berg, Rueckert, Nichols, Mackay, Watkins, Ciccarelli, Cader, Matthews et al. (2006); Bach, Laun, Leemans, Tax, Biessels, Stieltjes and Maier-Hein (2014); Chandio (2020); Yendiki, Panneck, Srinivasan, Stevens, Zöllei, Augustinack, Wang, Salat, Ehrlich, Behrens et al. (2011); De Santis, Drakesmith, Bells, Assaf and Jones (2014); Bells, Cercignani, Deoni, Assaf, Pasternak, Evans, Leemans and Jones (2011).

The role of white matter has been studied in many neurodegenerative and psychiatric disorders, including Alzheimer’s disease De Vos, Grierson, Ackerley and Miller (2008); Nasrabady, Rizvi, Goldman and Brickman (2018); Papuć and Rejdak (2018), Parkinson’s disease Dean III, Sojkova, Hurley, Kecskemeti, Okonkwo, Bendlin, Theisen, Johnson, Alexander and Gallagher (2016), Multiple Sclerosis (MS) Albert, Antel, Brück and Stadelmann (2007); Haines, Inglese and Casaccia (2011), and Schizophrenia Cassoli, Guest, Malchow, Schmitt, Falkai and Martins-de Souza (2015); Whitford, Ford, Mathalon, Kubicki and Shenton (2012); Kubicki, McCarley and Shenton (2005). For example, the white matter of patients with MS Prineas, Barnard, Revesz, Kwon, Sharer and Cho (1993) has been extensively studied, and MS is linked to the immune system attacking the myelin sheath surrounding axon bundles, causing inflammation, and resulting in a breakdown in the transmission of neuronal signals in people with the disorder Albert et al. (2007); Haines et al. (2011); Brück (2005). Furthermore, changes in myelin have also been linked with several other brain conditions and diseases De Vos et al. (2008); Nasrabady et al. (2018); Papuć and Rejdak (2018); Cassoli et al. (2015); Whitford et al. (2012); Kubicki et al. (2005); Dean III et al. (2016). Hence, it is imperative to study the changes in white matter tracts related to brain diseases as we can learn more about the psychophysiology of these conditions, but also potentially learn crucial information which may aid in providing better diagnostic as well and possible prognostic information for patients.

Tractometry is an advanced method in the field of diffusion MRI that offers several advantages over traditional region-of-interest (ROI)-based or voxel-based analyses. Tractometry focuses specifically on the properties of white matter tracts and allows for the examination of diffusion properties along the entire extent of a fiber tract. Unlike voxel-based analyses, which provide a broad overview of tissue properties without specific reference to the connectivity between regions, or ROI-based analyses, which measure average properties within pre-defined areas, tractometry evaluates the microstructural properties along the length of individual white matter tracts. Tractometry allows precise mapping of brain connectivity and white matter alterations. By focusing on specific tracts, tractometry improves the sensitivity and specificity of detecting white matter abnormalities, allowing researchers to pinpoint where along a tract changes occur, which is advantageous for longitudinal studies tracking subtle disease progression or developmental changes. Evaluating white matter integrity along the tract enables direct links between structural alterations and functional outcomes in conditions such as Alzheimer’s disease and multiple sclerosis.

However, the effectiveness of tractometry for group analysis relies heavily on the quality of the registration process. Tractometry is used to measure and compare the properties of white matter tracts across individuals, which is essential for understanding brain structure and connectivity. Accurate alignment of diffusion MRI data to a common anatomical framework is crucial for these comparisons. If registration is inaccurate, tracts may not align properly across subjects, leading to potential mismatches and misinterpretations of white matter properties. Such misalignment can skew data, obscure real differences, and even create false appearances of abnormalities, all due to incorrect tract positioning rather than actual variations in tract properties. Additionally, poor registration increases data variability, reducing the statistical power of the study and increasing the risk of false findings, either by missing real differences (false negatives) or suggesting differences that are not actually there (false positives). Thus, precise registration is critical for reliable tractometry analysis in group studies.

Nonlinear registration is generally considered more effective than linear registration due to its ability to handle complex deformations and adapt to individual anatomical differences. This flexibility and precision make it particularly valuable in the field of neuroimaging, where accurate mapping of brain structures is crucial. Nonlinear registration’s capacity to model the real, often non-uniform, anatomical variability between subjects ensures more precise alignment of brain structures. This level of accuracy is essential for integrating and harmonizing data across multiple sites and scanners in medical imaging studies. By accommodating local variations and minimizing artifacts or distortions that might be introduced by different scanning equipment, nonlinear registration facilitates the pooling of data from various sources. This is crucial for multi-center studies where consistency and comparability of data are paramount. Ultimately, nonlinear registration enhances the quality and reliability of analyses, helping researchers to draw more accurate conclusions about brain structure and function from complex datasets.

White matter fibers of the brain are computationally reconstructed from dMRI Basser et al. (1994); Le Bihan et al. (2001); Alexander, Lee, Lazar and Field (2007) by means of tractography methods Farquharson, Tournier, Calamante, Fabinyi, Schneider-Kolsky, Jackson and Connelly (2013); Catani and De Schotten (2008); Gong, He, Concha, Lebel, Gross, Evans and Beaulieu (2008); Mori and Van Zijl (2002). A whole-brain tractography comprised of millions of digitally reconstructed white matter fibers from a single subject is called a tractogram. A single digital white matter fiber pathway is called a streamline, and a set of streamlines that travel together and share similar shapes, geometry, and functions is called a tract/bundle. In the context of streamlines, the geometry of a streamline refers to its local spatial configuration and physical shape (e.g., curvature, length, and orientation), while topology refers to the preserved order and connectivity of points along the streamline and across the bundle. For the segmentation of white matter bundles from whole-brain tractograms and their inter or intra-group analysis, data need to be aligned to a common reference. Similar to image-based group analysis pipelines Smith et al. (2006); Bach et al. (2014), streamline-based group analysis pipelines Chandio, Risacher, Pestilli, Bullock, Yeh, Koudoro, Rokem, Harezlak and Garyfallidis (2020); Yeatman, Dougherty, Myall, Wandell and Feldman (2012); Cousineau, Jodoin, Garyfallidis, Côté, Morency, Rozanski, Grand’Maison, Bedell and Descoteaux (2017); Dayan, Monohan, Pandya, Kuceyeski, Nguyen, Raj and Gauthier (2015); Chandio, Chattopadhyay, Owens-Walton, Reina, Nabulsi, Thomopoulos, Garyfallidis and Thompson (2022) also require registration as an intermediate step. Unlike image and voxel-based volumetric white matter tracts Hansen, Yang, Lyu, Rheault, Kerley, Chandio, Fadnavis, Williams, Shafer, Resnick et al. (2021), streamline-based white matter tracts Yeh, Panesar, Fernandes, Meola, Yoshino, Fernandez-Miranda, Vettel and Verstynen (2018a); Catani and de Schotten (2012) provide rich details about the brain’s white matter structural connectivity at the sub-millimeter scale as different voxels are integrated to generate streamlines. However, streamline-based registration is much more complicated. Streamline-based registration would require finding correspondences between streamlines of two bundles to map/match those corresponding streamlines and minimize the distance between bundles.

Tractography data comes with its challenges. Very often, two bundles will have a different number of streamlines in them, and mapping one streamline from source/moving to one streamline in a reference/static (one-to-one mapping) approach would not work. If the moving bundle has more streamlines than the static bundle, some of the streamlines in the moving bundle will be left unmapped. In the opposite case, when static has more streamlines than moving bundles, some of the streamlines of static will not be utilized in registration. Moreover, bundles of the same type in a healthy population usually have a similar overall shape, but streamlines within the bundles could have different lengths and geometries. A pure nonlinear streamline-based registration would require producing a warp/transformation matrix specific to each streamline and each point on the streamline. The desired output would be such that it closely aligns two bundles without excessively deviating from the original anatomical shape of the bundle or creating spurious streamlines.

The most common approach to aligning tractograms/tracts to a common space is to register the subject’s FA image to a template FA image and then apply the computed transformations (affine or deformable) to streamlines. Most commonly, ANTs’ nonlinear SyN registration Avants, Tustison and Song (2009) is performed on FA images, and resulting affine and warp maps are applied to streamlines of that subject. This process is also known as spatial normalization of tractography/streamlines Greene, Cieslak and Grafton (2018). However, image-based transformation maps do not consider the finer details of the tractography. Those transformations are derived from images and are not specific to streamlines, i.e., they do not consider neighborhood information on streamlines. Each streamline can have a different number of points and Euclidean length. Every streamline requires a deformation that is based on its geometry. Each point on the streamline needs to be warped in a manner such that the topological order of the points on the streamlines is preserved after the registration. Hence, it is desirable to have a robust streamline-based registration method for aligning tractography data directly in the streamline-space.

Several streamline-based registration methods have been proposed recently. Most of them fall in the category of linear registration Leemans, Sijbers, De Backer, Vandervliet and Parizel (2006); Durrleman, Fillard, Pennec, Trouvé and Ayache (2011); Durrleman, Prastawa, Charon, Korenberg, Joshi, Gerig and Trouvé (2014); Zvitia, Mayer, Shadmi, Miron and Greenspan (2009); Garyfallidis, Ocegueda, Wassermann and Descoteaux (2015); O’Donnell, Wells, Golby and Westin (2012). However, literature on nonlinear streamline-based registration methods is scarce, and very few methods are purely streamline-based Arsigny, Commowick, Ayache and Pennec (2009); Wassermann, Rathi, Bouix, Kubicki, Kikinis, Shenton and Westin (2011); Glozman, Bruckert, Pestilli, Yecies, Guibas and Yeom (2018); Lizarraga, Lee, Kubicki, Sahib, Nunez, Narr and Joshi (2021); St-Jean, Chamberland, Viergever and Leemans (2019); Olivetti, Gori, Astolfi, Bertó and Avesani (2020); Zhang, Wells and O’Donnell (2021). For example, Arsigny et al. (2009) generates scalar volumes from streamlines and then warps those volumes, applying the resulting transformations to streamlines. Similarly, Wassermann et al. (2011) uses tract density images to register bundles. Another method Glozman et al. (2018) implements a seed point matching strategy to establish correspondences between regions on the bundles, performing registration independently within those regions. A deep-learning-based approach Zhang et al. (2021) registers tract orientation maps (TOMs) and whole-brain FA images, where each voxel in a TOM file represents the local mean fiber orientation (one peak). While TOM files provide local mean fiber direction (peak), they do not include streamline connectivity information, i.e., which points belong to the same streamline or which points are neighbors along the same path. Another recently proposed method for nonlinear streamline registration uses deformation transfer via parallel transport Lizarraga et al. (2021), finding and applying transformations in latent space before reconstructing streamlines. A bundle profile–based approach has also been proposed for streamline registration using values of microstructural measures such as FA, where FA values are projected onto the streamlines and represented as a 1D list of pointwise diffusion values St-Jean et al. (2019).

Although these approaches capture local or regional deformations, they do not consider the topological connectivity of points along streamlines in the registration process. Recently, Olivetti et al. (2020) proposed a nonlinear streamline-matching framework that finds nonrigid correspondence directly between streamlines of two whole-brain tractograms. While it does not apply a spatial warp or generate a deformation field, it represents a key step toward direct streamline-based nonlinear registration of white matter tracts.

In our prior work, we introduced a preliminary streamline-based nonlinear registration framework (StnD) Chandio and Garyfallidis (2020), which first applies streamline-based linear registration (SLR) Garyfallidis et al. (2015) for coarse alignment and then models the bundles as point clouds for nonrigid registration using Coherent Point Drift (CPD) Myronenko and Song (2010). While this approach enables direct nonlinear alignment in streamline space, representing bundles as unordered point clouds disregards the inherent connectivity and ordering of points along individual streamlines. Consequently, this may lead to suboptimal alignment and reduced anatomical fidelity when mapping the deformation back to streamlines. Additionally, the computational cost can increase for dense bundles due to the point cloud representation. These limitations motivate the need for a framework that explicitly preserves streamline structure while enabling efficient and flexible nonlinear registration.

In this paper, building on our previous work on the direct alignment of tractograms/tracts in streamline-space Garyfallidis et al. (2015); Olivetti et al. (2020); Chandio and Garyfallidis (2020), we introduce a new method, BundleWarp. BundleWarp is a method for streamline-based non-linear registration of white matter tracts. Given two bundles, a static and a moving bundle, BundleWarp first linearly aligns two bundles using SLR Garyfallidis et al. (2015). It then finds pairs of matching streamlines in two bundles to establish correspondence among the streamlines of two bundles using iterLAP. iterLAP is a proposed extension of the rectangular sum of linear assignment problem (rLAP) for many-to-one streamline matching when static and moving bundles do not have the same number of streamlines. Finally, it warps the matching pair of streamlines using the proposed memoryless coherent point drift (mlCPD) method Myronenko, Song and Carreira-Perpinan (2006); Myronenko and Song (2010). The velocity field is regularized Myronenko and Song (2010); Yuille and Grzywacz (1989) using motion coherence theory Yuille and Grzywacz (1988) to coherently move points on the streamline as a group to maintain the topological structure of the streamlines.

BundleWarp provides users with an option to control the level of deformations in the registration through a single regularization parameter *λ*. Given two bundles, static and moving, BundleWarp can *partially* deform the moving bundle with a high value of *λ* or *fully* deform the moving bundle using a low value of *λ*. By default, BundleWarp partially deforms the bundle. We show that partially deforming the bundle improves the linear registration and, at the same time, preserves the original anatomical shape of the input bundle. We also show that nonlinearly (partially warped) aligned bundles of groups and populations to an atlas in standard space improve tract correspondence among subjects, resulting in robust statistical analysis of them. Additionally, we propose quantifying bundle shape differences through the nonlinear displacement field generated by fully deforming the moving bundle to precisely match the static bundle. To the best of our knowledge, BundleWarp is the only streamline-based white matter tract registration method that provides users with an option to control the level of deformations in the registration.

To evaluate the performance and reliability of BundleWarp registration, we applied it to 1,728 pairs of bundles, including 27 different types, extracted from Parkinson’s Progression Markers Initiative (PPMI) data Marek, Jennings, Lasch, Siderowf, Tanner, Simuni, Coffey, Kieburtz, Flagg, Chowdhury et al. (2011). We compared BundleWarp with other streamline-based linear and image-based nonlinear registration methods. Our results demonstrate that BundleWarp outperforms these methods based on both qualitative (visual) assessments and three quantitative metrics: bundle-based minimum distance, bundle shape similarity score, and volumetric DICE coefficient.

We demonstrate an important application of BundleWarp in tractometry by the integration of BundleWarp non-linear registration in BUndle ANalytics (BUAN) tractometry Chandio (2020), highlighting its effectiveness in enhancing the reliability of group-level tract analysis. By aligning tracts with BundleWarp, we improve the consistency of segment correspondence along white matter tracts across subjects, ensuring that corresponding segments represent the same anatomical regions in the bundles. To assess BundleWarp’s impact on tractometric analysis, we evaluate its impact on detecting white matter tract differences in subjects with mild cognitive impairment (MCI) and dementia and their associations with early-stage biomarkers of Alzheimer’s disease (AD), including amyloid and tau Goedert, Spillantini and Crowther (1991); Sepulcre, Grothe, d’Oleire Uquillas, Ortiz-Terán, Diez, Yang, Jacobs, Hanseeuw, Li, El-Fakhri et al. (2018); Villemagne, Doré, Burnham, Masters and Rowe (2018). By incorporating amyloid and tau, which are early pathological hallmarks of AD manifesting years before clinical symptoms, we demonstrate BundleWarp’s ability to capture subtle effects even in the preclinical stages of the disease.

Using data from 730 ADNI3 (phase 3 of the Alzheimer’s Disease Neuroimaging Initiative) dataset Zavaliangos-Petropulu, Nir, Thomopoulos, Reid, Bernstein, Borowski, Jack Jr., Weiner, Jahanshad and Thompson (2019) subjects, we analyzed 36 tracts across these groups, comparing results with and without BundleWarp integrated into BUAN. We conducted four key experiments investigating differences in microstructural measures along the length of 36 white matter tracts between (1) subjects with mild cognitive impairment (MCI) and cognitively healthy control subjects, (2) dementia patients and cognitively healthy controls, (3) amyloid-positive and amyloid-negative subjects, and (4) tau-positive and tau-negative. Each experiment was performed twice: once with BundleWarp nonlinear registration integrated into BUAN and once with SLR affine registration integrated into BUAN. BundleWarp consistently improved the accuracy of tractometric analyses by aligning tracts more precisely across subjects while preserving their anatomical structure. Additionally, it helped mitigate potential artifacts arising from preprocessing and tractography, resulting in more harmonized bundle shapes and enhanced sensitivity to disease-related changes in certain tracts. Notably, the inclusion of BundleWarp in BUAN generated *p*-value curves (representing along-tract statistics) with higher auto-correlation, indicating improved segment mapping and consistency in alignment. These results highlight BundleWarp’s role in advancing the precision and robustness of tractometry for detecting subtle white matter changes in neurodegenerative diseases.

To further evaluate BundleWarp’s ability to enhance subject-level reproducibility, we conducted a test–retest fingerprinting analysis using 44 subjects from the Human Connectome Project (HCP) dataset, each scanned twice. By comparing bundle shape similarity scores and along-tract microstructural profiles (FA, MD, RD, AD) across the two sessions, we found that BundleWarp substantially increased within-subject correspondence relative to SLR. Paired t-tests on subject-level shape similarity scores showed significant improvements across the majority of bundles, and test–retest correlations of microstructural profiles were consistently higher with BundleWarp. These results demonstrate that BundleWarp enhances subject fingerprinting, yielding more stable and reproducible representations of white matter tracts across repeated scans.

Additionally, we demonstrate how the displacement field generated by BundleWarp enables tract morphometry by visually and analytically quantifying shape differences between white matter bundles. This approach provides a detailed assessment of structural changes, offering insights into localized deformations and alterations in bundle geometry, which are critical for understanding disease-related white matter disruptions. We apply this morphometry framework to the ADNI3 dataset, evaluating bundle-shape differences between amyloid-positive vs. amyloid-negative and tau-positive vs. tau-negative groups. These experiments show that BundleWarp’s shape-based metrics are sensitive enough to detect subtle, spatially localized morphological deviations associated with early Alzheimer’s pathology, highlighting the utility of BundleWarp for clinical and preclinical investigations.

## 2. Methods

### 2.1. Data

We use 730 ADNI3 participants (phase 3 of the Alzheimer’s Disease Neuroimaging Initiative; age range: 55-95 years, 349M/381F, 214 with MCI, 69 with AD, and 447 cognitively healthy controls (CN)) scanned with 7 acquisition protocols (GE36, GE54, P33, P36, S127, S31, S55) were included. Tables 1 and 2 in Fig. 1 detail demographic and acquisition protocol information. More information on the MRI acquisition and processing can be found online at https://adni.loni.usc.edu/adni-3/. A*β*-status, i.e., positive (A*β*+) or negative (A*β*–), was determined by either mean 18F-florbetapir (A*β*+ defined as >1.11) Landau, Breault, Joshi, Pontecorvo, Mathis, Jagust and Mintun (2013a); Landau, Thomas, Thurfjell, Schmidt, Margolin, Mintun, Pontecorvo, Baker, Jagusst and Initiative (2014) or florbetaben (A*β*+ defined as >1.20)Landau, Mintun, Joshi, Koeppe, Petersen, Aisen, Weiner, Jagust and Initiative (2012); Landau, Lu, Joshi, Pontecorvo, Mintun, Trojanowski, Shaw, Jagust and Initiative (2013b). PET cortical SUVR uptake, normalized by using a whole cerebellum reference region. Tau positivity was defined as a tau SUVR > 1.23.

**Table 1.** Summary of BundleWarp (BW) improvements across diagnostic and biomarker groups. Each entry shows the number (and percentage) of bundles (out of 36) with higher mean − log_10_(*P*-values) (statistical significance) and higher autocorrelation (smoothness) compared to Streamline-based Linear Registration (SLR).

| Metric | MCI |  | Dementia |  |
| --- | --- | --- | --- | --- |
|  | ↑ Significance | ↑ Smoothness | ↑ Significance | ↑ Smoothness |
| AD | 22/36 (61.1%) | 31/36 (86.1%) | 22/36 (61.1%) | 27/36 (75.0%) |
| FA | 24/36 (66.7%) | 33/36 (91.7%) | 24/36 (66.7%) | 26/36 (72.2%) |
| MD | 25/36 (69.4%) | 27/36 (75.0%) | 25/36 (69.4%) | 25/36 (69.4%) |
| RD | 29/36 (80.6%) | 31/36 (86.1%) | 29/36 (80.6%) | 26/36 (72.2%) |

| Metric | Amyloid+ |  | TAU+ |  |
| --- | --- | --- | --- | --- |
|  | ↑ Significance | ↑ Smoothness | ↑ Significance | ↑ Smoothness |
| AD | 22/36 (61.1%) | 24/36 (66.7%) | 22/36 (61.1%) | 27/36 (75.0%) |
| FA | 24/36 (66.7%) | 26/36 (72.2%) | 24/36 (66.7%) | 25/36 (69.4%) |
| MD | 25/36 (69.4%) | 23/36 (63.9%) | 25/36 (69.4%) | 26/36 (72.2%) |
| RD | 29/36 (80.6%) | 22/36 (61.1%) | 29/36 (80.6%) | 22/36 (61.1%) |

**Figure 1.**
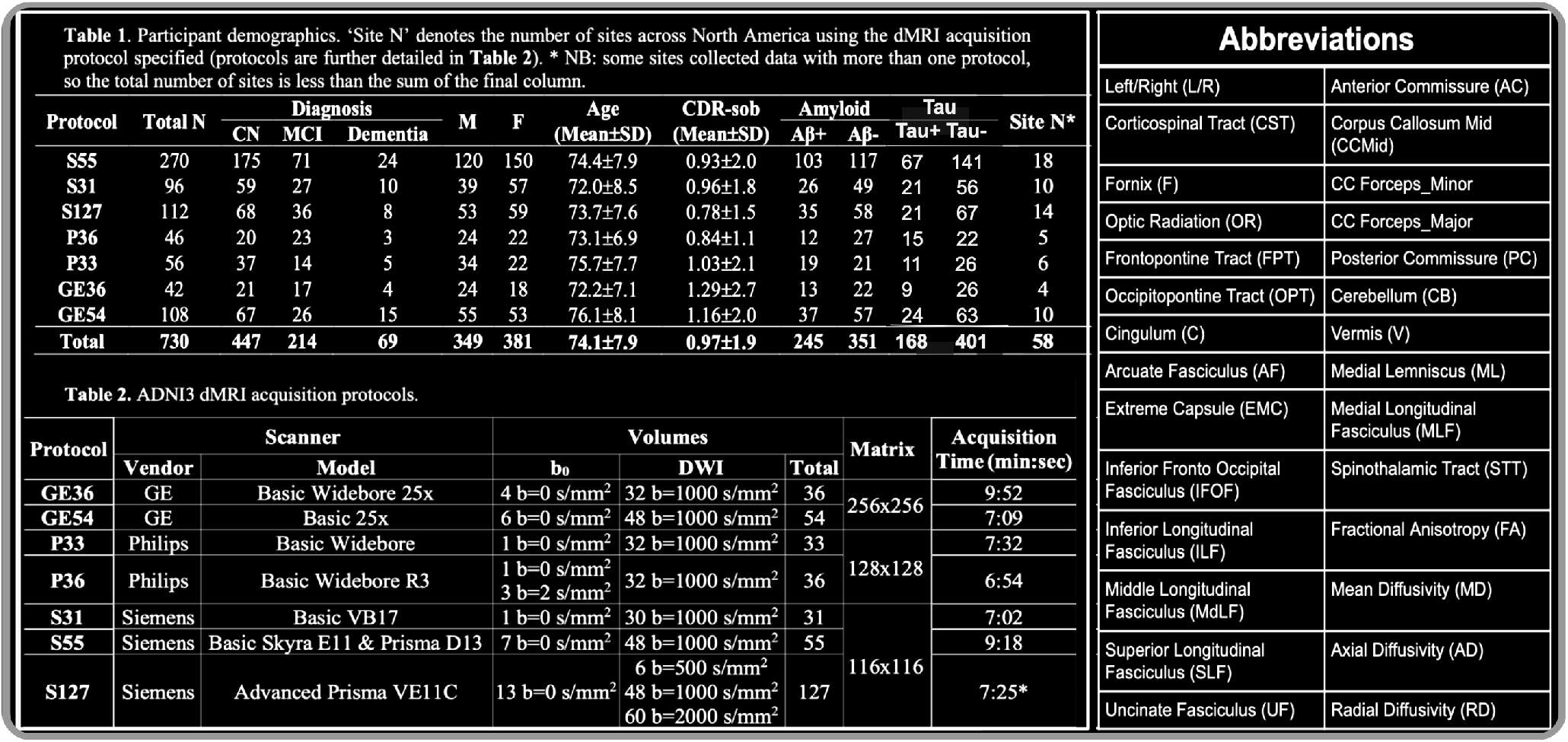
Tables 1 and 2 detail demographic and scanner protocol information for the ADNI3 data used in our experiments. The abbreviation table on the right lists the 36 white matter tracts and four microstructural measures analyzed in this work.

Additionally, we used a subset of 64 subjects obtained from the Parkinson’s Progression Markers Initiative (PPMI) Marek et al. (2011) database. PPMI diffusion MRI data were acquired using a standardized protocol used on Siemens TIM Trio and Siemens Verio 3 Tesla MRI machines from 32 different international sites. Diffusion-weighted images were acquired along 64 uniformly distributed directions using a b-value of 1000 *s*/*mm*^2^ and a single b=0 image. More information on the MRI acquisition and processing can be found online at www.ppmi-info.org. We have processed 64 subjects: 32 controls and 32. We selected subjects in the age range of 39-61. Both groups have 16 females and 18 males.

For the test–retest reproducibility analysis, we used the Human Connectome Project (HCP) test–retest dataset Sotiropoulos, Jbabdi, Xu, Andersson, Moeller, Auerbach, Glasser, Hernandez, Sapiro, Jenkinson et al. (2013) that included 44 healthy young adults (ages 22–35 years), each scanned twice on a 3 Tesla MRI scanner with an average inter-scan interval of 4.7 / 2 months (range: 1–11 months). Diffusion MRI was acquired with a high-angular-resolution protocol consisting of 18 b0 images and 90 gradient directions at b = 3000 *s*/*mm*^2^, with 1.25 mm isotropic voxel resolution (TE/TR = 89/5520 ms).

### 2.2. Data Preparation

Raw ADNI3 diffusion MRI (dMRI) were preprocessed using the ADNI3 dMRI protocol Jahanshad, Kochunov, Sprooten, Mandl, Nichols, Almasy, Blangero, Brouwer, Curran, de Zubicaray et al. (2013); Thomopoulos, Nir, Reina, Jahanshad and Thompson (2020). Preprocessing of data involved several steps: denoising raw dMRI data using DIPY’s principal component analysis (PCA) for GE data, and Marchenko-Pastur PCA for Siemens and Philips data denoising Manjón, Coupé, Concha, Buades, Collins and Robles (2013); Garyfallidis, Brett, Amirbekian, Rokem, Van Der Walt, Descoteaux, Nimmo-Smith and Contributors (2014a). Gibbs artifacts were corrected using MRtrix’s *degibbs* tool Kellner, Dhital, Kiselev and Reisert (2016); Tournier, Smith, Raffelt, Tabbara, Dhollander, Pietsch, Christiaens, Jeurissen, Yeh and Connelly (2019), and extracerebral tissue was removed (skull stripping) with FSL’s BET Smith (2002); Jenkinson, Beckmann, Behrens, Woolrich and Smith (2012). Eddy currents and motion were corrected using FSL’s eddy_cuda tool with additional corrections for slice-to-volume and outlier detectionAndersson and Sotiropoulos (2016); Jenkinson et al. (2012). Bias field inhomogeneities were corrected using MRtrix’s *dwibiascorrection* ANTS function. Preprocessed T1w images from the ADNI database were further processed and aligned to the dMRI data Tustison, Avants, Cook, Zheng, Egan, Yushkevich and Gee (2010); Tournier et al. (2019). ADNI3 dMRI data lacked reversed phase-encode blips, so echo-planar imaging (EPI) distortion corrections were made using nonlinear registrations to T1-weighted anatomical images. The processed dMRI data were converted back to native space through a series of inversions of the registration matrices, with final outputs visually inspected and manually adjusted as necessary. The diffusion tensor imaging (DTI) model was used to extract 4 microstructural measures from processed dMRI: fractional anisotropy (FA), mean, axial, and radial diffusivity (MD, AD, RD).

We applied a robust and unbiased model-based spherical deconvolution Canales-Rodríguez, Daducci, Sotiropoulos, Caruyer, Aja-Fernández, Radua, Yurramendi Mendizabal, Iturria-Medina, Melie-García, Alemán-Gómez et al. (2015) reconstruction method and a probabilistic particle filtering tracking algorithm that uses tissue partial volume estimation (PVE) to reconstruct Girard, Whittingstall, Deriche and Descoteaux (2014) whole-brain tractograms. For tracking, the seed mask was created from the white matter (WM) PVE (WM PVE > 0.5), seed density per voxel was set to 2, and step size was set to 0.5. We extracted thirty-six white matter (WM) tracts from tractograms using auto-calibrated RecoBundles Garyfallidis, Côté, Rheault, Sidhu, Hau, Petit, Fortin, Cunanne and Descoteaux (2018); Chandio et al. (2020) (see Fig. 1 for full names) using model bundles from the HCP-842 tractography atlasYeh, Panesar, Fernandes, Meola, Yoshino, Fernandez-Miranda, Vettel and Verstynen (2018b).

PPMI Diffusion MRI data was denoised using the local principal component analysis (LPCA) noise reduction method Manjón et al. (2013). For brain tissue extraction, the median Otsu algorithm Yang, Shen, Long and Chen (2012) was used. The distortions induced by eddy currents and motion were corrected by registering the diffusion-weighted volumes to the b0 volume. An affine transformation was computed to register b0 and non-b0 3D volumes by maximizing normalized mutual information. The optimization strategy used in DIPY is similar to that implemented in ANTs Avants et al. (2009). The B-matrix (b-vectors) were rotated to preserve the correct orientational information as described in this paper by Leemans and Jones (2009). The whole-brain tractograms were generated by fitting a constrained spherical deconvolution (CSD) model to get directional information from dMRI data Tournier, Calamante and Connelly (2007), from which a simplified peaks representation was extracted. The obtained peaks were then used as input for a local tracking algorithm. We performed deterministic tracking Mori, Crain, Chacko and Van Zijl (1999); Basser, Pajevic, Pierpaoli, Duda and Aldroubi (2000) using EuDX Garyfallidis (2012). The EuDX tracking algorithm was initialized with the following parameters: tracking starts from voxels where fractional anisotropy (FA) > 0.3, number of seeds per voxel = 15, step size = 0.5, angular threshold = 60 degrees, and stops tracking if the FA value drops below 0.1. Each generated tractogram comprised 5-10 million streamlines. Twenty-seven white matter tracts were extracted from each subject’s whole brain tractogram by deploying auto-calibrated RecoBundles Garyfallidis, Côté, Rheault, Sidhu, Hau, Petit, Fortin, Cunanne and Descoteaux (2017); Chandio et al. (2020).

We used 44 subjects from the HCP test–retest dataset, each scanned twice. The preprocessed diffusion MRI data provided by HCP were used for tractography. The DTI model was fit to extract microstructural metrics using DIPY Garyfallidis, Brett, Amirbekian, Rokem, van der Walt, Descoteaux, Nimmo-Smith and Dipy Contributors (2014b). Whole-brain tractograms were generated using multi-shell multi-tissue constrained spherical deconvolution (MSMT-CSD) Jeurissen, Tournier, Dhollander, Connelly and Sijbers (2014) and deterministic local tracking. Tracking was initiated from voxels with fractional anisotropy (FA) > 0.15, using 8 seeds per voxel, a step size of 0.5 mm, and an angular threshold of 60°. Tracking was terminated when FA dropped below 0.15. Thirty-six white matter tracts were then extracted from each subject’s whole-brain tractogram using the auto-calibrated RecoBundles algorithm.

For each of the three datasets used in this paper, auto-calibrated RecoBundles Garyfallidis et al. (2018); Chandio et al. (2020), leveraging model bundles from the HCP-842 tractography atlas Yeh et al. (2018b) was used for bundle segmentation. We used the default parameters of the auto-calibrated RecoBundles: reduction_thr = 15.0, pruning_thr = 8.0, and pruning_distance = “mdf”, followed by a second round of refinement using r_reduction_thr = 12.0 and r_pruning_thr = 6.0. The initial reduction step decreases the search space by creating a local neighborhood around the model bundle, facilitating more efficient streamline matching. The subsequent pruning step removes streamlines whose distance from the model bundle exceeds the specified threshold. The second-round parameters (r_reduction_thr and r_pruning_thr) provide additional refinement of the segmented bundle, as proposed in the BUAN pipeline Chandio et al. (2020), to further eliminate outliers and ensure anatomically coherent bundle representations.

### 2.3. Tractography Data Representation

This section explains the streamline data structure to help readers better understand the methods presented in the following sections.

A streamline is stored as an ordered sequence of 3D vector points. The collection of streamlines is called a tractogram/tract. Let *S*_*i*_ be a streamline in a whole-brain tractogram/tract *T*.

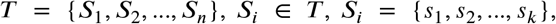

where *n* is the total number of streamlines in *T* and *s*_*i*_ denotes a 3D vector point. Streamlines in *T* can have different numbers of points (*k*) and different lengths. Here, the length of a streamline implies its Euclidean length in millimeters. Since each streamline is a set of 3D vectors, it can be considered a point-set.

### 2.4. BundleWarp Registration

Given two bundles, a static bundle (often referred to as fixed in most registration methods) and a moving bundle (bundle to be registered), BundleWarp performs a two-step streamline-based registration to align the moving bundle with the static bundle.

In the first part, we linearly align two bundles using SLR Garyfallidis et al. (2015).

The SLR can be summarized in the following steps:

1. Set all streamlines from both static and moving bundles to have the same number of points K per streamline
2. Minimize BMD cost function
3. Apply the transformation to the moving bundle to align it with the static bundle.

Where BMD stands for bundle-based minimum distance Garyfallidis et al. (2015) described in Supplement S.1.2.

After streamlines are roughly aligned and are in the same space, we start the deformation process. BundleWarp method is visually explained in Fig. 2 and Fig. 3. In the deformable registration, every streamline and every point on a streamline gets a nonlinear transformation map (warp map) specific to it, unlike linear registration, where all streamlines share one global transformation map. We create streamline correspondences between streamlines in moving and static bundles so that we can warp one streamline from moving to one streamline in static. However, static and moving bundles often do not have the same number of streamlines, and one-to-one mapping would not suffice in that case. Therefore, we design a many-to-one pairwise streamline correspondence approach based on the rectangular Linear Assignment Problem (rLAP) Olivetti et al. (2020); Kuhn (1955) for bundles with a varying number of streamlines that we call iterLAP.

**Figure 2.**
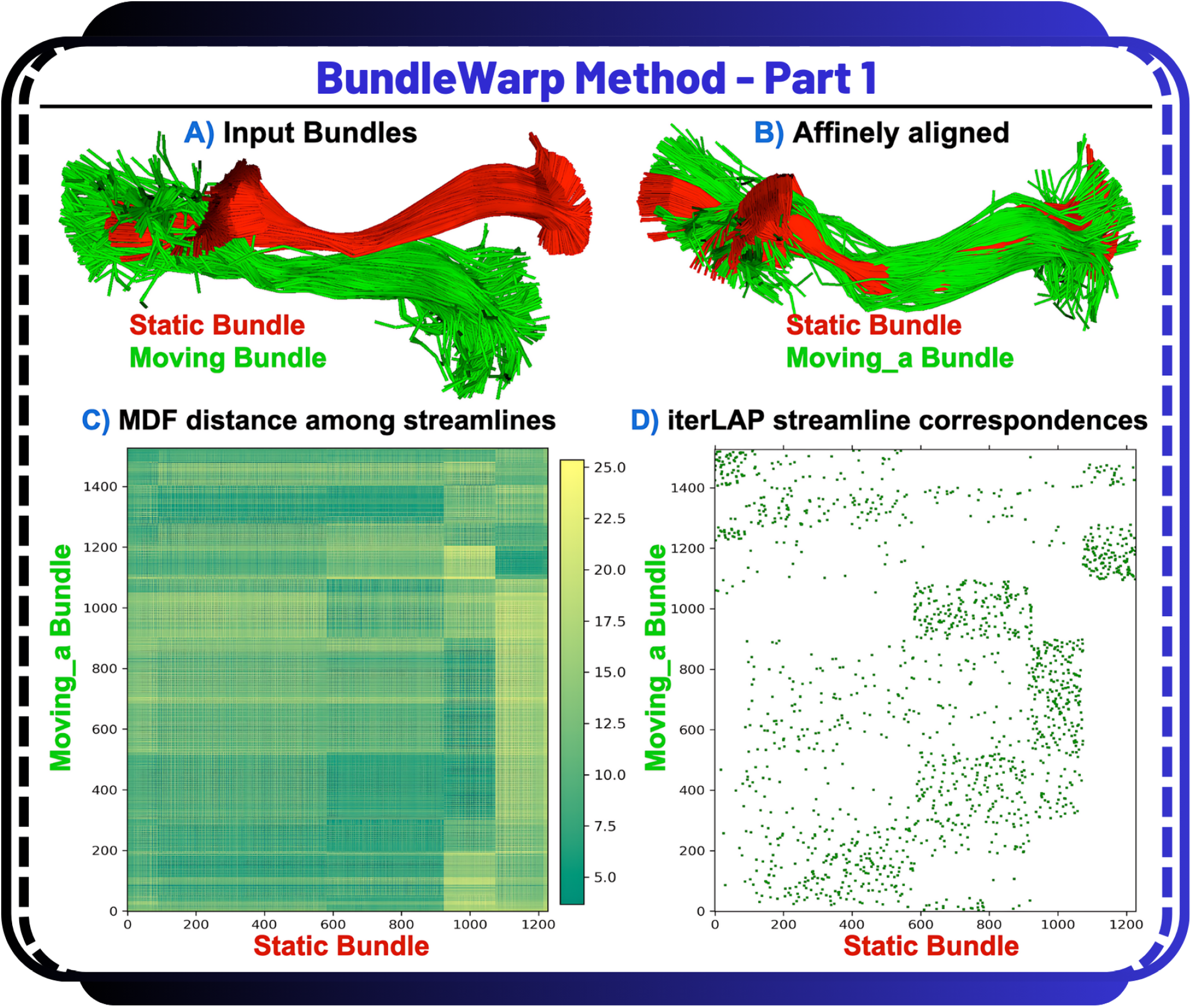
Steps A-D in BundleWarp (BW) method. A) Input static (red) and moving (green) IFOF_L bundles. B) The moving bundle is affinely aligned with the static bundle. C) MDF distance is calculated among streamlines of the affinely moved (moving_a) bundle and the static bundle. D) Using the distance matrix from (C), the many-to-one rectangular linear assignment problem (iterLAP) is applied to find streamline correspondence in two bundles. In (D), green pixels show correspondence between the x-axis (static bundle’s streamlines) and y-axis (moving_a bundle’s streamlines). The pixel size has been enlarged for visibility. See Fig 3 for the next steps in BW.

**Figure 3.**
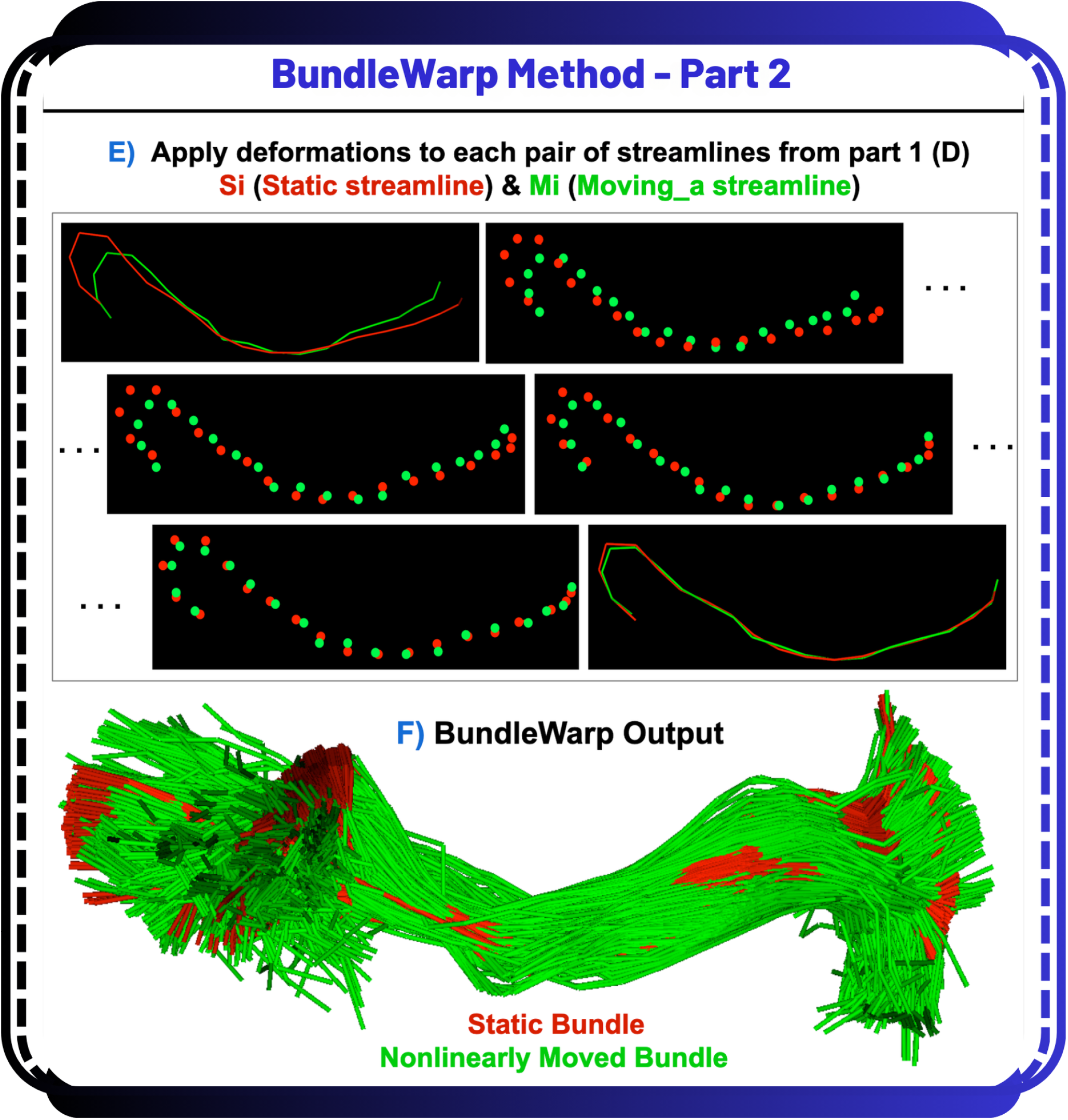
Steps E and F in BundleWarp method. Iterate over matched pairs of streamlines from Fig. 2.D and deform moving_a streamline. (E) Streamlines are considered point-set clouds, and each streamline is deformed independently. (F) The final output of the BundleWarp method with *λ*=0.5.

The linear assignment problem (LAP)Kuhn (1955) is also known as minimum weight matching in bipartite graphs. The problem is described by a cost matrix C, where each *C*[*i, j*] element is the cost of matching a vertex (streamline) *i* of the first partite set (moving bundle) with a vertex (streamline) *j* of the second set (static bundle). The goal is to find a complete assignment of moving bundle’s streamlines to static bundle’s streamlines that results in minimal cost. Where LAP can be written as

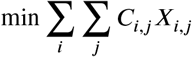

Here, *X* is a boolean matrix, and if *X*[*i, j*] is equal to 1, then the moving bundle’s *i*^*th*^ streamline is assigned/matched to *j*^*th*^ streamline of the static bundle. In classic LAP, the cost matrix *C* is a square matrix. The method can be extended to rectangular LAP (rLAP) to have a rectangular cost function when two sets do not have the same number of elements. However, both LAP and rLAP result in one-to-one matching/assignment. In rLAP, when there are more rows than columns in *C*_*m*×*n*_, it only assigns *n* rows to *n* columns.

We design a many-to-one rectangular LAP (iterLAP) to obtain a many-to-one correspondence of streamlines when *m* > *n*.

We begin by computing the Minimum-average Direct Flip (MDF) distance Garyfallidis, Brett, Correia, Williams and Nimmo-Smith (2012) between each streamline in the moving bundle and each streamline in the static bundle. This results in an *m* × *n* distance matrix, which serves as the cost matrix *C* for subsequent assignment (Fig. 2C).

Briefly, the MDF distance between two streamlines *s* and *t*, each represented by *K* corresponding points, is defined as:

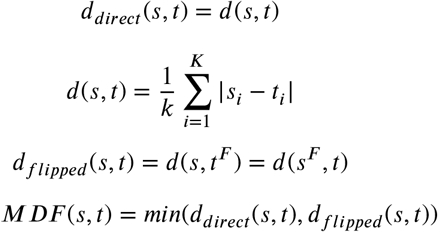

Next, we feed this *m* × *n* MDF distance matrix to rLAP. rLAP returns *n* matched pairs. We are left with *m* − *n* streamlines in the moving bundle and need to be matched with streamlines in the static. We run rLAP again on the sub-matrix *Ĉ* of MDF distance matrix *C*. Where, *Ĉ* is a (*m* − *n*) × *n*. We rerun rLAP using *Ĉ* as a cost matrix. If (*m* − *n*) > *n* and still some moving bundle’s streamlines are unmatched, we repeat the process until all streamlines have found a corresponding streamline in the static bundle. In the case of *m* < *n*, the remaining unmatched streamlines in the static bundle are discarded since all streamlines in the moving bundle have found corresponding streamlines in a static bundle.

iterLAP can be summarized as follows:

1. Calculate *m* × *n* MDF distance matrix *C* from moving and static bundles’ streamlines
2. While all streamlines in moving are not matched
  a. Run rLAP on *C* to get pairwise streamline correspondences
  b. *k* = *m* − *n*
  c. if *k* < *n* :
    i. Ĉ = *k* × *n* sub-matrix of *C Ĉ* has the distance between unmatched k streamlines in moving bundle and all n streamlines in static bundle
    ii. *C= Ĉ*
3. *matched*_ *pairs* is a list of elements. Each element is a tuple (*i, j*). Where *i* is the moving bundle’s streamline index, and *j* is the index of its corresponding streamline in a static bundle.

In the second part, after we have built the correspondences among streamlines of moving and static bundles, we start to warp them (find nonlinear transformations) using the proposed memoryless Coherent Point Drift (mlCPD) registration method for a bundle of streamlines. In mlCPD, for each pair of corresponding streamlines in *matched*_*pairs*, we perform deformable point-cloud registration using the Coherent Point Drift (CPD) methodMyronenko et al. (2006); Myronenko and Song (2010). Since a single streamline is an ordered set of points, it can be treated as a point-cloud set (Fig. 3E). Here, we iterate and apply CPD registration separately on each pair of streamlines. Through iteratively applying CPD, (1) we move the points on the same streamline together, taking into account the local neighborhood information/connectivity, this step helps produce smooth transformations and (2) we save memory and computational power since we work with small point-clouds of two streamlines instead of entire bundles. Dealing with a large number of point-clouds could become a computationally extensive task.

If the whole bundle is considered one large point-cloud representing the entire bundle, registration would lose the streamline connectivity information. For instance, consider two points along the same streamline in motion; they could be registered to two distinct streamlines by aligning one point with one streamline and the second point with another. This process might inadvertently create spurious streamlines when converting the point cloud back into a bundle. Moreover, the original CPD method, if applied to the entire bundles, would fail due to large memory allocation requests. Hence, we propose mlCPD to overcome the above mentioned limitations. mlCPD does require the initial matching of pair streamlines by iterLAP (see above). iterLAP is calculated once, and mlCPD is calculated as many times as the number of streamlines in the moving bundle.

The mlCPD algorithm takes a probabilistic approach to align two point-sets. Registration is formulated as a maximum likelihood (ML) estimation problem using the Gaussian Mixture Models (GMMs) method. The points in the moving set *Y* are considered as the GMM centroids, while the static point-set *X* represents the data points generated by the GMM. The GMM centroids from the *Y* point-set are fitted to the *X* point-set by maximizing the likelihood. When the two point-sets are optimally aligned, the correspondence is the maximum of the GMM posterior probability for a given data point. To preserve the topological structure of the point-sets (streamlines), the GMM centroids are forced to move coherently as a group. The expectation-maximization (EM) algorithm is used to optimize the cost function. It concurrently finds the nonrigid transformation and the correspondence amongst point-sets of two streamlines. It regularizes the velocity field such that points move coherently using motion coherence theoryYuille and Grzywacz (1988, 1989).

Here, *Y* = (*y*_1_, *y*_2_, …, *y*_*m*_)^*T*^ is a moving streamline represented as a *m* × 3 matrix. *m* is the number of points in the moving streamline. *X* = (*x*_1_, *x*_2_, …, *x*_*n*_)^*T*^ is a static streamline represented as a *n* × 3 matrix. *n* is the number of points in the static streamline. Points in *Y* are considered as the centroids of a GMM. Note that the number of points on streamlines of two bundles can vary. Points in *Y* are fitted to the data points *X* by maximizing the likelihood function. The GMM probability density function is written as

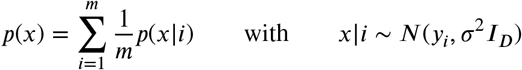

Where *D* is the dimensions of the centroids (*D* = 3 in our case).

Let *Y*_*o*_ be the initial centroid values, *G*_*m*×*m*_ a square symmetric Gram matrix with elements

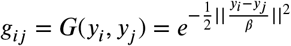

*W*_*m*×*m*_ = (*w*_1_, *w*_2_, …, *w*_*m*_)^*T*^ is a Gaussian kernel weights, *I* is identity matrix, and 1 is a column vector of all ones. The estimated velocity field *v* represents the underlying non-rigid transformation. For free parameters, *λ* represents the trade-off between data fitting and smoothness regularization, and *β* represents the strength of the interaction between points. *σ* captures the range for each Gaussian mixture component. *α* is the annealing rate. In the M-step of the EM algorithm, a system of nonlinear equations is solved for *W*. mlCPD algorithm for streamlines can be summarized as follows.

For each pair of streamlines in *matched*_*pairs*:

1. Initialize parameters *λ* > 0, *β* > 0, and *σ*.
2. Construct G matrix, initialize *Y* = *Y*_*o*_. Where *Y*_*o*_ is a moving streamline, and *X* is a static streamline.
3. Deterministic annealing:
  - EM optimization:
    - E-step: Compute posterior probabilities matrix *P*
    - M-step: solve for W:

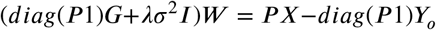
    - update *Y* = *Y*_*o*_ + *GW*
  - Anneal *σ* = *ασ*
4. Compute the velocity field *v*(*z*) = *G*(*z*,.)*W*
5. Get correspondence from posterior probabilities *P*

Where, *P*_*m*×*n*_ is a posterior probability matrix of GMM with:

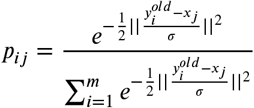

Regularization is imposed using Gaussian kernel, which takes a similar regularization form as the one in motion coherence theoryMyronenko and Song (2010); Yuille and Grzywacz (1988, 1989). This way, we regularize the deformation field to make it smooth and coherently move points on the same streamline to preserve its topology, preventing the creation of spurious streamlines.

BundleWarp method’s registration iterations are visualized in Fig. 4. Here, iteration 0 implies linear registration that could be affine or rigid (we keep affine by default). In the rest of the iterations, the moving bundle is deformed. BundleWarp converges within 15 iterations. For the purpose of showing how quickly and accurately BundleWarp converges and *fully* deforms the moving bundle, we selected the regularization parameter *λ* = 0.001. However, the level of deformations can be controlled by increasing the value of *λ*. See section Parameter Selection and Discussion section for detailed discussion on the selection of *λ*.

**Figure 4.**
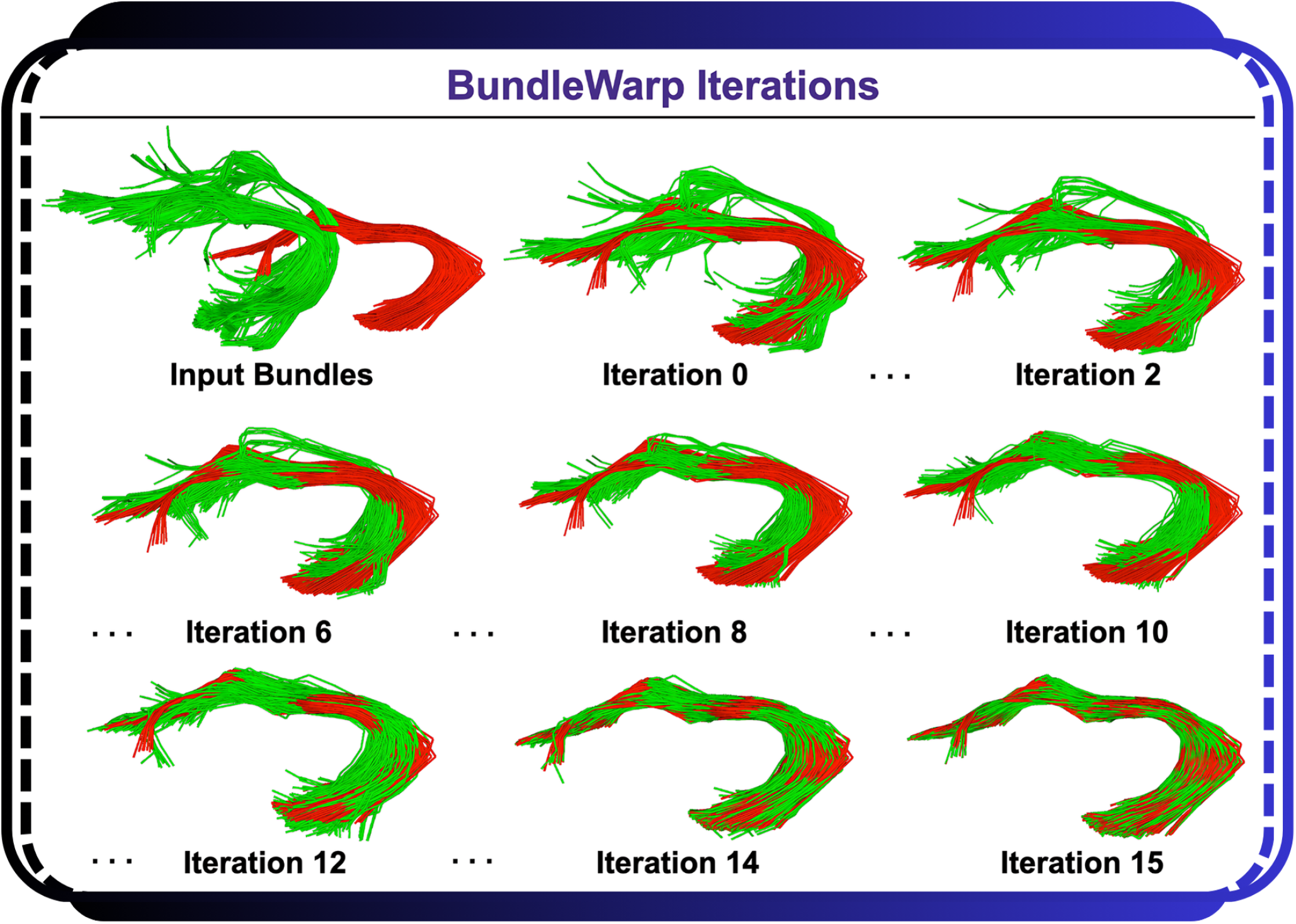
BundleWarp (BW) method iterations. Red is the static, and green is the moving bundle. Iteration 0 means affine registration. The rest of the iterations show a deformable registration process in BW. BW converges in 15 iterations.

### 2.5. Parameter Selection

The optimal expected output of nonlinear tract registration is non-trivial to decide. The same type of white matter bundle can have different shapes and sizes for different subjects. Deforming the original topological/anatomical shape of the bundle is not necessarily ideal, as it can lose the characteristics needed in the analysis. For example, a patient with a tumor will have some pathways missing from the area of the tumor location. Deforming the shape of the bundle too much, in this case, can lead to the nonrecognition of abnormalities/tumors in the analysis.

BundleWarp provides users with an option to control the level of deformations needed in the output. Users can control deformations by providing the values of *λ*, where *λ* controls the trade-off between regularizing the deformation and having points match very closely. The lower the value of *λ*, the more closely the bundles would match. Another parameter *β* controls the smoothness of the deformation. *β* represents the width of the smoothing Gaussian filter. By default, we set *β* to be 20 for all bundles except when the Euclidean length of the bundle is shorter than 50 mm. In the case of small bundles with length < 50mm, *β* is set to 10. A low value of *β* contributes to generating locally smooth transformations. A higher value of *β* will make the model produce more of a translation transformation. While regularization weights are well-established in registration frameworks such as ANTs SyN, to the best of our knowledge, BundleWarp is the first to offer an easy-to-use, tunable deformation parameter specifically designed for streamline-based bundle registration. This parameter allows users to directly control the degree of nonrigidity, enabling either *partial* or *full* deformation of a bundle within a tractography-focused registration framework.

As *λ* regularizes the velocity field. Setting *λ* to a relatively higher value, such as 0.5, we include fewer deformations in the model, and the moving bundle is *partially* deformed. The output will be an improvement over linear registration, yet the bundle will retain its original topological/anatomical shape, as shown in Fig. 9.

In Fig. 5, we explore different values of *λ* on the Corticospinal tract in the left hemisphere (CST_L). Here, (a) is the moving bundle (to be registered), and (b) is the static bundle. As we decrease the value of *λ* (c-f), we see the moving bundle is deformed more by the BundleWarp method and starts to look more like the static bundle. A lower value of *λ* will yield results such that the moving bundle will be completely deformed to look exactly like the static bundle, as seen in Fig: 5f. For all registrations, we kept the default value of *β* = 20 for spatially smooth transformation. Please note that *fully* deformable registration (f) is not recommended. However, in this demonstration, we showcase it to illustrate the capabilities of BundleWarp registration.

**Figure 5.**
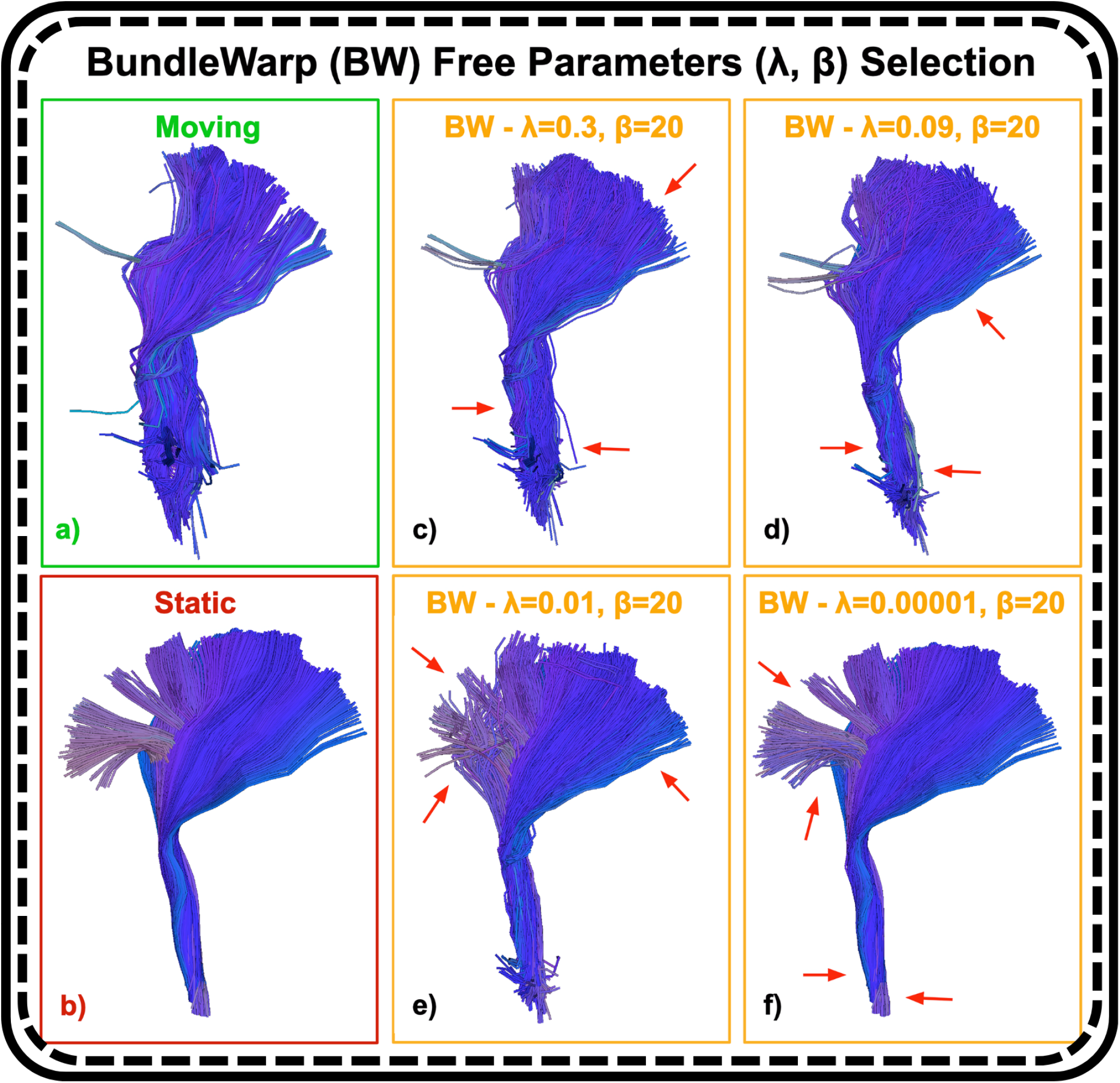
BundleWarp free parameter selection. Here, (a) is the moving bundle, and (b) is the static bundle. We show the warped bundle (output of BundleWarp) with different values of *λ. λ* represents the trade-off between the goodness of maximum likelihood fit and regularization. A higher value of *λ* penalizes the velocity field more, and the moving bundle is *partially* deformed (c-d). A smaller value of *λ* will force streamlines to *fully* deform and look exactly like the static bundle (e-f). As we decrease the value of *λ* (e-f), we see the moving bundle is deformed more and starts to look more like the static bundle. Here, we keep *β* = 20 for all registrations (c-f). However, this parameter can be easily adjusted in BundleWarp.

### 2.6. Robust Tractometry with BundleWarp

We show an application of BundleWarp registration in the tractometry pipeline. Adding BundleWarp into BUndle ANalytics (BUAN) tractometryChandio et al. (2020) pipeline improves the robustness of statistical analysis and increases the effect sizes.

Fig.6 visualizes steps in BUAN tractometry with BundleWarp integrated. BUAN extracts bundles from populations and analyzes the microstructural measures (e.g., FA) projected onto the bundles along the length of the tracts to find significant group differences. BUAN creates N horizontal segments along the length of the tracts to be analyzed. Horizontal segments are created based on points on the streamlines of a bundle belonging to the closest model bundle centroid point. Here, we add a *partially* deformable BundleWarp registration to nonlinearly align subjects’ bundles with model bundles before creating the horizontal segments to find better segment correspondences among populations. Note that the nonlinearly moved bundles are only used to assign segment numbers to streamlines (and points on the streamlines) in the bundles. Actual statistical analysis always takes place in the native space of diffusion data. The statistical analysis step uses bundles of the original shape and microstructural measures in the native space utilizing segment labels given during the assignment step for segment-specific group analysis.

**Figure 6:**
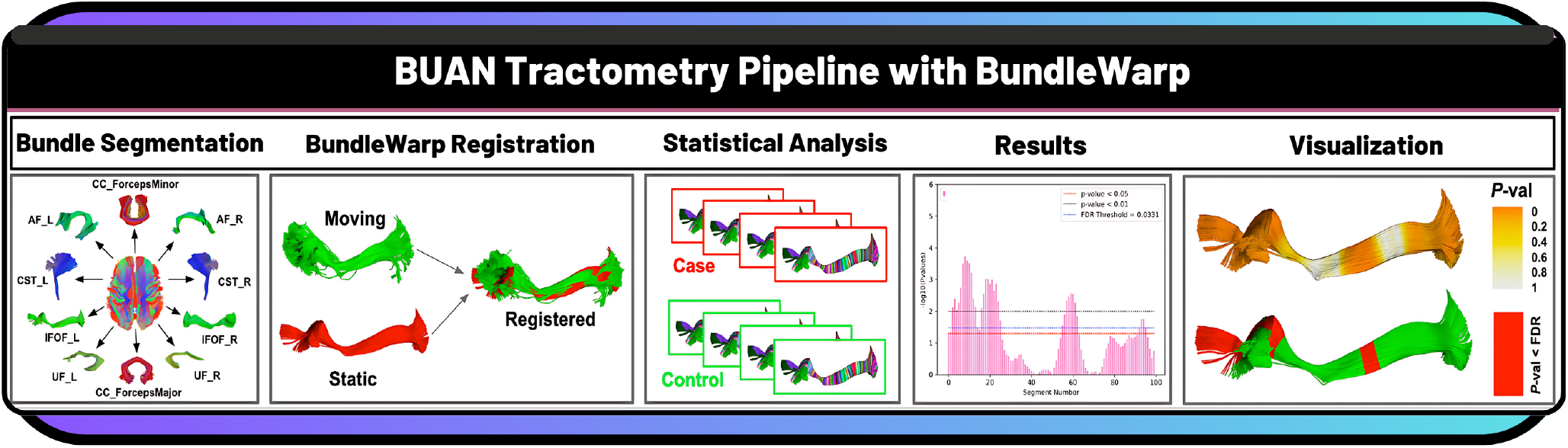
BUAN tractometry with BundleWarp registration: The pipeline starts with bundle segmentation from whole brain tractograms. Extracted bundles are then nonlinearly registered to atlas bundles using BundleWarp. N segments are created along the length of the bundles. Linear mixed models are run per segment along the length of the tracts. Results can be plotted as a graph or as *p*-values projected along the length of the tracts.

### 2.7. Quantifying Bundle Shape Differences (Tract Morphometry)

Apart from registering/aligning two bundles, the displacement vector field derived from *fully* deformable BundleWarp registration can be used to quantify bundle shape differences. We propose bundle shape differences analysis using the BundleWarp method when *λ* is set to a low value such as *λ* = 0.00001. The overview of the proposed approach is visualized in Fig. 7. In Fig. 7, static (red) and moving (blue) corticospinal tracts in the left hemisphere of the brain (CST_L) are visualized before registration is applied to them (A). Static (B) and moving (C) bundles are visualized individually. *Fully* deformable BundleWarp registration is applied with *λ* = 0.00001 and

**Figure 7:**
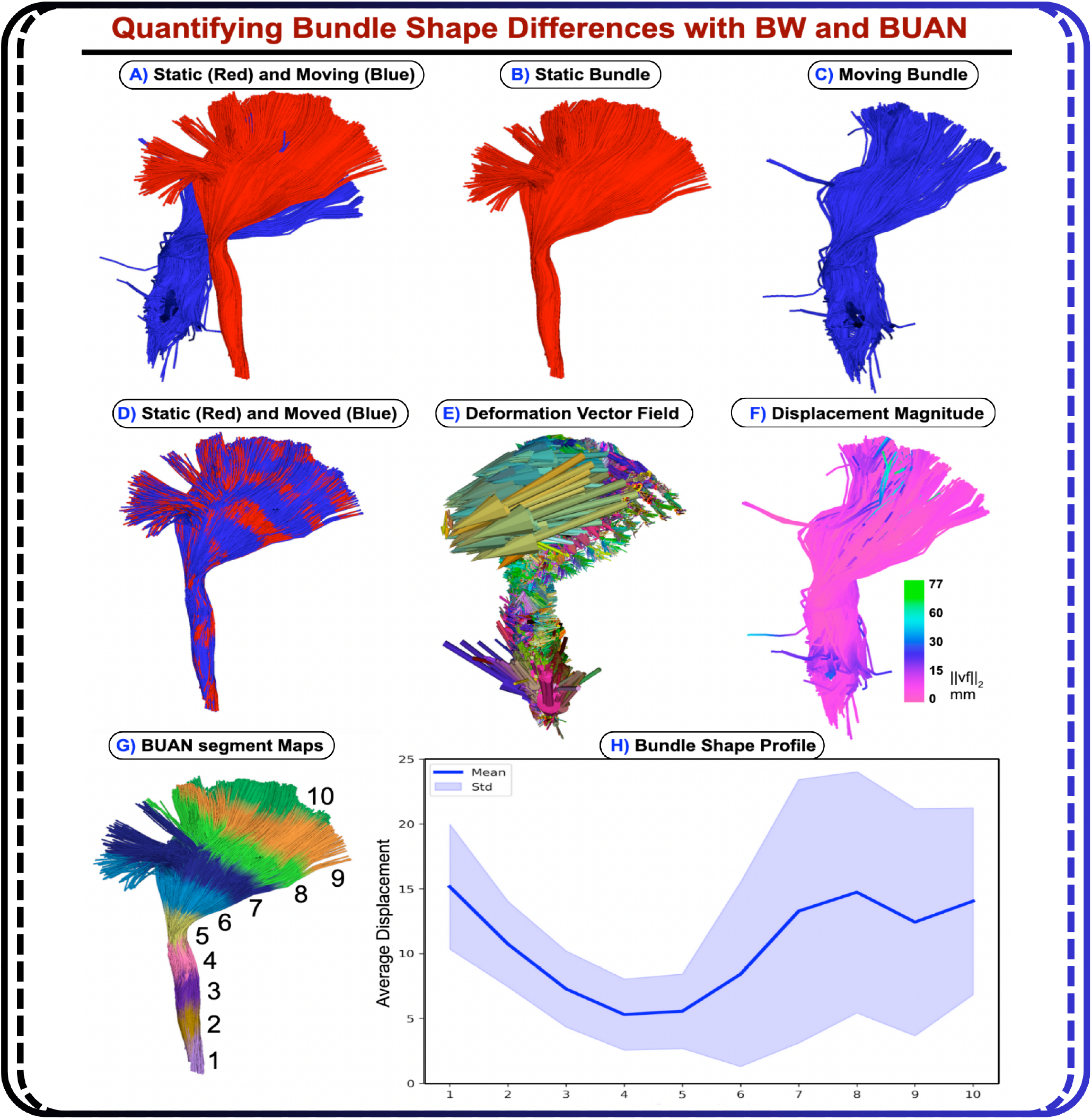
Quantifying bundle shape differences using the displacement field generated by BundleWarp (BW) and projecting it along the length of the bundle with BUAN. A) shows input static and moving bundles overlapped before registration. B) static bundle, C) moving bundle. D) shows static and moved (deformed) bundle overlapping after BW registration is applied. E) shows the deformation vector field generated by BW. F) The displacement magnitude of the vector field has been visualized over the moved bundle. G) 10 segment maps created on moved bundle using BUAN. H) Mean displacement in mm is quantified along the length of the bundle with BUAN tractometry.

*β* = 20. The output of the BundleWarp registration is shown in (D), where static (red) and moved (registered moving bundle) bundles are seen completely overlapping (showing perfect *fully* deformable registration).

When the moving bundle is completely deformed (D), one can compute the displacement vector field *vf* as the difference between the streamline points on the affinely moved bundle (*ap*) and the corresponding points after deformation (*dp*): *vf* = *dp* − *ap*.

This vector field provides information about the quantity and the direction of the deformation applied to every streamline point and can be used to quantify shape differences between two bundles.

The displacement vector field (*vf*) of the *fully* deformed bundle is visualized in Fig. 7E. Here, Fig. 7F. shows the magnitude of the displacement field in mm projected on the deformed bundle. Here, the green color shows areas on the bundle that were deformed the most. Hence the areas with the most shape differences (as they required extreme deformations). We show a novel use of along the length quantification of bundle shape differences with displacement field using BUANChandio et al. (2020) tractometry. Fig. 7G. shows ten segments created along the length of the deformed bundle. The displacement magnitude of all the points belonging to the *i*^*th*^ segment was averaged, and the bundle profile was created as shown in Fig. 7H. Here, the x-axis has the segment number along the length of the CST_L bundle, and the y-axis has the average displacement in millimeters. We can visually (E and F) and analytically (H) quantify from the BundleWarp displacement field that static and moving bundles have the most shape differences at the extremities; specifically the fanning part of the CST_L as those areas require more deformations for two bundles to match completely.

### 2.8. Quantitative Quality Assessment of the Registration

To quantitatively assess the performance of registration, we calculate two streamline-based metrics, bundle-based minimum distance (BMD)Garyfallidis et al. (2015) and bundle shape similarity score (SM)Garyfallidis et al. (2012); Chandio et al. (2020) between static and moved bundles. See Supplement S1 for the mathematical definitions of these metrics. BMD measures the distance between two bundles in millimeters, with a lower BMD indicating that the bundles are closely aligned. SM calculates the overall shape similarity of two bundles once they are in the same space Chandio et al. (2020). SM ranges from 0 to 1, where 0 indicates no similarity and 1 indicates identical shapes. SM requires a threshold to determine how closely we want the bundles to match; here, it was set to 5 mm, requiring the bundles to be very similar in shape to achieve a higher SM value. Additionally, we use the volumetric Dice similarity metric to evaluate the overlap between static and moved bundles. We generate binary fiber density maps for both the static and moved bundles to calculate the Dice similarity, which ranges from 0 to 1. A score of 0 indicates no overlap between the bundles, while a score of 1 indicates complete overlap.

### 2.9. Subject Fingerprinting via Test–Retest Reliability

To evaluate whether BundleWarp improves subject identifiability, we propose a test–retest analysis using extracted tracts from pairs of diffusion MRI scans from the same individuals. Both Streamline-Based Linear Registration (SLR) and BundleWarp nonlinear registration are applied separately to each time point, aligning each subject’s bundles to the corresponding atlas bundles in MNI space. For example, a subject’s left arcuate fasciculus from session 1 and session 2 was independently registered to the left arcuate fasciculus atlas/model bundle.

To assess bundle-shape correspondence, we computed shape similarity scores for each tract, quantifying geometric similarity between corresponding bundles across two sessions on a scale from 0 to 1. For each tract, the time point 1 and time point 2 bundles, after being registered to the atlas bundle using SLR and BundleWarp, were compared to determine their shape similarity. This allowed a direct evaluation of how each registration method affected within-subject bundle alignment across repeated scans. For statistical comparison, we performed a paired *t*-test across subjects for each tract. The input to the test was a pair of vectors containing shape similarity scores (sm) for all subjects (1, 2, … , *n*) for each bundle *i* ∈ {1, 2, … , 36}. Thus, for each bundle *i*, we compared the SLR and BundleWarp (BW) score vectors:

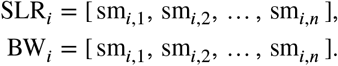

The paired *t*-test evaluated whether the mean shape similarity obtained with BundleWarp was significantly higher than that obtained with SLR for the same subjects. The output of the test included the t-statistic and associated p-value for each tract, indicating whether the improvement in shape correspondence with BundleWarp was statistically significant. We then applied a false-discovery-rate (FDR) correction at *q* < 0.05 across tracts to account for multiple comparisons.

We also evaluated test–retest reproducibility of microstructural bundle profiles, including FA, MD, RD, and AD metrics. In the BUAN tractometry pipeline, subject-specific bundles are aligned to atlas bundles in MNI space using either SLR or BundleWarp, which generates consistent along-tract segment assignments for points on the streamlines. Using these anatomical segment assignments, microstructural values are projected along the length of each bundle in native space. Because tractometry relies on accurate anatomical correspondence across individuals and sessions, registration quality directly influences the consistency of these profiles. To quantify test–retest reliability, we computed Pearson correlations between the microstructural profiles from time point 1 and time point 2 for each subject and for each tract under both registration methods. This produced two correlation vectors per bundle, one for SLR and one for BundleWarp, each containing the test–retest correlation values across all subjects. For statistical comparison, we performed a paired *t*-test on these vectors for each tract. The paired *t*-test thus evaluated whether BundleWarp produced systematically higher within-subject test–retest correlations than SLR for each tract. FDR correction at *q* < 0.05 was applied across all bundles and metrics to account for multiple comparisons.

## 3. Results

The Results section is divided into two parts: the first details and evaluates the method and presents method comparisons, while the second focuses on tractometry and tract morphometry results.

First part summary: This section illustrates the results of BundleWarp registration on 1,728 pairs of bundles (including 27 different types of bundles) extracted from Parkinson’s Progression Markers Initiative (PPMI) dataMarek et al. (2011). We show in detail the impact of regularization, and results of *partially* and *fully* deformable bundlewarp registration. We compare the BundleWarp method with other streamline-based linear and image-based nonlinear registration methods. We show that BundleWarp outperforms based on qualitative (visual) and three quantitative metrics: bundle-based minimum distance, bundle shape similarity score, and volumetric DICE coefficient.

Second part summary: In this study, we integrated BundleWarp into the Bundle Analytics (BUAN) tractometry pipeline and evaluated the impact on results when BundleWarp is used versus when it is not used in BUAN. We utilized 3D diffusion MRI data from 730 participants in ADNI3 to study the impact of Mild Cognitive Impairment (MCI), dementia, amyloid, and tau on the microstructure of the brain’s white matter tracts. We conducted four experiments to detect associations of various variables on 36 white matter tracts of the brain. We tested microstructural associations with (1) MCI, (2) dementia, (3) amyloid positivity, and (4) tau positivity. This paper focuses on evaluating the performance of BundleWarp in tractometry, with results presented comparing BUAN outcomes with and without BundleWarp. We also show the application of BundleWarp in tract morphometry and bundle shape difference detection using the generated deformation field.

Please note that “AD” refers to axial diffusivity, and in this manuscript, we use the term “dementia” to refer to patients with Alzheimer’s disease.

### 3.1. *Partially*-to-*Fully* Deformable BundleWarp Registration

BundleWarp provides users with a free regularization parameter *λ* that can be tuned to get the desired level of deformations in the registration. In Fig. 8, we explore different values of *λ* and show its effect on the BundleWarp registration output. The top panel in Fig. 8 shows static (red) and moving (green) arcuate fasciculus (AF_L) bundles in the left hemisphere of the brain. The bottom panel in Fig. 8 shows static (red) and moving (green) arcuate fasciculus (AF_R) bundles in the right hemisphere of the brain. In both panels, the output of the BundleWarp registration with varying values of *λ* is shown in the first row, and the second row visualizes the displacement magnitude in mm projected over the moving bundle. *β* was set to 20 for all registrations. Yellow-red colors on the bundle indicate a higher amount of displacement introduced in the deformation step of the BundleWarp registration. Note that we only visualize the displacement generated by the deformation step (after affine registration) of BundleWarp as each point on the streamlines gets its warp map, unlike linear registration, where the same affine map is applied to all the points and streamlines. As we move from the left (*λ* = 0.5) to the right (*λ* = 0.001), we observe that the moving bundle is deformed more to match the static bundle. *λ* in range of [0.3-0.5] *partially* deforms the moving bundle. These values of *λ* warp the moving bundle to better align with the static bundle (compared to linear registration) and preserve the bundle’s original shape. *λ* in range of [0.001-0.1] *λ* < 0.001 *fully* deforms the moving bundle. These values of *λ* try to deform the moving bundle to completely overlap with the static bundle, and in achieving such registration, the moving bundle loses its original shape. From the second row in both panels, we can see that the extremities of the AF_L and AF_R needed more deformations than the middle part of the bundles to match the static bundles. Note that we only use *fully* deformable registration for bundle shape difference quantification through the displacement field. By default, we always use *λ* = 0.5 to only partially deform the bundle and preserve its anatomical shape.

**Figure 8:**
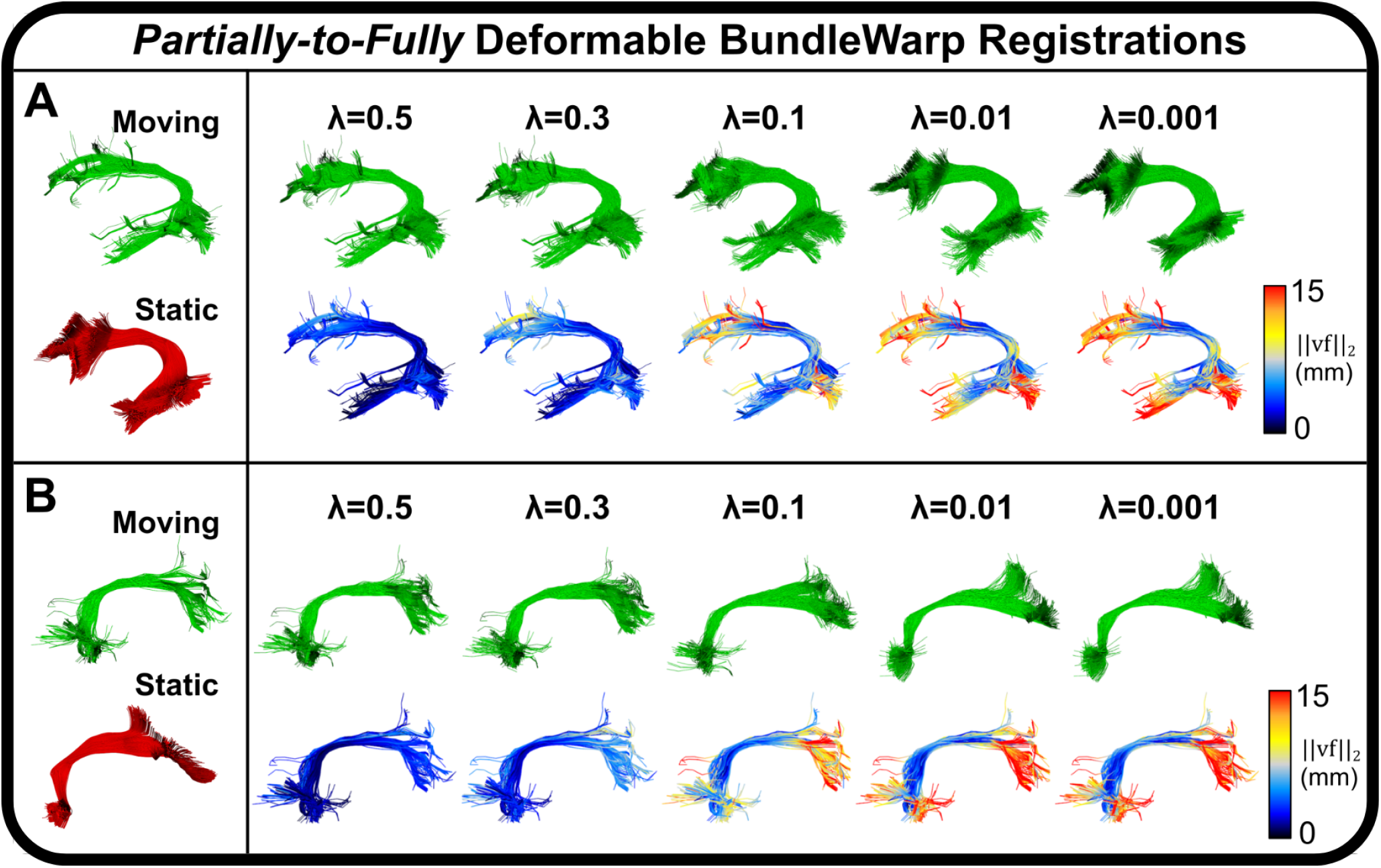
The effect of BundleWarp (BW) regularization free parameter *λ* on the registration. Arcuate fasciculus (green) in the left (top panel) and right (bottom panel) hemispheres of the brain are shown to be registered to a static bundle (red). Each panel shows BW registration output (first row in both panels) with varying values of the *λ*. As we move from left to right (higher value to a lower value of *λ*), more deformations are introduced in the registration. The second rows in both panels show displacement vector field magnitude visualized on the affinely aligned bundle. Here, yellow-to-red colors indicate areas on the bundles that were deformed the most. Areas that required higher deformations were the areas with the most shape differences in static and moving bundles. Note by default we always use *λ* = 0.5 in BundleWarp registration.

Figure 9 illustrates the *partially* deformable BundleWarp registration used in methods comparisons and BUAN tractometry. Here, (a) moving Cingulum bundle (in red color) is registered to static bundle (in green color) with *λ* = 0.5 and *β* = 10 (because the length of the cingulum bundle is less than 50mm) to improve linear registration (b) and, at the same time, preserve the original anatomical structure of the moving Cingulum bundle (c). In Fig. 9d, we visualize the vector field of deformations over an affinely registered moving bundle. Vector field arrows show how much and in which direction BundleWarp adds deformations after linear registration. Additionally, blue arrows in (c) point towards areas that are deformed and add significant improvement over the linear alignment of two bundles, yet the moving bundle retains its original characteristics. Here, a smooth and coherent vector field (d) indicates a smooth mapping of the streamlines, as we can see the structure of the Cingulum bundle in the vector field.

**Figure 9.**
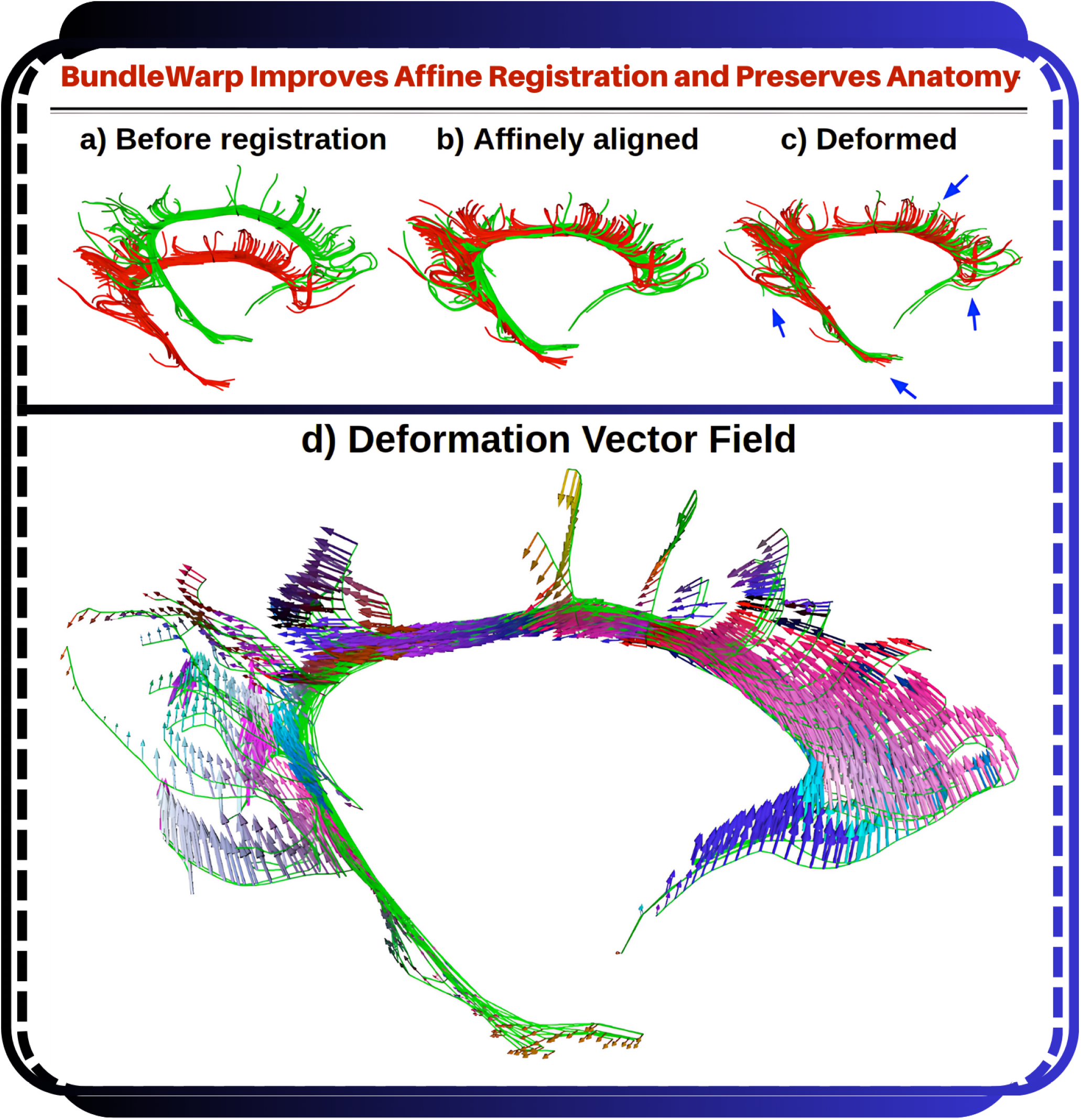
The BundleWarp method improves the linear registration by adding smooth deformations where needed and, at the same time, preserving the topological structure of the bundle. a) Shows input Cingulum static (red) and moving (green) bundles. b) Shows moving bundle affinely registered to static bundle. c) Shows deformations introduced by BundleWarp. Blue arrows indicate where deformations improved the output of affine registration. d) Shows a smooth velocity vector field visualized over an affinely registered bundle, as shown in b.

### 3.2. Method Comparisons

We performed BundleWarp registrations on 1,728 pairs of bundles. We extracted 27 white matter tracts from whole-brain tractograms of 64 subjects from PPMI data (a total of 1,728 tracts were extracted, 27 different types of bundles, 64 bundles per type). We used 27 atlas bundles (model/template bundles) from the HCP-842 template Yeh et al. (2018a) atlas as static bundles, and extracted bundles from 64 subjects were considered moving bundles (bundles to be registered to static bundle’s space). Static bundles were always kept the same coming from the HCP-842 atlas. Moving bundles come from different subjects. The same type of extracted bundles were registered to the same type of atlas bundle. For example, each subject’s arcuate fasciculus bundle was registered to the atlas arcuate fasciculus bundle. For full names of bundles see Fig.S3 in supplement. For BundleWarp registration of all the bundles, we kept *λ* = 0.5 and *β* = 20. NOTE: We kept a higher value of regularization, which means we introduced fewer deformations and did not completely deform the moving bundle.

For comparisons with other registration methods, we selected streamline-based linear registration (SLR) Gary-fallidis et al. (2015) and Advanced Normalization Tools (ANTs) Avants et al. (2009) methods. SLR provides direct streamline-based affine registration of white matter tracts. ANTs provides nonlinear registration of images. In order to apply image transformations to streamlines, we registered each subject’s FA image to FSL’sJenkinson et al. (2012) FMRIB58_FA template available at (https://fsl.fmrib.ox.ac.uk/fsl/fslwiki/FMRIB58_FA) using ANTs SyN registration using ANTspyAvants et al. (2009); Tustison, Cook, Holbrook, Johnson, Muschelli, Devenyi, Duda, Das, Cullen, Gillen et al. (2021). Our curated version of the HCP-842 bundle template Yeh et al. (2018a) is in the same space as the FMRIB58_FA template. The generic affine and warp map produced by ANTs SyN registration of two FA images were used to apply transformations to that subject’s 27 bundles using scilpy(https://github.com/scilus/scilpy).

This section presents qualitative and quantitative comparisons of three registration methods: BundleWarp, SLR, and ANTs. Fig. 10 shows qualitative/visual results and comparisons of the three methods. Here, we show visual results on the arcuate fasciculus (AF_L) and corticospinal tract (CST_L) in the brain’s left hemisphere selected randomly from one subject. In Fig. 10, static and moving bundles are visualized in panels A and C, followed by registered bundles by ANTs, SLR, and BundleWarp methods. The overlap of the static bundle with the moving bundle and the registration output of the three methods is visualized in panels B and D.

**Figure 10:**
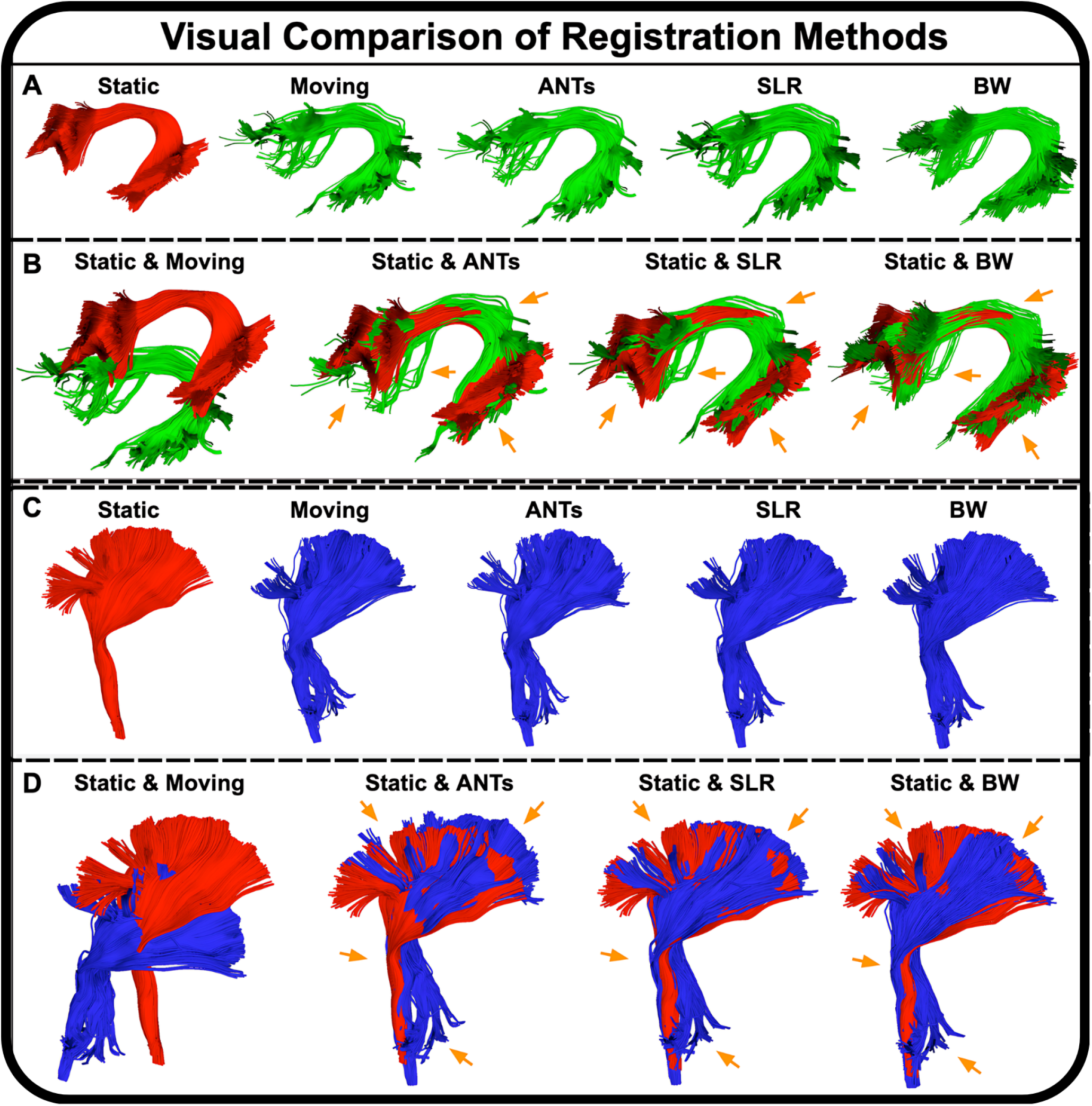
Visual comparison of BundleWarp (BW) with streamline-based linear registration method SLR, and image-based nonlinear registration method ANTs. (A and C) static, moving, ANTs output, SLR output, and BW output. (B and D) the static bundle is visualized with moving, ANTs output, SLR output, and BW output bundles. Orange arrows point at areas that are most affected by registration. BW produces the best registration results. *λ* was set to 0.5 in the BW method.

Panel A shows results on the AF_L bundle, whereas panel B shows results on the CST_L bundle. Here, orange arrows point toward the areas that are improved by registration methods. We can visually see that BundleWarp yields the optimal registration results for both bundles.

For a quantitative comparison of methods, we used two streamline-based metrics, bundle shape similarity metric (SM) and bundle-based minimum distance (BMD). We also used the volumetric dice similarity metric (DICE) to compare the overlap of fiber density images of tracts. Twentyseven bundles extracted from 64 subjects (1,728 total bundles) were registered to 27 atlas bundles of the same type. Three metrics were calculated between static and moved (registered) bundles.

To summarize the results of the quantitative comparisons, Fig. 11 presents boxplots summarizing the quantitative comparisons of ANTs (red), SLR (blue), and BundleWarp (green) methods using SM, BMD, and DICE metrics. Three panels (A, B, C) show boxplots of three metrics, respectively. Each panel has boxplots for a metric (y-axis) for 27 bundles (x-axis). Fig. 11a shows SM results. BundleWarp has the highest SM score. Fig. 11b shows BMD results. BundleWarp has the lowest BMD score (desirable). Fig. 11c shows DICE results. BundleWarp has the highest DICE score. Please see supplement Figure S4., Figure S5., and Figure S6. for a detailed quantitative comparison of each of the metrics; SM, BMD, and DICE.

**Figure 11.**
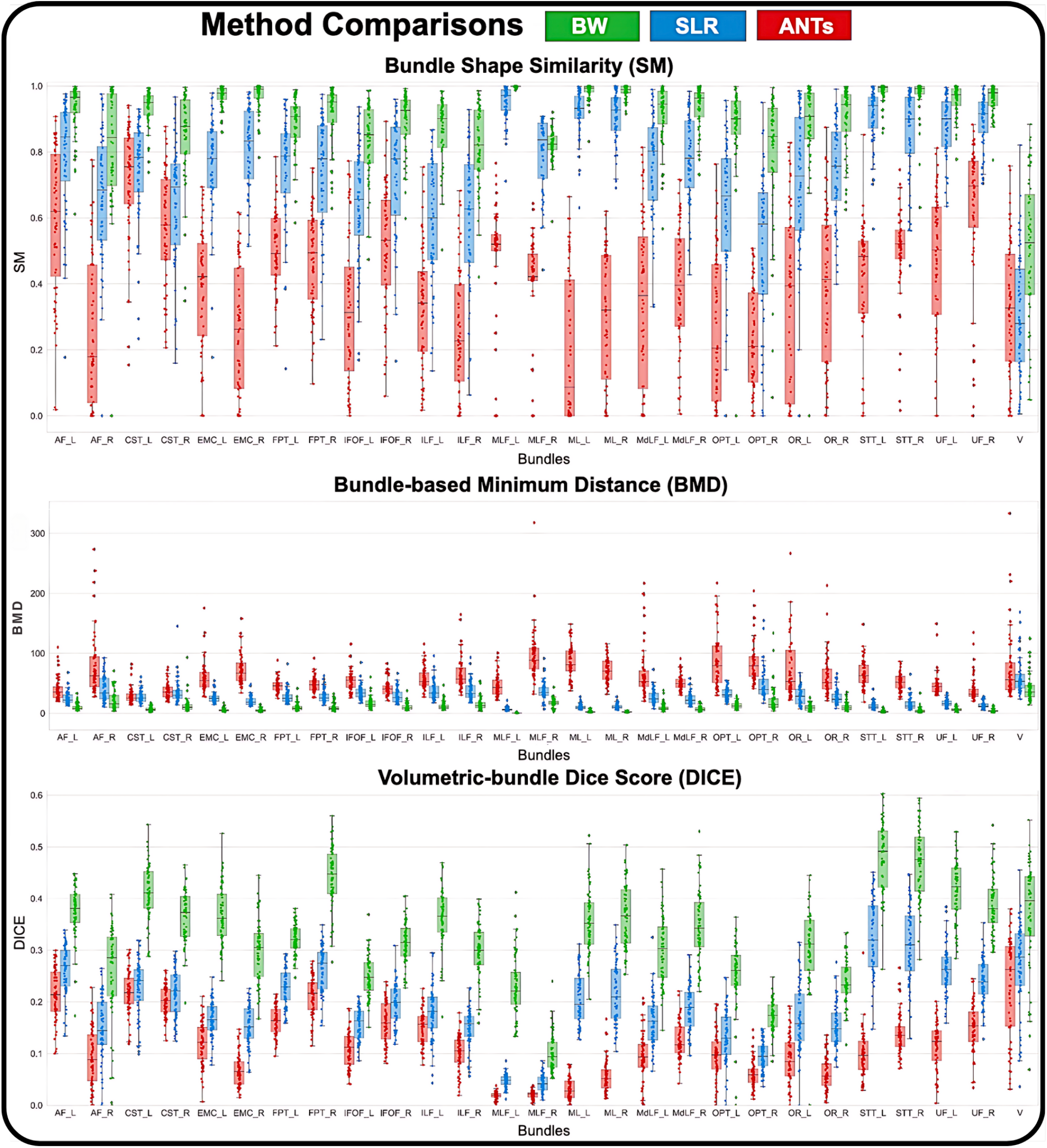
We present a quantitative comparison of the ANTs (red), SLR (blue), and BundleWarp (BW) (green) registration methods using three metrics: Shape Similarity Score (SM), Bundle-based Minimum Distance (BMD), and DICE coefficient. Each panel features boxplots for a metric (y-axis) across 27 bundles (x-axis). For SM and DICE, higher values indicate better performance, while for BMD, lower values are desirable.

We show with qualitative and quantitative methods that BundleWarp outperforms every other method in comparison. While SLR performed second best, and ANTs registration was placed at the end. Although SLR is a linear registration method, it still performs better than ANTs nonlinear registration. However, BundleWarp and SLR are direct streamline registration methods. While we applied ANTs SyN registration to FA images, the resulting affine and warp maps were used to transform streamlines. Direct streamline registration methods consider streamline geometry and shape, while image-based registration cannot incorporate streamline shape, size, and geometry into its method. We show that direct streamline registration is beneficial and outperforms image-based registration for streamlines.

Note that we kept the regularization term relatively high *λ* = 0.5. This means we introduced minimal deformations in the BundleWarp registration or, as we call it, *partially* deformed. If we set *λ* to a low value, such as *λ* = 0.0001, BundleWarp would completely deform the moving bundle to look like the static bundle. See Fig. 5 for the impact of *λ* on the registration. If we *fully* deform the bundle, we would get perfect SM and DICE scores of 1 and 0 BMD between the static and moved bundle. However, this is not advised and would completely change the shape and anatomy of the original bundles. Staying close to the original anatomy helps with statistical correspondence across the length of the bundles.

### 3.3. Tractometry

BundleWarp can deform bundles either *partially* or *fully*, depending on the regularization parameter *λ* (see methods section for details). In BUAN tractometry, we integrated *partially* deformable registration with *λ* = 0.5. BundleWarp improves tract alignment for robust tract segment correspondence while preserving the anatomical details of the bundles.

#### 3.3.1. Test–Retest Evaluation of Tract Correspondence and Subject Fingerprinting

To evaluate whether BundleWarp enhances tract correspondence and subject-level reproducibility, we performed a test–retest analysis using the HCP dataset (44 subjects with two diffusion MRI sessions each). See details of assessment methodology provided in Methods. Sec. 2.9

Using our bundle shape similarity framework, we quantified within-subject geometric agreement between session 1 and session 2 bundles under both SLR and BundleWarp alignment. Across the 36 bundles analyzed, BundleWarp consistently produced higher shape similarity scores than SLR. After false-discovery-rate (FDR) correction, the majority of bundles showed statistically significant improvements (Fig. 12a). The x-axis displays bundle indices, and the y-axis shows mean shape similarity for SLR (blue) and BundleWarp (green). Bundles marked with a red asterisk denote statistically significant improvements, indicating that BundleWarp yields more anatomically consistent and reliable geometric alignment across repeated scans.

**Figure 12.**
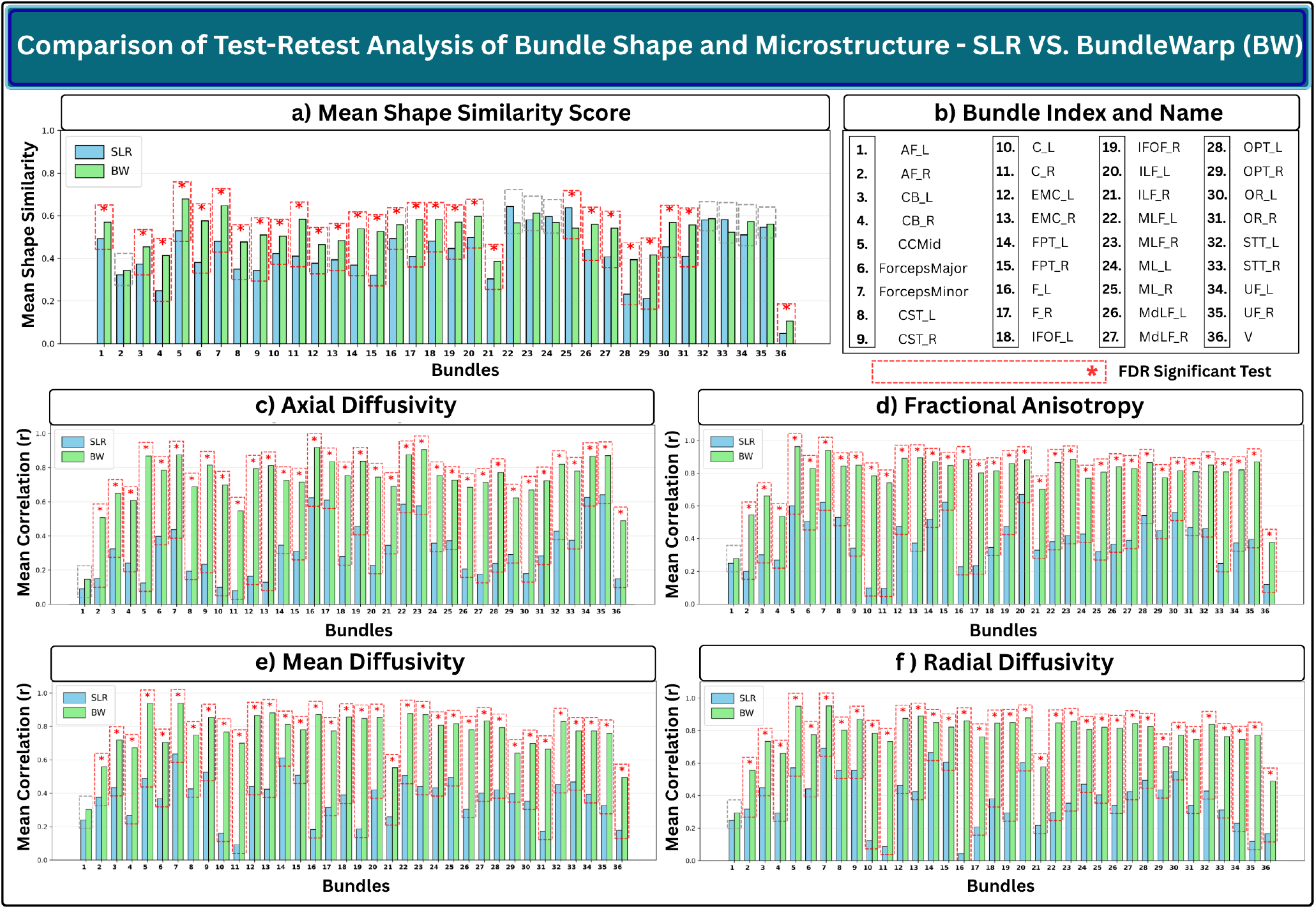
(a) Bundle shape similarity scores after Streamline-Based Linear Registration (SLR) and BundleWarp (BW). (b) Bundle indices and corresponding bundle names shown along the x-axis. (c–f) Pearson correlations of test–retest microstructural profiles for (c) FA, (d) MD, (e) RD, and (f) AD obtained using SLR and BW. Red boxes and asterisks indicate tracts showing statistically significant improvement after FDR correction as tested by paired *t*-test. Most bundles exhibit higher shape similarity and greater test–retest correlations with BW compared to SLR, demonstrating improved tract correspondence and reproducibility.

We further evaluated subject fingerprinting at the microstructural level using BUAN tractometry. For each diffusion metric (FA, MD, RD, AD), we extracted along-tract microstructural profiles for each bundle in both sessions and computed test–retest Pearson correlations under SLR and BundleWarp alignment. BundleWarp produced substantially higher correlations across all four metrics (Fig. 12, panels c–f), demonstrating stronger consistency of microstructural patterns across visits. A paired *t*-test, followed by FDR correction across all tests, showed BundleWarp consistently improved correlations, with many tracts showing statistically significant gains.

Together, these analyses demonstrate that BundleWarp enhances both geometric and microstructural test–retest reliability. Compared to linear SLR registration, BundleWarp provides more stable, anatomically meaningful, and subject-specific tract representations, strengthening its utility for precision white-matter tractometry analysis.

#### 3.3.2. Tractometry Alignment, Smoothness, and Reliability in Alzheimer’s Disease

To evaluate the impact of deformable bundle-specific registration on tractometry analysis, BUAN was executed twice: once incorporating the *partially* deformable BundleWarp registration within the pipeline and once using the standard SLR registration. We use 730 ADNI3 participants, see Methods Sec. 2.1 for data and preprocessing details. We compared FA, MD, RD, and AD microstructural measures along the tracts’ length to identify group differences between (1) MCI and cognitively normal control groups, (2) dementia and cognitively normal control groups, (3) amyloid-positive and amyloid-negative subjects, and (4) tau-positive and tau-negative subjects. We study their localized effects on 36 white matter tracts of the brain(see Fig.1 for abbreviations). 50 segments along the length of tracts are created in these four experiments. Linear Mixed Models are applied to 36 representative bundles with group, age, and sex modeled as fixed effects and subject and scanner modeled as random effect terms. Where the subject term accounts for multiple points on streamlines coming from the same subject, and the scanner term is used for mitigating scanner-based effects. The response variable is each DTI metric. We corrected for multiple comparisons using the false discovery rate (FDR) method Benjamini and Hochberg (1995). We propose two ways of calculating the FDR-corrected threshold for bundles, global and local. For each DTI metric, the global threshold is calculated by taking into account 50 segments from 36 bundles (1800 segments/*p-*values). The local threshold is bundle-specific, and we only utilize 50 segments of the bundle to calculate the FDR threshold. We show results using both FDR thresholds. Note that the nonlinearly moved bundles are only used to assign segment numbers to streamlines (and points on the streamlines) in the bundles. The actual statistical analysis always takes place in the native space of diffusion data. This step uses bundles of the original shape and microstructural measures in the native space, utilizing segment labels given during the assignment step for segment-specific group analysis.

Fig.13 consists of two panels that illustrate the effects of MCI and dementia on the Left Extreme Capsule (EMC_L) tract, comparing outcomes with BundleWarp and with SLR registration in BUAN tractometry. The EMC tract, located in the temporal lobe, is crucial for auditory and language processing Concha, Beaulieu and Gross (2005). Damage to this tract is associated with communication and language deficits, common in Alzheimer’s disease.

**Figure 13.**
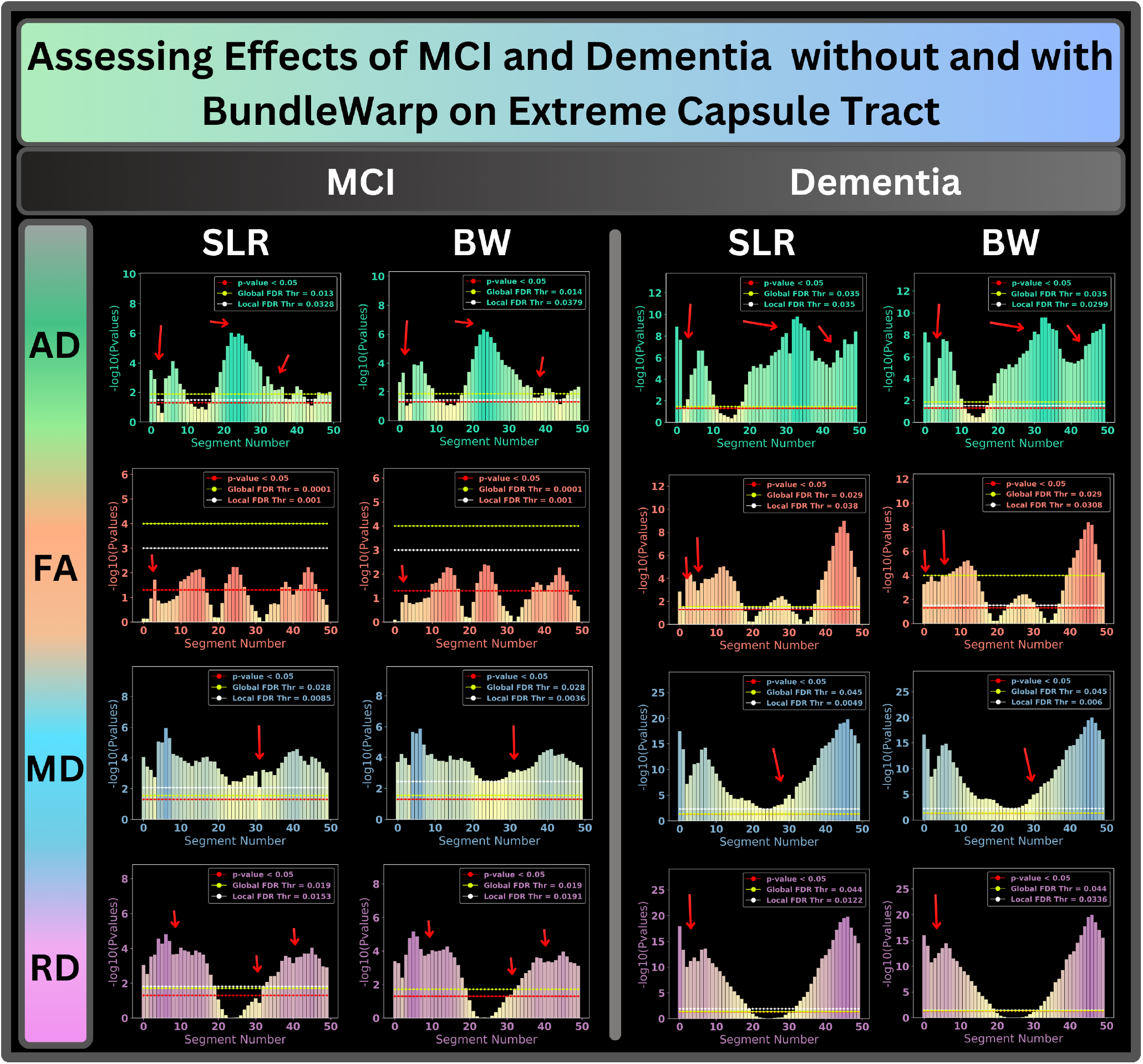
The two panels illustrate the effects of MCI and Dementia on the Left Extreme Capsule (EMC_L) tract, comparing results with streamline-based linear registration (SLR) and with BundleWarp nonlinear registration (BW) in BUAN tractometry. Each panel presents outcomes for four microstructural measures: axial diffusivity (AD), fractional anisotropy (FA), mean diffusivity (MD), and radial diffusivity (RD). The x-axis represents segment numbers along the tract, while the y-axis shows the negative logarithm of *p*-values. Red arrows highlight the segments where *p*-values improved with BW registration as compared to SLR in BUAN.

Each panel displays results for four microstructural measures: AD, FA, MD, and RD. The x-axis represents the 50 segment numbers along the tract, while the y-axis shows the negative logarithm of *p*-values. Red arrows indicate segments where *p*-values improve after incorporating BundleWarp registration. It enhances the alignment of tracts, enabling more precise anatomical correspondence across subjects and groups. This improved alignment ensures that comparable anatomical regions are analyzed, resulting in more reliable statistical outcomes. With SLR, certain areas in the *p*-value distribution exhibit discontinuities and abrupt shifts between segments, often due to misalignment or inconsistencies in tract correspondence. By minimizing these artifacts, BundleWarp produces a more continuous and consistent *p*-value trajectory, accurately reflecting the true underlying microstructural differences. Moreover, the incorporation of nonlinear registration does not introduce artifacts that significantly alter the results compared to those obtained with SLR.

Fig.14 presents the comprehensive BUAN results for the effects of tau positivity on the left cingulum (C_L) bundle, highlighting the impact of BundleWarp integration on the analysis of the AD metric. The cingulum bundle is essential for cognitive and emotional processing. Disruption of its microstructure can impair connectivity between the frontal lobe and other brain regions, which may contribute to the cognitive decline observed in Alzheimer’s disease patients Rose, Chen, Chalk, Zelaya, Strugnell, Benson, Semple and Doddrell (2000); Yang, Schilling, Shashikumar, Jasodanand, Moore, Pechman, Bilgel, Beason-Held, An, Shafer et al. (2023). The first column shows results with SLR, while the second column displays results with BundleWarp applied in BUAN. The top row plots display segment numbers on the x-axis, ranging from 1 to 50, and the negative logarithm of *p*-values on the y-axis. The horizontal blue line marks *p*-value< 0.05, with green and white lines indicating the FDR-corrected global and local significance thresholds, respectively. Segments with *p*-values passing these thresholds are considered significant and are highlighted in darker red on the plots. The last row visualizes the *p*-values projected along the length of the C_L bundle, where red/dark orange indicates lower *p*-values and greater group differences. Note that in this analysis, we focused only on the middle part of the C_L, which is more consistent in shape and easier to segment from clinical subjects. Consequently, the ends of the C_L were excluded from the statistical analysis because they do not consistently correspond to specific segments. The extremities of the C_L bundle are not represented in the *p*-value plots and are visualized in beige color in the 3D C_L visualizations shown in the last row.

**Figure 14.**
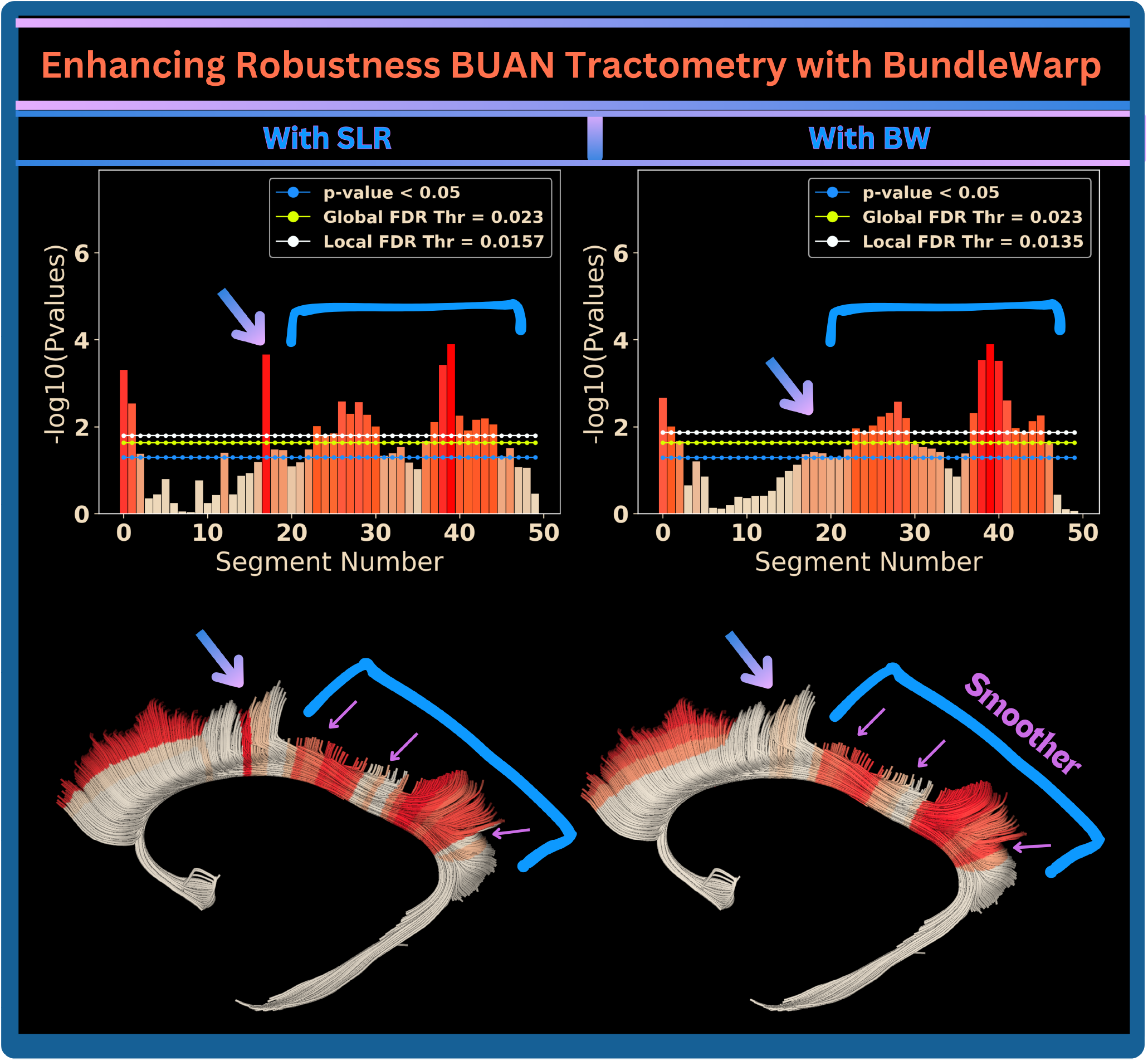
BUAN results for the left Cingulum bundle (C_L) with axial diffusivity (AD) metric are shown with SLR (first column) and with BundleWarp (second column). In the top row, the x-axis represents the segment number, while the y-axis shows the negative logarithm of the *p*-values. Horizontal green and white lines indicate the FDR-corrected global and local significance thresholds, respectively. The BundleWarp registration in BUAN tractometry leads to smoother *p*-values along the tract.

With BundleWarp registration, we observe higher coherence and a more continuous *p*-value distribution. Suboptimal registration, as shown in the first column, can lead to false positives, evidenced by sudden spikes in the *p*-value plot. Since neighboring segments within the bundle should exhibit similar behavior, it is unlikely for a single segment to show significant *p*-values when adjacent segments do not. As seen in the second column, incorporating BundleWarp registration results in a smoother *p*-value trajectory, free from unnatural variations between neighboring segments.

To summarize the impact of BundleWarp on the *p*-value curves of 36 bundles across 4 experiments, we use auto-correlation as a measure of the smoothness of each bundel’s *p*-value curve. Auto-correlation assesses how similar a signal (in this case, the *p*-value curve) is to a shifted version of itself. A high auto-correlation indicates that the *p*-value curve is smooth and consistent, while a low auto-correlation suggests irregularities or abrupt changes. We obtain one auto-correlation value per bundle, reflecting the overall smoothness of the *p*-value curve for that bundle.

In Fig.15, we show that BundleWarp registration enhances the smoothness and coherence of *p*-values along the length of the tracts as compared to SLR registration in BUAN tractometry across all four experiments, which focus on MCI, dementia, amyloid positivity, and tau positivity, respectively. Each of the four panels corresponds to a separate experiment, displaying results for four DTI metrics. The x-axis represents the bundles, and the y-axis shows the auto-correlation values of each bundle’s *p*-value curve. Auto-correlation values range from 0 to 1, where 1 indicates a higher auto-correlation (i.e., a smoother *p*-value curve) along the tract. Pink dots represent the auto-correlation of *p*-values when BUAN is run with SLR registration, while cyan dots show the auto-correlation with BundleWarp registration.

**Figure 15.**
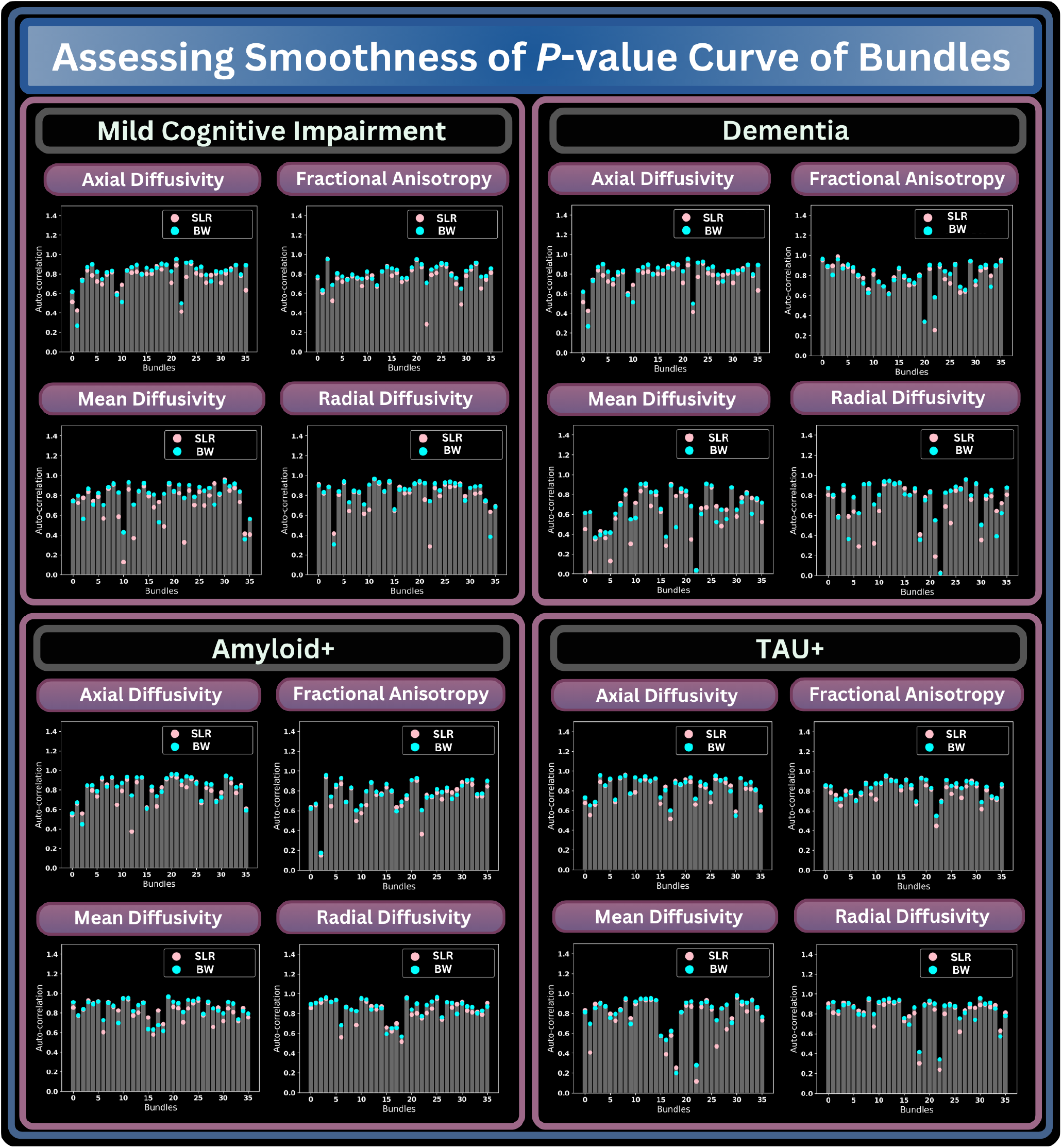
We evaluate the impact of BundleWarp (BW) on the smoothness and coherence of *p*-values along the tracts in BUAN tractometry as compared to streamline-based linear registration (SLR). Each panel displays results for four DTI metrics, with the x-axis representing bundles and the y-axis showing auto-correlation values ranging from 0 to 1. Pink dots indicate auto-correlation with SLR, while cyan dots show results with BW integrated into BUAN. Higher auto-correlation with BW indicates improved segment correspondence.

Fig.16 evaluates the impact of BundleWarp registration on the statistical power of BUAN. The four panels correspond to four distinct experiments examining the effects of MCI, dementia, amyloid positivity, and tau positivity on white matter tracts. Each panel displays results for four DTI metrics, with rows representing bundles and columns indicating 50 segments along the length of the tracts. We analyze the impact of BundleWarp on statistical power by examining changes in the *p*-value curves with BundleWarp compared to SLR in BUAN. A green pixel signifies an increase in significance for a specific bundle and segment, while a red pixel indicates a decrease. The color gradient reflects the magnitude of change in *p*-values: darker green represents a decrease in *p*-values (greater significance) with BundleWarp registration compared to SLR, while darker red indicates an increase in *p*-values (reduced significance).

**Figure 16.**
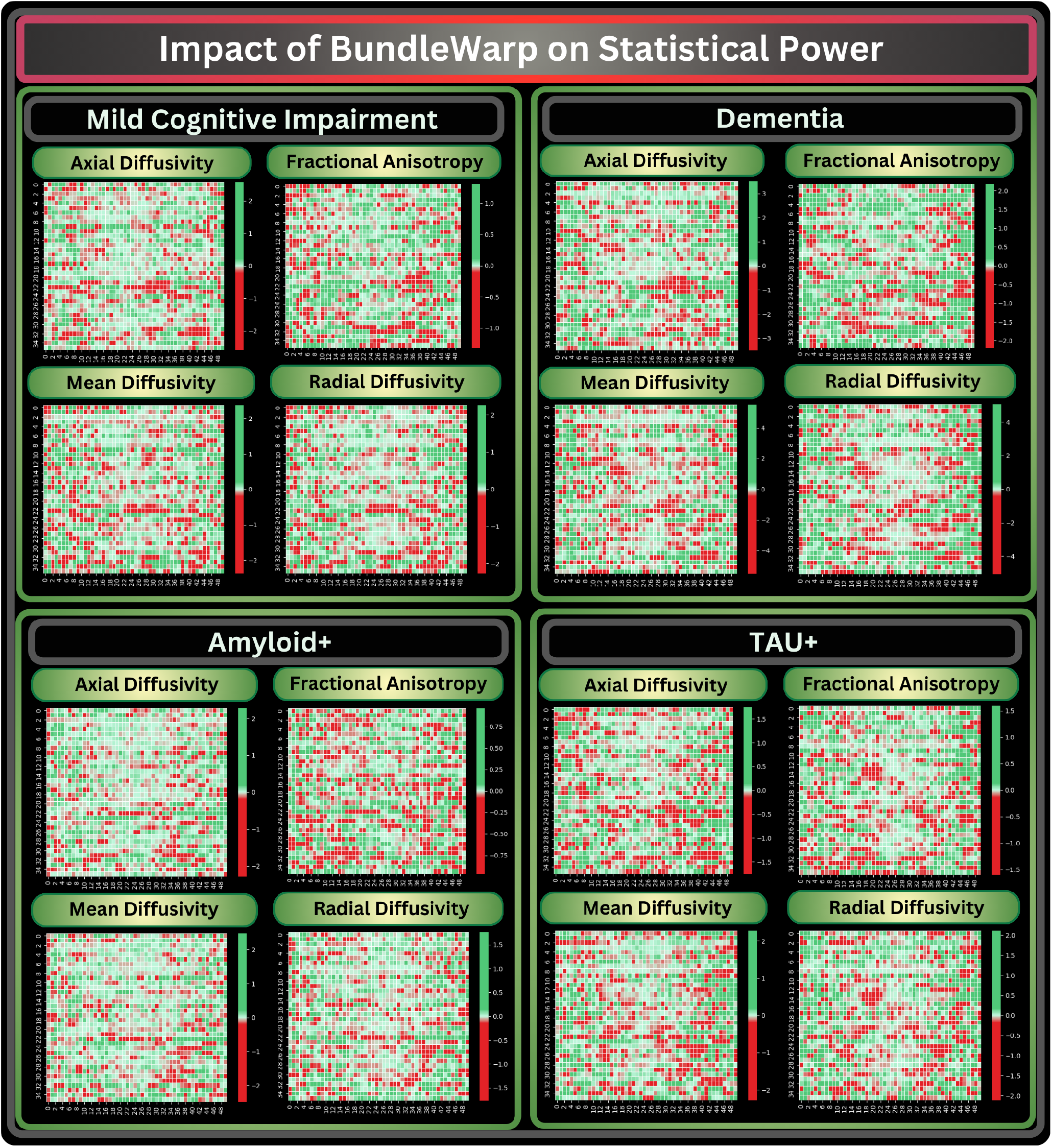
We evaluate the impact of BundleWarp (BW) on the statistical power of BUAN as compared to streamline-based linear registration (SLR). Each panel shows results for four DTI metrics, with rows representing bundles and columns indicating 50 segments along the tracts. Green pixels mark increased significance, while red pixels denote decreased significance. The color gradient reflects the magnitude of *p*-value changes, with darker green for greater significance and darker red for reduced significance with BW registration as compared to SLR.

Table 1 summarizes the improvements in statistical significance and smoothness across diagnostic and biomarker groups. BundleWarp consistently enhanced both the sensitivity and spatial coherence of tract-level analyses compared to SLR. In MCI, 61–81% of bundles showed higher mean negative logarithm of *p*-values across diffusion metrics (AD, FA, MD, RD), and 75–92% exhibited higher autocorrelation, indicating smoother along-tract profiles. Similar trends were observed in dementia, where 61–81% of bundles improved in significance and 69–75% in smoothness. Among amyloid-positive participants, 61–81% of bundles demonstrated greater significance and 61–72% improved smoothness, while in the tau-positive group, 61–81% of bundles also showed enhanced significance with 61–75% exhibiting smoother profiles. Overall, BundleWarp produced more spatially coherent and statistically robust tractometry patterns across all diffusion metrics and clinical groups, reinforcing its ability to improve both sensitivity and anatomical alignment in along-tract analyses. However, in some instances, a decrease in significance indicates that certain confounding effects, such as abrupt fluctuations in the *p*-value curves (as demonstrated in Fig. 14) caused by tract misalignment, may have been corrected through improved tract segment alignment with BundleWarp.

### 3.4. Tract Morphometry: Bundle Shape Difference Analysis

We conducted a bundle shape difference analysis using the displacement field generated by *fully* deformable BundleWarp. In this experiment, we used data from the PPMI database. See Methods Sec. 2.1 for data and preprocessing details. This analysis involved visualizing the magnitude of displacement on the moving bundle and quantifying bundle shape differences along the tract’s length using BUAN tractometry. This example illustrates individual-level analysis.

Figure 17 displays the bundle shape analysis for a randomly selected subject from the PPMI dataset. It includes four rows representing four bundles: left arcuate fasciculus (AF_L), uncinate fasciculus (UF_L), inferior longitudinal fasciculus (ILF_L), and optic radiation (OPT_L). Each column shows (a) the static bundles and (b) the moving bundles. We applied *fully* deformable BundleWarp registration (*λ* < 0.001) and visualized the magnitude of the displacement field over the moving bundle that was affinely registered to the static bundle (c). The blue-green areas indicate regions with higher displacement. A bundle shape differences profile is created using BUAN (d). 20 segments along the length of tracts are created. Note that each row (bundle) has different min and max displacement magnitude/y-axis (c-d) values. The analysis reveals that the extremities of the bundles generally require more displacement to align with the static bundle, indicating greater shape differences in these regions. This observation is consistent with the fact that the ends of some white matter tracts often approach or intersect with cortical and gray matter regions, such as those near the cortex. These areas may exhibit more variability in shape and alignment due to the transition from white matter tracts to the surrounding cortical and gray matter structures.

**Figure 17:**
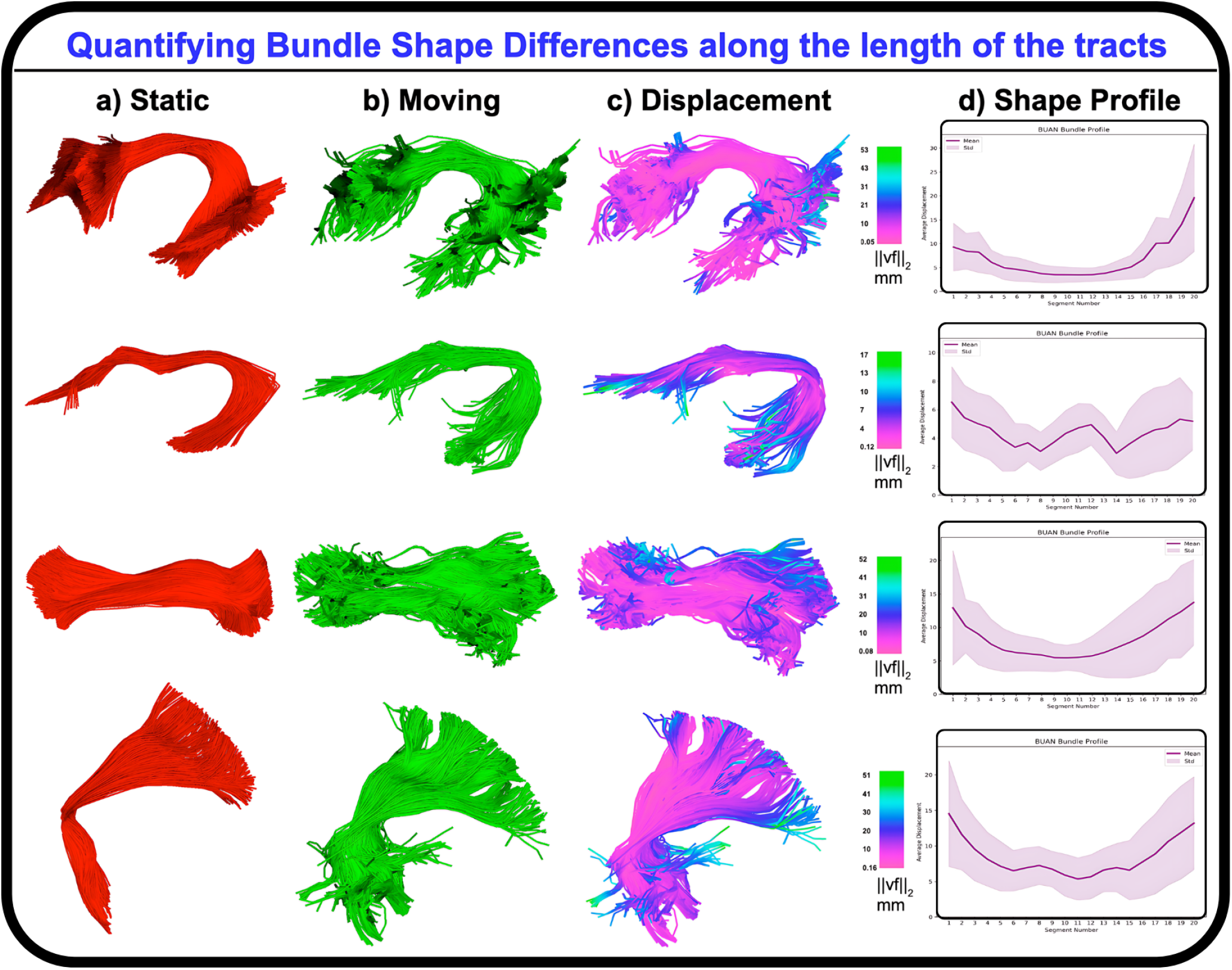
Quantifying bundle shape differences along the length of the tracts using displacement fields generated by BundleWarp. Four rows for four bundles, AF_L, UF_L, ILF_L, and OPT_L, respectively. Each column shows (a) static and moving bundles. *Fully* deformable BundleWarp registration is applied, and the generated displacement field’s magnitude is visualized on the moving bundle (c). A bundle shape differences profile is created using BUAN (d).

We conducted a bundle shape difference analysis at group-level using the displacement fields generated by the *fully* deformable BundleWarp framework, applied to diffusion MRI data from the ADNI3 cohort. Along-tract shape-difference profiles were derived from BundleWarp deformation fields and analyzed using the BUAN tractometry framework. We first evaluated the effect of amyloid positivity on white matter tract shape based on data from 329 amyloid-negative participants (CN: 235, MCI: 86, Dementia: 8; 156M, 173F) and 277 amyloid-positive participants (CN: 139, MCI: 87, Dementia: 51; 131M, 146F). In a parallel analysis, we assessed the effect of tau positivity on white matter tract shape using data from 401 tau-negative participants (CN: 293, MCI: 95, Dementia: 13; 192M, 209F) and 168 tau-positive participants (CN: 60, MCI: 68, Dementia: 40; 75M, 93F) from the same ADNI3 dataset. These analyses reveal spatially localized along-tract shape differences associated with amyloid and tau pathology, highlighting the sensitivity of BundleWarp morphometry to disease-related white-matter alterations.

Figure 18 displays the bundle shape analysis conducted using the *fully* deformable BundleWarp morphometry framework on ADNI3 diffusion MRI data. The first panel shows amyloid-positive versus amyloid-negative group comparisons, and the second panel shows tau-positive versus tau-negative comparisons. The x-axis represents segment position along each tract (1-50), and the y-axis indicates the negative logarithm of the *p*-values. Bundles exhibiting statistically significant shape differences after FDR correction are plotted, highlighting regions most affected by amyloid and tau pathology.

**Figure 18.**
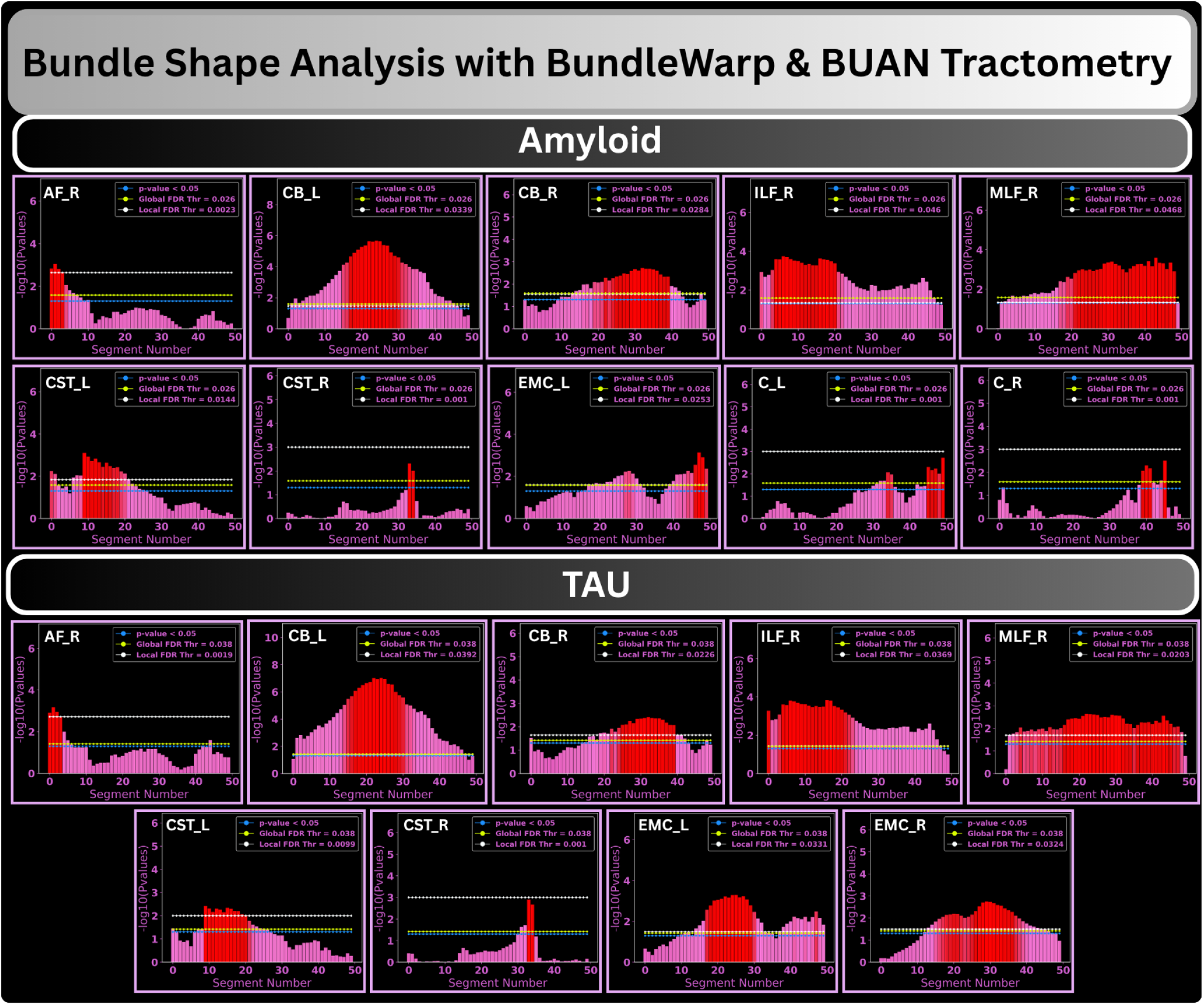
Quantifying bundle shape differences between amyloid-positive vs. amyloid-negative groups (first panel) and tau-positive vs. tau-negative groups (second panel) using the BundleWarp (BW) morphometry framework. Along-tract shape-difference profiles were derived from BW displacement fields and analyzed using the BUAN framework. The x-axis represents bundle segment position along the tract, and the y-axis shows the negative logarithm of the p-values. Tracts exhibiting statistically significant shape differences after FDR correction are highlighted/plotted, indicating regions most affected by amyloid and tau pathology.

## 4. Discussion

Nonlinear registration of any data, in general, is a non-trivial task. When dealing with medical data, a question arises: what is a perfect registration? From a mathematical point of view, a perfect registration should fully map moving data to be completely overlapped and look precisely like static/reference data. However, from a medical point of view, a perfect registration would be such that it aligns moving data to be in the same space as the static, and the overlap between them is higher than before. At the same time, moving data should not lose their original anatomical shape and characteristics. From a technical point of view, a desired method should not be memory or computation-heavy. Often, nonlinear registration methods are susceptible to introducing significant deformations to closely align input data to static/template data. For example, in the case of image registration, structure, and intensity values are often changed in moving images to match those of the static image. Neuroimaging data are widely used to understand the disease and find significant group differences between patient data and healthy controls. Most group analysis pipelines rely on nonlinear registration to align data and find differences among the corresponding areas. This could generate artifacts and/or remove the actual effects of the disease from the dataRohlfing (2006); Tsang, Dougherty and Wandell (2010) if registration completely changes the moving data to match the static data.

White matter bundles extracted from whole-brain tractograms of patient and control groups are analyzed using tractometry pipelines to find significant group differences along the length of the tracts. Like image-based group analysis methods, tractometry methods rely on linear or non-linear registration of white matter tracts or tractograms. Due to the lack of nonlinear streamline-based methods, most tractometry pipelines rely on image-based registra-tionYeatman et al. (2012); Yendiki et al. (2011); Cousineau et al. (2017); Dayan et al. (2015); Colby, Soderberg, Lebel, Dinov, Thompson and Sowell (2012). While some tractometry pipelines Chandio et al. (2020); Chandio (2022) have also utilized streamline-based linear registration. Hence, there is a need for a nonlinear streamline-based registration method. However, the nonlinear registration of tractograms/tracts needs careful evaluation. For example, if we fully deform a moving bundle to look exactly like a static bundle, we lose all the anatomical information specific to that subject. Moreover, if registration is sub-optimal and does not deform properly, it could make streamlines of the moving bundle spurious and introduce unnecessary artifacts.

We introduced a novel and robust method for *partially-to-fully* deformable registration of white matter tracts in the streamline-space. We created a method keeping in mind how important it is to preserve the anatomical structure of the tracts during nonlinear registration. By default, BundleWarp partially deforms the moving bundle and preserves the anatomical characteristics of the bundle. However, we provide a free regularization parameter *λ* that can control the level of deformations needed in the registration. *λ* decides the trade-off between regularizing the deformations and moving the bundle to match exactly like the static bundle. Using BundleWarp, a user can *partially* deform the bundle by setting a higher value of *λ* such as in the range of [0.2 −1] or they can *fully* deform the moving bundle by setting a low value of *λ* such as in the range of [0.1 − 0.00001]. There is another regularization parameter for smoothing, *β. β* determines the width of the Gaussian kernel. *β* decides the neighborhood of points on the streamline to consider when applying the topological constraint. Points closer to each other on the streamline will be moved as a group to preserve the topological structure of the tracts. For most of the bundles with an average length greater than 50 mm, *β* = 20 is sufficient and generates smooth transformations. By default, BundleWarp uses *β* = 20. For smaller bundles, such as the Cingulum bundle with an average length of less than 50 mm, BundleWarp automatically assigns *β* = 10 to yield the best results. To the best of our knowledge, BundleWarp is the only streamline-specific method for white matter tract registration that provides this capability. BundleWarp is a fast method that converges within 15 iterations of mlCPDs. For a typical bundle size with a streamline count in the range of [200-8000], BundleWarp execution time would be less than a minute or at most a minute. BundleWarp returns a non-linearly registered bundle (deformed), an affinely registered bundle, the deformations, and a pairwise MDF distance matrix with MDF distances among all the streamlines of static and moving bundles. It also returns many-to-one streamline correspondences among moving and static bundles. Pairwise distances and streamline correspondence can be further utilized in other methods, such as clustering and streamline filtering Chandio, Chattopadhyay, Owens-Walton, Reina, Nabulsi, Thomopoulos, Garyfallidis and Thompson (2021).

### 4.1. Advantages

Direct nonlinear alignment of the tracts in the streamline space has the main advantage of taking into account the topological connectivity of streamlines as opposed to voxel or volumetric-based representation of white matter tracts. Voxel-based representation and registration of tracts do not provide adequate precision because the tract size and shape vary considerably among subjects Wassermann et al. (2011). Therefore, it cannot ensure that a given voxel corresponds to the same tract across different subjects. For this reason, voxel/image-based registration methods, when applied to streamlines, would not be able to accurately align tractsGaryfallidis et al. (2015).

In our experiments, we compare BundleWarp with image-based nonlinear registration, Advanced Normalization Tools, ANTs SyNAvants et al. (2009), and streamline-based linear registration, SLRGaryfallidis et al. (2015). We show qualitatively and quantitatively that BundleWarp outperforms linear and nonlinear methods for the registration of white matter tracts. Even the linear streamline-based method SLR performs better than nonlinear ANTs normalization applied to streamlines. These outcomes are expected as ANTs registration is image-based and cannot take into account the geometrical shape and structure of individual streamlines. A single streamline travels through many voxels and integrates information across long regions. Hence, applying image-based affine and warp maps to streamlines results in sub-optimal alignment of the tracts. Whereas BundleWarp takes into account the streamlines’ topological connectivity and shape differences by constraining the optimization solution and making the algorithm more robust to outliers.

It is important to emphasize that this comparison is not intended as a critique of ANTs or a claim that BundleWarp replaces image-based registration. ANTs remains one of the most powerful and widely used tools for multimodal MRI alignment. Instead, BundleWarp is a streamline-specific approach that is designed to operate where voxelwise methods inherently lack anatomical precision, along individual fiber trajectories.

In our experiments, we show that *partially* deformable BundleWarp registration with *λ* = 0.05. Fig. 9d shows a smooth deformation vector field over an affinely moved Cingulum bundle. It shows that deformations added by BundleWarp after linear registration are coherent and smooth, as we can see in the structure of the Cingulum bundle in the vector field.

### 4.2. Preserving Anatomical Structure

Every subject is different and has unique white matter structural fingerprints. Most white matter bundles have a known shape, structure, function, and location in the brain that defines them. However, the same type of bundle can still have different shapes and sizes for different subjects. Deforming the original anatomical shape of the bundle is not necessarily ideal, as it can lose the characteristics needed in the analysis. For example, a patient with a brain injury, a tumor, or atrophies in the brain due to a neurodegenerative disease will have some pathways missing from the affected areas in the brain. Deforming the shape of the bundle too much, in these cases, can remove the disease/pathology effects, which are crucial to individual or group-level analysis. If every subject’s bundles start to look the same, we will lose the disease effects when dealing with groups of patients and healthy controls. Hence, it is of utmost importance to preserve most of the original anatomical structure of bundles in the registration process. BundleWarp preserves the core characteristics of the bundles, as shown in Fig. 9.

### 4.3. Recommended Parameter Values

In machine learning approaches, the regularization term *λ* is often selected using a cross-validation strategy to find its optimal value. For example, the Mean Squared Error (MSE) can be minimized across validation folds, and the *λ* that yields the minimal MSE is selected. Similarly, one could use a shape similarity score or a bundle-based distance metric (BMD) as the objective for selecting an optimal *λ* in BundleWarp registration. However, as discussed earlier in the manuscript, defining an “optimal” registration output is non-trivial. It is also difficult to define the optimal value of the cost function being minimized in cross-validation. An ideal shape similarity score of 1 or a BMD of 0 would indicate perfect overlap, but would result in over-deformation and unrealistic bundle shapes. Therefore, we empirically found that *λ* values in the range of [0.3–0.5] produce stable and anatomically plausible registration results. All comparison results in this study used *λ* = 0.5.

BundleWarp is a powerful method that can fully deform the moving bundle to look exactly like the static bundle. However, it should not be used to completely deform the bundle. By default, BundleWarp will only partially deform the moving bundle to align better with the static bundle. We recommend *λ* = 0.3 (or higher, such as *λ* = 0.5), and 0.5 is the default value of *λ* in the BundleWarp method. Since *λ* is not scale-dependent (scale of bundles) and controls the fluidity of the data points, we kept it the same for all bundles regardless of their shapes and sizes. In case a user sets *λ* to a value such as *λ* < 0.2, a warning will be displayed on the console explaining and making the user aware of the effects of setting a lower value of *λ* on the data. We have kept the default value of *β* depending on the average length of the bundle. BundleWarp automatically sets *β* = 10 when bundle length < 50 mm and *β* = 20 for bundles with length > 50 mm. BundleWarp converges within 15 iterations, and the default number of maximum iterations is set to 15 as well.

Different bundles may require varying deformation levels based on anatomical variability or pathology, and can be adapted by the user. A limitation of the current implementation is the use of a single global regularization parameter (*λ*) that uniformly controls deformation across all bundles. While users can manually adjust *λ* based on their data, future work could enable adaptive or bundle-specific tuning to improve alignment in heterogeneous or disease-affected regions.

### 4.4. BundleWarp RunTime

BundleWarp is designed for fast deformable registration of white-matter bundles across a wide range of bundle sizes. Because bundles can vary substantially in the number of constituent streamlines, even within the same tract type, we evaluated BundleWarp’s runtime as a function of streamline count on a standard workstation (32 GB RAM; Intel Core i7-7700K, 8 cores). A detailed analysis is provided in Supplement S.2.

Briefly, when registering subject bundles to an atlas (the most common use case), the atlas bundle is static, and the moving bundle varies. BundleWarp consistently completes in a few seconds when the moving bundle contains fewer than 4,000 streamlines, typical for most tractography-derived bundles, and remains well under one minute. When the size of both static and moving bundles increases, runtime remains in the seconds range for bundle sizes below 4,000 streamlines, and increases gradually as both bundles approach 10,000 streamlines. Even at these larger sizes, BundleWarp completes within 2–3 minutes. Only wholebrain tractogram–level registrations would incur substantially higher runtimes, which is outside BundleWarp’s intended scope.

These runtimes are enabled by BundleWarp’s two-stage strategy: (1) fast streamline-to-streamline correspondence estimation, followed by (2) deformation using mlCPD, which is highly efficient for the small point clouds represented by individual streamlines. Please see Supplement Section S.2. for more details on runtime experiments.

### 4.5. Applications of BundleWarp

BundleWarp can be integrated into various tractography-based methods, including bundle segmentation, bundle shape analysis, bundle atlasing, and tractometry.

In this paper, we demonstrate two applications of BundleWarp registration. First, we incorporated BundleWarp into the BUAN tractometry pipeline to investigate the effect of non-linear registration on tractometry analysis. Tractometry involves creating segments or nodes along the length of bundles, which are independently analyzed under the assumption that corresponding segments represent the same areas in the bundle across subjects. BundleWarp improves the robustness of BUAN tractometry by precisely aligning bundles with template bundles, ensuring reliable and consistent results. Second, we introduce a novel application of the displacement vector field generated by *fully* deformable BundleWarp registration for quantifying differences in bundle shape.

#### 4.5.1. Tractometry

We show that BundleWarp can be easily incorporated into our BUAN tractometry Chandio et al. (2020) pipeline. BUAN extracts bundles from populations and analyzes the microstructural measures projected onto the bundles along the length of the tracts to find significant group differences. BUAN creates horizontal segments along the length of the tracts to be analyzed. Horizontal segments are created based on points on the streamlines belonging to the closest model bundle centroid point. Here, a *partially* deformable BundleWarp is added to nonlinearly align subjects’ bundles with model bundles to find better segment correspondences among populations.

We visually and analytically demonstrate that BundleWarp is a valuable addition to the BUAN tractometry pipeline for robust tractometric analysis. We conducted four experiments, once with SLR registration and once with BundleWarp registration, to assess localized white matter microstructural alterations along the length of 36 tracts due to (1) MCI, (2) Dementia, (3) amyloid positivity, and (4) tau positivity, using data obtained from ADNI3. Nonlinearly aligning subjects’ bundles to model bundles with BundleWarp improved along-tract segment correspondence as compared to linear affine SLR registration, increasing the sensitivity of group statistical analyses by eliminating errors due to misalignment across subjects. Auto-correlation analysis is employed to evaluate the smoothness of *p*-value plots, showing that BundleWarp registration results in more consistent and smoother *p*-value curves. This improvement is reflected in the reduction of sudden spikes in *p*-values across segments, which can arise from misalignment or spurious outlier streamlines. The incorporation of BundleWarp effectively generates smoother *p*-value distributions, enhancing overall analytical accuracy and robustness. BundleWarp could also be utilized similarly in other tractometry pipelines Yeatman et al. (2012); Yendiki et al. (2011).

Moreover, our test–retest experiments show that BundleWarp substantially enhances subject-level fingerprinting of white-matter organization. By improving anatomical correspondence across sessions, BundleWarp yields higher within-subject shape similarity and markedly stronger reproducibility of along-tract microstructural profiles across all diffusion metrics. These improvements translate into clearer and more reliable subject-specific signatures, reducing variability that typically arises from misalignment rather than true biological change. Such stability is especially important for longitudinal studies, where detecting subtle within-subject changes over time requires highly consistent tract representations. In addition, improved subject fingerprinting benefits applications such as individualized modeling, subject identification, developmental and aging trajectories, and monitoring treatment or disease progression. Together, these findings suggest that nonlinear, bundle-specific registration is essential for precision neuroimaging and for analyses that depend on robust subject-level correspondence.

#### 4.5.2. Tract Morphometry: Bundle Shape Analysis

We propose a novel approach for quantifying bundle shape differences using *fully* deformable BundleWarp registration. We project and examine the displacement field magnitudes as they change along the length of the tracts using BUAN tractometry. We show an example of how the deformation field can be used to learn bundle shape differences. This is an exceptional application where we do recommend *fully* deforming the moving bundle by using *λ* < 0.001 for accurately quantifying bundle shape differences. When the moving bundle is completely deformed to look exactly like the static bundle, the deformation field can be visualized to show how much the points on streamlines were moved/deformed to match the static bundle, as explained and shown in Fig. 7 and Fig. 17.

However, the displacement field has been used before to quantify bundle shape differences. For example, the authors of the paperGlozman et al. (2018) used a seed point matching strategy to establish correspondences between bundle regions to perform registration and compute shape differences. In contrast, BundleWarp uses streamline-matching, and deformations are added, taking into account the entire streamline/bundle. Furthermore, we add a step in which local deformations are averaged along the length of the bundle. In this paper, we examined bundle shape differences at both the individual and group levels. Using PPMI data, we performed subject-specific shape analysis, both visually and quantitatively, by evaluating displacement-magnitude profiles along the tract. At the group-level, using the large ADNI3 cohort (N=730), we assessed structural bundle shape differences associated with amyloid-positive and tau-positive status. Together, these experiments demonstrate that BundleWarp’s morphometry framework can detect subtle, spatially localized shape alterations that are relevant to aging, neurodegeneration, and longitudinal disease progression. Other measures derived from vector fields, such as Jacobian determinant maps Ashburner and Friston (2000); Ashburner (2009), will also be incorporated in bundle shape analysis.

### 4.6. Limitations and Future Work

In future research, we plan to investigate the application of BundleWarp registration to Bundle Atlasing. Bundle Atlasing Romero-Bascones, Qamar Chandio, Fadnavis, Sung Park, Koudoro, Ayala, Barrenechea and Garyfallidis (2022) is a method recently proposed for constructing population- and bundle-specific streamline atlases, involving two main steps: unbiased registration and bundle combination Romero-Bascones et al. (2022). Currently, affine registration is carried out using SLR, but integrating a *partially* deformable BundleWarp registration could improve the accuracy of bundle matching. This improvement would potentially lead to atlases that better represent the underlying anatomy of the studied group. Thus, exploring the impact of BundleWarp on bundle atlasing represents a promising avenue for future research.

In this study, we focused on the registration of specific bundles due to their significance in current tractometric research Yeatman et al. (2012); Chandio et al. (2020); Cousineau et al. (2017); Yendiki et al. (2011); Chandio et al. (2022); Chandio, Villalon-Reina, Nir, Thomopoulos, Feng, Benavidez, Jahanshad, Harezlak, Garyfallidis and Thompson (2024). BundleWarp is designed for the nonlinear registration of individual white matter tracts and operates efficiently with relatively short execution times. While it could, in theory, be applied to whole-brain tractograms, this would dramatically increase execution time due to the extensive pairwise minimum-average direct flip (MDF) distanceGaryfallidis et al. (2012) calculations needed for streamline correspondences. Future research will explore extending the BundleWarp framework for whole-brain tractogram alignment by incorporating methods for faster streamline distance calculations St-Onge, Garyfallidis and Collins (2021); Wang (2022) and investigating approaches such as clustering whole-brain tractograms and aligning individual clusters with BundleWarp Olivetti et al. (2020).

BundleWarp uses a single global regularization parameter *λ* that provides a stable and consistent deformation model, which helps preserve anatomical coherence across the bundle. Empirically, we find that this formulation performs robustly across a wide range of datasets and bundle geometries, including cases with moderate anatomical variability. While this formulation performs well in practice, it does not explicitly model spatial variability in deformation requirements. This may be relevant in regions with complex geometries, such as fanning fiber structures. Nevertheless, the motion coherence constraint mitigates abrupt local deformations, and future work will explore spatially adaptive extensions.

## 5. Conclusion

BundleWarp is an efficient and reliable streamline-based nonlinear registration method for the registration of individual white matter tracts. It generates warping fields that preserve crucial anatomical structures within bundles, with the flexibility to *partially* or *fully* deform bundles by adjusting a single regularization parameter. Our qualitative and quantitative results show that BundleWarp outperforms other streamline-based and image-based registration methods for white matter tracts. The integration of BundleWarp into the BUAN tractometry pipeline reduces misalignment errors along bundles and their segments under diverse experimental conditions, delivering consistent and precise results. Furthermore, our test–retest fingerprinting experiments demonstrate that BundleWarp substantially enhances within-subject reproducibility at both geometric and microstructural levels, enabling more stable subject-specific tract representations. Using data from Parkinson’s and Alzheimer’s disease cohorts, we showcase its applicability in clinical research. Moreover, by incorporating amyloid and tau as early-stage biomarkers of Alzheimer’s disease, we highlight BundleWarp’s ability to detect subtle structural associations in preclinical stages, offering new insights into disease progression. Additionally, the displacement fields generated by BundleWarp facilitate both visual and analytical quantification of bundle shape differences, providing a tract morphometry framework.

## Data and materials availability

Code and tutorials for BundleWarp are publicly available through DIPYGaryfallidis, Brett, Amirbekian, Rokem, van der Walt, Descoteaux, Nimmo-Smith and Dipy Contributors (2014c) software in both Pythonic API and command-line API.

BundleWarp code can be found at https://github.com/dipy/dipy. Pythonic API tutorial: https://docs.dipy.org/stable/examples_built/registration/bundlewarp_registration.html. Command-line interface: https://docs.dipy.org/stable/interfaces/bundlewarp_registration_flow.html. PPMI data is publicly available at: www.ppmi-info.organd PPMI data derivatives Chandio (2020) used in the experiments can be found here: https://doi.org/10.35092/yhjc.12033390.v1. ADNI3 data is publicly available at https://adni.loni.usc.edu/adni-3/.

## Acknowledgements

We would like to acknowledge that the research reported in this publication was supported by the National Institutes of Health (NIH) under award numbers R01 EB027585, U01 AG068057, P41 EB015922, RF1 AG057892, RF1 NS136995, and the US Alzheimer’s Association under grant AARG-23-1149996. We would like to acknowledge that data used in the preparation of this article were obtained from the Alzheimer’s Disease Neuroimaging Initiative (ADNI) database (adni.loni.usc.edu). As such, the investigators within the ADNI contributed to the design and implementation of ADNI and/or provided data, but most of them did not participate in the analysis or writing of this report. A complete listing of ADNI investigators may be found at: http://adni.loni.usc.edu/wp-content/uploads/how_to_apply/ADNI_Acknowledgement_List.pdf

## Author Contributions

Conceptualization: B.Q.C, E.G, and J.H. Methodology: B.Q.C, E.G, D.R.B, and E.O. Implementation: B.Q.C. Data processing: B.Q.C, S.I.T, J.E.VR, and T.M.N Experiments and investigation: B.Q.C and D.R.B. Writing-original draft: B.Q.C, E.G, and D.R.B. Writing-review and editing: P.M.T, and S.I.T.

## Competing Interests

The authors declare that they have no competing financial interests.

## Notes

https://docs.dipy.org/1.12.0/examples_built/registration/bundlewarp_registration.html

### Competing Interest Statement

The authors have declared no competing interest.

### Summary of Updates

New figures and experiments have been added in this revised version.

## References

Albert, M., Antel, J., Brück, W., Stadelmann, C., 2007. Extensive cortical remyelination in patients with chronic multiple sclerosis. Brain pathology 17, 129–138.

Alexander, A.L., Lee, J.E., Lazar, M., Field, A.S., 2007. Diffusion tensor imaging of the brain. Neurotherapeutics 4, 316–329.

Andersson, J.L., Sotiropoulos, S.N., 2016. An integrated approach to correction for off-resonance effects and subject movement in diffusion MR imaging. Neuroimage 125, 1063–1078.

Arsigny, V., Commowick, O., Ayache, N., Pennec, X., 2009. A fast and log-euclidean polyaffine framework for locally linear registration. Journal of Mathematical Imaging and Vision 33, 222–238.

Ashburner, J., 2009. Computational anatomy with the spm software. Magnetic resonance imaging 27, 1163–1174.

Ashburner, J., Friston, K.J., 2000. Voxel-based morphometry—the methods. Neuroimage 11, 805–821.

Avants, B.B., Tustison, N., Song, G., 2009. Advanced normalization tools (ANTS). Insight J 2, 1–35.

Bach, M., Laun, F.B., Leemans, A., Tax, C.M., Biessels, G.J., Stieltjes, B., Maier-Hein, K.H., 2014. Methodological considerations on tract-based spatial statistics (tbss). NeuroImage 100, 358–369.

Basser, P.J., Mattiello, J., LeBihan, D., 1994. MR diffusion tensor spectroscopy and imaging. Biophysical Journal 66, 259–267.

Basser, P.J., Pajevic, S., Pierpaoli, C., Duda, J., Aldroubi, A., 2000. In vivo fiber tractography using DT-MRI data. Magnetic resonance in medicine 44, 625–632.

Bells, S., Cercignani, M., Deoni, S., Assaf, Y., Pasternak, O., Evans, C., Leemans, A., Jones, D., 2011. Tractometry–comprehensive multi-modal quantitative assessment of white matter along specific tracts, in: Proc. ISMRM, p. 1.

Benjamini, Y., Hochberg, Y., 1995. Controlling the false discovery rate: a practical and powerful approach to multiple testing. Journal of the Royal statistical society: series B (Methodological) 57, 289–300.

Brück, W., 2005. The pathology of multiple sclerosis is the result of focal inflammatory demyelination with axonal damage. Journal of neurology 252, v3–v9.

Canales-Rodríguez, E.J., Daducci, A., Sotiropoulos, S.N., Caruyer, E., Aja-Fernández, S., Radua, J., Yurramendi Mendizabal, J.M., Iturria-Medina, Y., Melie-García, L., Alemán-Gómez, Y., et al., 2015. Spherical deconvolution of multichannel diffusion MRI data with non-Gaussian noise models and spatial regularization. PLoS ONE 10, e0138910.

Cassoli, J.S., Guest, P.C., Malchow, B., Schmitt, A., Falkai, P., Martins-de Souza, D., 2015. Disturbed macro-connectivity in schizophrenia linked to oligodendrocyte dysfunction: from structural findings to molecules. npj Schizophrenia 1, 1–10.

Catani, M., De Schotten, M.T., 2008. A diffusion tensor imaging tractography atlas for virtual in vivo dissections. cortex 44, 1105–1132.

Catani, M., de Schotten, M.T., 2012. Atlas of human brain connections. Oxford University Press.

Chandio, B., 2020. Dipy processed parkinson’s progression markers initiative (ppmi) data derivatives. URL: 10.35092/yhjc.12033390.v1.

Chandio, B.Q., 2022. Advancing White Matter Tractometry of the Brain Using Diffusion MRI and Machine Learning. Ph.D. thesis. Indiana University.

Chandio, B.Q., Chattopadhyay, T., Owens-Walton, C., Reina, J.E.V., Nabulsi, L., Thomopoulos, S.I., Garyfallidis, E., Thompson, P.M., 2021. Fiberneat: unsupervised streamline clustering and white matter tract filtering in latent space. bioRxiv.

Chandio, B.Q., Chattopadhyay, T., Owens-Walton, C., Reina, J.E.V., Nabulsi, L., Thomopoulos, S.I., Garyfallidis, E., Thompson, P.M., 2022. Fiberneat: Unsupervised white matter tract filtering, in: 2022 44th Annual International Conference of the IEEE Engineering in Medicine & Biology Society (EMBC), IEEE. pp. 5055–5061.

Chandio, B.Q., Garyfallidis, E., 2020. Stnd: Streamline-based non-rigid partial-deformation tractography registration. Medical Imaging Meets NeurIPS.

Chandio, B.Q., Risacher, S.L., Pestilli, F., Bullock, D., Yeh, F.C., Koudoro, S., Rokem, A., Harezlak, J., Garyfallidis, E., 2020. Bundle analytics, a computational framework for investigating the shapes and profiles of brain pathways across populations. Scientific Reports 10, 17149.

Chandio, B.Q., Villalon-Reina, J.E., Nir, T.M., Thomopoulos, S.I., Feng, Y., Benavidez, S., Jahanshad, N., Harezlak, J., Garyfallidis, E., Thompson, P.M., 2024. Bundle analytics based data harmonization for multi-site diffusion mri tractometry. bioRxiv , 2024–02.

Colby, J.B., Soderberg, L., Lebel, C., Dinov, I.D., Thompson, P.M., Sowell, E.R., 2012. Along-tract statistics allow for enhanced tractography analysis. Neuroimage 59, 3227–3242.

Concha, L., Beaulieu, C., Gross, D.W., 2005. Bilateral limbic diffusion abnormalities in unilateral temporal lobe epilepsy. Annals of neurology 57, 188–196.

Cousineau, M., Jodoin, P.M., Garyfallidis, E., Côté, M.A., Morency, F.C., Rozanski, V., Grand’Maison, M., Bedell, B.J., Descoteaux, M., 2017. A test-retest study on parkinson’s ppmi dataset yields statistically significant white matter fascicles. NeuroImage: Clinical 16, 222–233.

Dayan, M., Monohan, E., Pandya, S., Kuceyeski, A., Nguyen, T.D., Raj, A., Gauthier, S.A., 2015. Profilometry: A new statistical framework for the characterization of white matter pathways, with application to multiple sclerosis. Human brain mapping.

De Santis, S., Drakesmith, M., Bells, S., Assaf, Y., Jones, D.K., 2014. Why diffusion tensor mri does well only some of the time: variance and covariance of white matter tissue microstructure attributes in the living human brain. Neuroimage 89, 35–44.

De Vos, K.J., Grierson, A.J., Ackerley, S., Miller, C.C., 2008. Role of axonal transport in neurodegenerative diseases. Annu. Rev. Neurosci. 31, 151– 173.

Dean III, D.C., Sojkova, J., Hurley, S., Kecskemeti, S., Okonkwo, O., Bendlin, B.B., Theisen, F., Johnson, S.C., Alexander, A.L., Gallagher, C.L., 2016. Alterations of myelin content in parkinson’s disease: a crosssectional neuroimaging study. PloS one 11, e0163774.

Durrleman, S., Fillard, P., Pennec, X., Trouvé, A., Ayache, N., 2011. Registration, atlas estimation and variability analysis of white matter fiber bundles modeled as currents. NeuroImage 55, 1073–1090.

Durrleman, S., Prastawa, M., Charon, N., Korenberg, J.R., Joshi, S., Gerig, G., Trouvé, A., 2014. Morphometry of anatomical shape complexes with dense deformations and sparse parameters. NeuroImage 101, 35–49.

Farquharson, S., Tournier, J.D., Calamante, F., Fabinyi, G., Schneider-Kolsky, M., Jackson, G.D., Connelly, A., 2013. White matter fiber tractography: why we need to move beyond dti. Journal of neurosurgery 118, 1367–1377.

Garyfallidis, E., 2012. Towards an accurate brain tractography. Ph.D. thesis. University of Cambridge.

Garyfallidis, E., Brett, M., Amirbekian, B., Rokem, A., Van Der Walt, S., Descoteaux, M., Nimmo-Smith, I., Contributors, D., 2014a. Dipy, a library for the analysis of diffusion MRI data. Frontiers in Neuroinformatics 8, 8.

Garyfallidis, E., Brett, M., Amirbekian, B., Rokem, A., van der Walt, S., Descoteaux, M., Nimmo-Smith, I., Dipy Contributors, 2014b. Dipy, a library for the analysis of diffusion MRI data. Frontiers in Neuroinformatics 8. URL: http://journal.frontiersin.org/article/10.3389/fninf.2014.00008/abstract, doi:10.3389/fninf.2014.00008.

Garyfallidis, E., Brett, M., Amirbekian, B., Rokem, A., van der Walt, S., Descoteaux, M., Nimmo-Smith, I., Dipy Contributors, 2014c. Dipy, a library for the analysis of diffusion MRI data. Front. Neuroinform. 8, 8.

Garyfallidis, E., Brett, M., Correia, M.M., Williams, G.B., Nimmo-Smith, I., 2012. Quickbundles, a method for tractography simplification. Frontiers in Neuroscience 6, 175.

Garyfallidis, E., Côté, M.A., Rheault, F., Sidhu, J., Hau, J., Petit, L., Fortin, D., Cunanne, S., Descoteaux, M., 2017. Recognition of white matter bundles using local and global streamline-based registration and clustering. NeuroImage.

Garyfallidis, E., Côté, M.A., Rheault, F., Sidhu, J., Hau, J., Petit, L., Fortin, D., Cunanne, S., Descoteaux, M., 2018. Recognition of white matter bundles using local and global streamline-based registration and clustering. NeuroImage 170, 283–295. doi:10.1016/j.neuroimage.2017.07.015.

Garyfallidis, E., Ocegueda, O., Wassermann, D., Descoteaux, M., 2015. Robust and efficient linear registration of white-matter fascicles in the space of streamlines. NeuroImage 117, 124–140.

Girard, G., Whittingstall, K., Deriche, R., Descoteaux, M., 2014. Towards quantitative connectivity analysis: reducing tractography biases. Neuroimage 98, 266–278.

Glozman, T., Bruckert, L., Pestilli, F., Yecies, D.W., Guibas, L.J., Yeom, K.W., 2018. Framework for shape analysis of white matter fiber bundles. NeuroImage 167, 466–477.

Goedert, M., Spillantini, M., Crowther, R., 1991. Tau proteins and neurofibrillary degeneration. Brain pathology 1, 279–286.

Gong, G., He, Y., Concha, L., Lebel, C., Gross, D.W., Evans, A.C., Beaulieu, C., 2008. Mapping anatomical connectivity patterns of human cerebral cortex using in vivo diffusion tensor imaging tractography. Cerebral cortex 19, 524–536.

Greene, C., Cieslak, M., Grafton, S.T., 2018. Effect of different spatial normalization approaches on tractography and structural brain networks. Network Neuroscience 2, 362–380.

Haines, J.D., Inglese, M., Casaccia, P., 2011. Axonal damage in multiple sclerosis. Mount Sinai Journal of Medicine: A Journal of Translational and Personalized Medicine 78, 231–243.

Hansen, C.B., Yang, Q., Lyu, I., Rheault, F., Kerley, C., Chandio, B.Q., Fadnavis, S., Williams, O., Shafer, A.T., Resnick, S.M., et al., 2021. Pandora: 4-d white matter bundle population-based atlases derived from diffusion mri fiber tractography. Neuroinformatics 19, 447–460.

Jahanshad, N., Kochunov, P.V., Sprooten, E., Mandl, R.C., Nichols, T.E., Almasy, L., Blangero, J., Brouwer, R.M., Curran, J.E., de Zubicaray, G.I., et al., 2013. Multi-site genetic analysis of diffusion images and voxelwise heritability analysis: A pilot project of the ENIGMA–DTI working group. Neuroimage 81, 455–469.

Jenkinson, M., Beckmann, C.F., Behrens, T.E., Woolrich, M.W., Smith, S.M., 2012. FSL. Neuroimage 62, 782–790.

Jeurissen, B., Tournier, J.D., Dhollander, T., Connelly, A., Sijbers, J., 2014. Multi-tissue constrained spherical deconvolution for improved analysis of multi-shell diffusion MRI data. NeuroImage 103, 411–426.

Kellner, E., Dhital, B., Kiselev, V.G., Reisert, M., 2016. Gibbs-ringing artifact removal based on local subvoxel-shifts. Magnetic Resonance in Medicine 76, 1574–1581.

Kubicki, M., McCarley, R.W., Shenton, M.E., 2005. Evidence for white matter abnormalities in schizophrenia. Current opinion in psychiatry 18, 121.

Kuhn, H.W., 1955. The hungarian method for the assignment problem. Naval research logistics quarterly 2, 83–97.

Landau, S., Thomas, B., Thurfjell, L., Schmidt, M., Margolin, R., Mintun, M., Pontecorvo, M., Baker, S., Jagust, W., Initiative, A.D.N., 2014. Amyloid pet imaging in alzheimer’s disease: a comparison of three radiotracers. European journal of nuclear medicine and molecular imaging 41, 1398–1407.

Landau, S.M., Breault, C., Joshi, A.D., Pontecorvo, M., Mathis, C.A., Jagust, W.J., Mintun, M.A., 2013a. Amyloid-β imaging with pittsburgh compound b and florbetapir: comparing radiotracers and quantification methods. Journal of Nuclear Medicine 54, 70–77.

Landau, S.M., Lu, M., Joshi, A.D., Pontecorvo, M., Mintun, M.A., Trojanowski, J.Q., Shaw, L.M., Jagust, W.J., Initiative, A.D.N., 2013b. Comparing positron emission tomography imaging and cerebrospinal fluid measurements of β-amyloid. Annals of neurology 74, 826–836.

Landau, S.M., Mintun, M.A., Joshi, A.D., Koeppe, R.A., Petersen, R.C., Aisen, P.S., Weiner, M.W., Jagust, W.J., Initiative, A.D.N., 2012. Amyloid deposition, hypometabolism, and longitudinal cognitive decline. Annals of neurology 72, 578–586.

Le Bihan, D., Mangin, J.F., Poupon, C., Clark, C.A., Pappata, S., Molko, N., Chabriat, H., 2001. Diffusion tensor imaging: concepts and applications. Journal of Magnetic Resonance Imaging: An Official Journal of the International Society for Magnetic Resonance in Medicine 13, 534–546.

Leemans, A., Jones, D.K., 2009. The B-matrix must be rotated when correcting for subject motion in DTI data. Magnetic Resonance in Medicine: An Official Journal of the International Society for Magnetic Resonance in Medicine 61, 1336–1349.

Leemans, A., Sijbers, J., De Backer, S., Vandervliet, E., Parizel, P., 2006. Multiscale white matter fiber tract coregistration: A new feature-based approach to align diffusion tensor data. Magnetic Resonance in Medicine: An Official Journal of the International Society for Magnetic Resonance in Medicine 55, 1414–1423.

Lizarraga, A., Lee, D., Kubicki, A., Sahib, A., Nunez, E., Narr, K., Joshi, S.H., 2021. Alignment of tractography streamlines using deformation transfer via parallel transport, in: International Workshop on Computational Diffusion MRI, Springer. pp. 96–105.

Manjón, J.V., Coupé, P., Concha, L., Buades, A., Collins, D.L., Robles, M., 2013. Diffusion weighted image denoising using overcomplete local PCA. PLoS ONE 8.

Marek, K., Jennings, D., Lasch, S., Siderowf, A., Tanner, C., Simuni, T., Coffey, C., Kieburtz, K., Flagg, E., Chowdhury, S., et al., 2011. The parkinson progression marker initiative (PPMI). Progress in neurobiology 95, 629–635.

Mori, S., Crain, B.J., Chacko, V.P., Van Zijl, P.C., 1999. Three-dimensional tracking of axonal projections in the brain by magnetic resonance imaging. Annals of Neurology: Official Journal of the American Neurological Association and the Child Neurology Society 45, 265–269.

Mori, S., Van Zijl, P.C., 2002. Fiber tracking: principles and strategies– a technical review. NMR in Biomedicine: An International Journal Devoted to the Development and Application of Magnetic Resonance In Vivo 15, 468–480.

Myronenko, A., Song, X., 2010. Point set registration: Coherent point drift. IEEE transactions on pattern analysis and machine intelligence 32, 2262–2275.

Myronenko, A., Song, X., Carreira-Perpinan, M., 2006. Non-rigid point set registration: Coherent point drift. Advances in neural information processing systems 19.

Nasrabady, S.E., Rizvi, B., Goldman, J.E., Brickman, A.M., 2018. White matter changes in alzheimer’s disease: a focus on myelin and oligodendrocytes. Acta neuropathologica communications 6, 1–10.

Olivetti, E., Gori, P., Astolfi, P., Bertó, G., Avesani, P., 2020. Nonlinear alignment of whole tractograms with the linear assignment problem, in: International Workshop on Biomedical Image Registration, Springer. pp. 3–11.

O’Donnell, L.J., Wells, W.M., Golby, A.J., Westin, C.F., 2012. Unbiased groupwise registration of white matter tractography, in: International Conference on Medical Image Computing and Computer-Assisted Intervention, Springer. pp. 123–130.

Papuć, E., Rejdak, K., 2018. The role of myelin damage in alzheimer’s disease pathology. Archives of Medical Science 16, 345–341.

Prineas, J.W., Barnard, R.O., Revesz, T., Kwon, E.E., Sharer, L., Cho, E.S., 1993. Multiple sclerosis: pathology of recurrent lesions. Brain 116, 681–693.

Rohlfing, T., 2006. Transformation model and constraints cause bias in statistics on deformation fields, in: International Conference on Medical Image Computing and Computer-Assisted Intervention, Springer. pp. 207–214.

Romero-Bascones, D., Qamar Chandio, B., Fadnavis, S., Sung Park, J., Koudoro, S., Ayala, U., Barrenechea, M., Garyfallidis, E., 2022. Bundleatlasing: unbiased population-specific atlasing of bundles in streamline space, in: Proc. ISMRM.

Rose, S.E., Chen, F., Chalk, J.B., Zelaya, F.O., Strugnell, W.E., Benson, M., Semple, J., Doddrell, D.M., 2000. Loss of connectivity in alzheimer’s disease: an evaluation of white matter tract integrity with colour coded mr diffusion tensor imaging. Journal of Neurology, Neurosurgery & Psychiatry 69, 528–530.

Sepulcre, J., Grothe, M.J., d’Oleire Uquillas, F., Ortiz-Terán, L., Diez, I., Yang, H.S., Jacobs, H.I., Hanseeuw, B.J., Li, Q., El-Fakhri, G., et al., 2018. Neurogenetic contributions to amyloid beta and tau spreading in the human cortex. Nature medicine 24, 1910–1918.

Smith, S.M., 2002. Fast robust automated brain extraction. Human Brain Mapping 17, 143–155.

Smith, S.M., Jenkinson, M., Johansen-Berg, H., Rueckert, D., Nichols, T.E., Mackay, C.E., Watkins, K.E., Ciccarelli, O., Cader, M.Z., Matthews, P.M., et al., 2006. Tract-based spatial statistics: voxelwise analysis of multi-subject diffusion data. Neuroimage 31, 1487–1505.

Sotiropoulos, S.N., Jbabdi, S., Xu, J., Andersson, J.L., Moeller, S., Auerbach, E.J., Glasser, M.F., Hernandez, M., Sapiro, G., Jenkinson, M., et al., 2013. Advances in diffusion mri acquisition and processing in the human connectome project. Neuroimage 80, 125–143.

St-Jean, S., Chamberland, M., Viergever, M.A., Leemans, A., 2019. Reducing variability in along-tract analysis with diffusion profile realignment. Neuroimage 199, 663–679.

St-Onge, E., Garyfallidis, E., Collins, D.L., 2021. Fast tractography streamline search, in: International Workshop on Computational Diffusion MRI, Springer. pp. 82–95.

Thomopoulos, S.I., Nir, T.M., Reina, J.E.V., Jahanshad, N., Thompson, P.M., 2020. Diffusion MRI metrics of brain microstructure in alzheimer’s disease: Boosting disease sensitivity with multi-shell imaging and advanced pre-processing: Neuroimaging/new imaging methods. Alzheimer’s & Dementia 16, e046654.

Tournier, J.D., Calamante, F., Connelly, A., 2007. Robust determination of the fibre orientation distribution in diffusion MRI: Non-negativity constrained super-resolved spherical deconvolution. NeuroImage 35, 1459–1472.

Tournier, J.D., Smith, R., Raffelt, D., Tabbara, R., Dhollander, T., Pietsch, M., Christiaens, D., Jeurissen, B., Yeh, C.H., Connelly, A., 2019. MR-trix3: a fast, flexible and open software framework for medical image processing and visualisation. Neuroimage 202, 116137.

Tsang, J., Dougherty, R., Wandell, B., 2010. Tract alignment errors decrease detection power in group analyses of diffusion data with tbss. Society for Neuroscience. San Diego, CA.

Tustison, N.J., Avants, B.B., Cook, P.A., Zheng, Y., Egan, A., Yushkevich, P.A., Gee, J.C., 2010. N4itk: improved N3 bias correction. IEEE transactions on medical imaging 29, 1310–1320.

Tustison, N.J., Cook, P.A., Holbrook, A.J., Johnson, H.J., Muschelli, J., Devenyi, G.A., Duda, J.T., Das, S.R., Cullen, N.C., Gillen, D.L., et al., 2021. The antsx ecosystem for quantitative biological and medical imaging. Scientific reports 11, 1–13.

Villemagne, V.L., Doré, V., Burnham, S.C., Masters, C.L., Rowe, C.C., 2018. Imaging tau and amyloid-β proteinopathies in alzheimer disease and other conditions. Nature Reviews Neurology 14, 225–236.

Wang, J., 2022. Fast non-rigid whole brain streamline matching based on fast fiber k-nn and group-wise mds.

Wassermann, D., Rathi, Y., Bouix, S., Kubicki, M., Kikinis, R., Shenton, M., Westin, C.F., 2011. White matter bundle registration and population analysis based on gaussian processes, in: Biennial International Conference on Information Processing in Medical Imaging, Springer. pp. 320– 332.

Whitford, T.J., Ford, J.M., Mathalon, D.H., Kubicki, M., Shenton, M.E., 2012. Schizophrenia, myelination, and delayed corollary discharges: a hypothesis. Schizophrenia bulletin 38, 486–494.

Yang, X., Shen, X., Long, J., Chen, H., 2012. An improved median-based Otsu image thresholding algorithm. Aasri Procedia 3, 468–473.

Yang, Y., Schilling, K., Shashikumar, N., Jasodanand, V., Moore, E.E., Pechman, K.R., Bilgel, M., Beason-Held, L.L., An, Y., Shafer, A., et al., 2023. White matter microstructural metrics are sensitively associated with clinical staging in alzheimer’s disease. Alzheimer’s & Dementia: Diagnosis, Assessment & Disease Monitoring 15, e12425.

Yeatman, J.D., Dougherty, R.F., Myall, N.J., Wandell, B.A., Feldman, H.M., 2012. Tract Profiles of White Matter Properties: Automating Fiber-Tract Quantification. PLoS ONE 7, e49790.

Yeh, F.C., Panesar, S., Fernandes, D., Meola, A., Yoshino, M., Fernandez-Miranda, J.C., Vettel, J.M., Verstynen, T., 2018a. Population-averaged atlas of the macroscale human structural connectome and its network topology. NeuroImage 178, 57–68.

Yeh, F.C., Panesar, S., Fernandes, D., Meola, A., Yoshino, M., Fernandez-Miranda, J.C., Vettel, J.M., Verstynen, T., 2018b. Population-averaged atlas of the macroscale human structural connectome and its network topology. NeuroImage 178, 57–68. URL: https://linkinghub.elsevier.com/retrieve/pii/S1053811918304324, doi:10.1016/j.neuroimage.2018.05.027.

Yendiki, A., Panneck, P., Srinivasan, P., Stevens, A., Zöllei, L., Augustinack, J., Wang, R., Salat, D., Ehrlich, S., Behrens, T., et al., 2011. Automated probabilistic reconstruction of white-matter pathways in health and disease using an atlas of the underlying anatomy. Frontiers in neuroinformatics 5, 23.

Yuille, A.L., Grzywacz, N.M., 1988. The motion coherence theory, in: ICCV.

Yuille, A.L., Grzywacz, N.M., 1989. A mathematical analysis of the motion coherence theory. International Journal of Computer Vision 3, 155–175.

Zavaliangos-Petropulu, A., Nir, T.M., Thomopoulos, S.I., Reid, R.I., Bernstein, M.A., Borowski, B., Jack Jr., C.R., Weiner, M.W., Jahanshad, N., Thompson, P.M., 2019. Diffusion MRI Indices and Their Relation to Cognitive Impairment in Brain Aging: The Updated Multi-protocol Approach in ADNI3. Frontiers in Neuroinformatics 13, 2. URL: https://www.frontiersin.org/article/10.3389/fninf.2019.00002/full, doi:10.3389/fninf.2019.00002.

Zhang, F., Wells, W.M., O’Donnell, L.J., 2021. Deep diffusion mri registration (ddmreg): a deep learning method for diffusion mri registration. IEEE Transactions on Medical Imaging 41, 1454–1467.

Zvitia, O., Mayer, A., Shadmi, R., Miron, S., Greenspan, H.K., 2009. Coregistration of white matter tractographies by adaptive-mean-shift and gaussian mixture modeling. IEEE Transactions on Medical Imaging 29, 132–145.

