## Supplement for "BundleWarp: Enhancing White Matter Tractometry and Morphometry with Precise Neuronal Mapping using Streamline-based Nonlinear Registration"

**This PDF file includes:**

Text S1

Figs. S1 to S8

### S1 Text S1

#### S1.1 Streamline-based Distances and Cost Functions

Here, we explain streamline-based distances and cost functions that will be used throughout the manuscript. These functions are either used for optimization or for quantitative assessment of the methods.

##### S1.1.1 Minimum Direct Flip Distance

The minimum average direct-flip distance (MDF) was introduced in the QuickBundles paper Garyfallidis, Brett, Correia, Williams and Nimmo-Smith (2012) and is widely used for calculating a distance between two streamlines that have the same number of points. It is defined as follows:
Let $S_1$ and $S_2$ be two streamlines with the same number of points.

$$d(S_1, S_2) = \frac{1}{k}\sum_{i=1}^{K} |S_{1i} - S_{2i}|$$

$$d_{direct}(S_1, S_2) = d(S_1, S_2)$$

$$d_{flipped}(S_1, S_2) = d(S_1, S_2^F) = d(S_1^F, S_2)$$

$$MDF(S_1, S_2) = min(d_{direct}(S_1, S_2), d_{flipped}(S_1, S_2))$$

##### S1.1.2 Bundle-based Minimum Distance

The bundle-based minimum distance (BMD) was introduced and is used as a cost function in the streamline-based linear registration (SLR) method Garyfallidis, Ocegueda, Wassermann and Descoteaux (2015). It is used for calculating the distance between two bundles (two sets of streamlines) in millimeters. The BMD is defined as follows:
Let $B_1$ and $B_2$ be bundles where $n$ and $m$ represent the total number of streamlines in each bundle, respectively. And $S_{1i}$ represents $i^{th}$ streamline in bundle $B_1$ and $S_{2j}$ represents $j^{th}$ streamline in bundle $B_2$. A rectangular matrix D is populated with all pairwise MDF streamline distances. Every element of the D matrix is calculated by taking the MDF distance of $S_{1i}$ streamline of $B_1$ bundle with $S_{2j}$ streamline of $B_2$ bundle, $D_{i,j} = MDF(S_{1i}, S_{2j})$. The most similar streamlines from one

bundle to the other are given more weightage by the MDF values of the rows and columns of the D matrix.

$$BMD(B_1, B_2) = \frac{1}{2}\left(\frac{1}{n_1}\sum_{i=1}^{n_1} \min_j D(i,j) + \frac{1}{n_2}\sum_{j=1}^{n_2} \min_i D(i,j)\right)^2$$

##### S1.1.3 Bundle Shape Similarity Score

Bundle shape similarity score (SM) introduced in the BUAN method Chandio, Risacher, Pestilli, Bullock, Yeh, Koudoro, Rokem, Harezlak and Garyfallidis (2020) as a quality control measure for assessing the quality of a large number of extracted bundles coming from different subjects and populations. SM internally uses bundle adjacency metric (BA) Garyfallidis et al. (2012) to calculate bundle shape similarity between two bundles.

Let B1 and B2 be two bundles, and $\theta > 0$ be a selected adjacency threshold. Where $S1_i$ is a streamline in $B1$ and $S2_i$ is a streamline belonging to $B2$. $S1_i$ is adjacent to $B2$ if $MDF(S1_i, S2_i) \leq \theta$.

Coverage of $B1$ by $B2$ is defined as following:

$coverage(B1, B2) = \frac{\text{number of adjacent streamlines}}{\text{total number of streamlines in B1}}$

SM/BA is defined as follows:

$$SM(B1, B2) = BA(B1, B2) = 0.5(coverage(B1, B2) + coverage(B2, B1))$$

SM ranges between 0 and 1, with 0 being the lowest score (completely different shape) and 1 being the highest score (extremely close in shape).

#### S1.2 BundleWarp Execution Time

Different types of bundles have different sizes and can be composed of different numbers of streamlines. Moreover, even two bundles of the same type can have a varying number of streamlines. Here, we investigate BundleWarp execution time based on the number of streamlines in static and moving bundles on a system with 32 GB RAM and one Intel Core i7-7700K CPU with 8 cores.

Fig. S1 shows the execution time of BundleWarp when the static bundle's streamline count is kept fixed at 1000 and the moving bundle's streamline count increases. Here, the x-axis has streamline count in the moving bundle, and the y-axis has BundleWarp execution time in minutes. One of the most common applications of BundleWarp would be to register different subjects' bundles to an atlas bundle, where the static bundle (atlas bundle) would always be the same, and the moving bundle would change. Most of the time, extracted bundles have a streamline count of less than 4,000. We can see from our experiments in Fig. S1 that BundleWarp is fast, and execution time is a few seconds, and would be less than a minute for bundles of typical size bundles (streamlines $<$ 4,000).

Fig. S2 shows the execution time of BundleWarp when both static and moving bundles' streamline count increases. Here, the x-axis has streamline count in both static and moving bundles, and the y-axis has BundleWarp execution time in minutes. Here, we investigate the BundleWarp execution time when the streamline count increases in both static and moving bundles. We can see that BundleWarp execution time in seconds when both bundles have a streamline count of less than 4,000. The execution time increases when the streamline count increases up to 10,000. Even at a higher number of streamlines in both bundles, BundleWarp only takes a few minutes (less than 3 minutes). Whereas, if we try to register two whole-brain tractograms, the high execution time would increase significantly. However, bundles are typically comprised of 4,000 or fewer streamlines, which at most is 1 minute.

We are able to perform deformable streamline-based registration due to the proposed approach of finding streamline correspondence in two bundles and then applying mlCPD. mlCPD performs deformations and is extremely fast for two small point clouds of two streamlines.

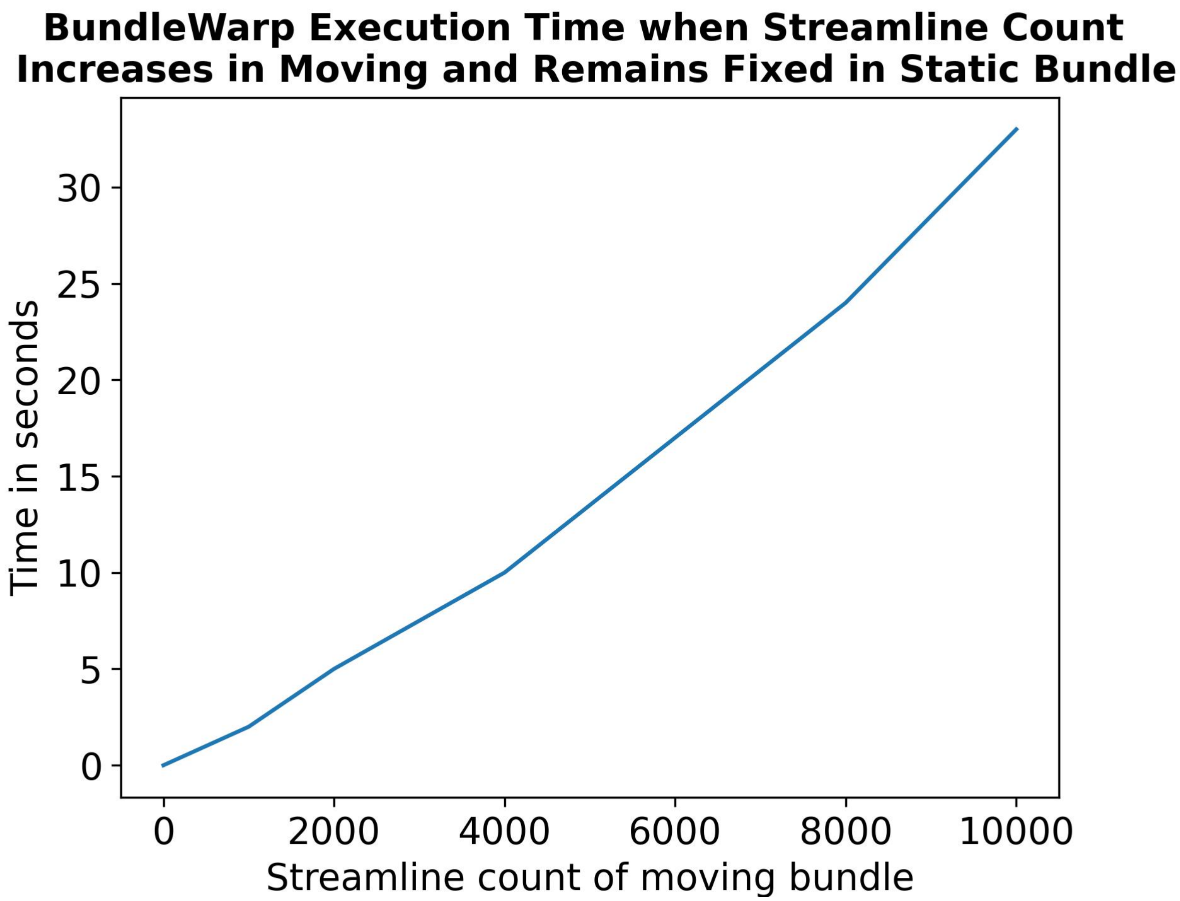


**Figure S1.** Execution time of BundleWarp in minutes when streamline count increases in moving bundle and remains fixed (1000 streamlines) in static bundle. The x-axis shows the count of streamlines in a moving bundle, where a static bundle always has 2010 streamlines. The y-axis shows BundleWarp execution time for registration of one pair of bundles.

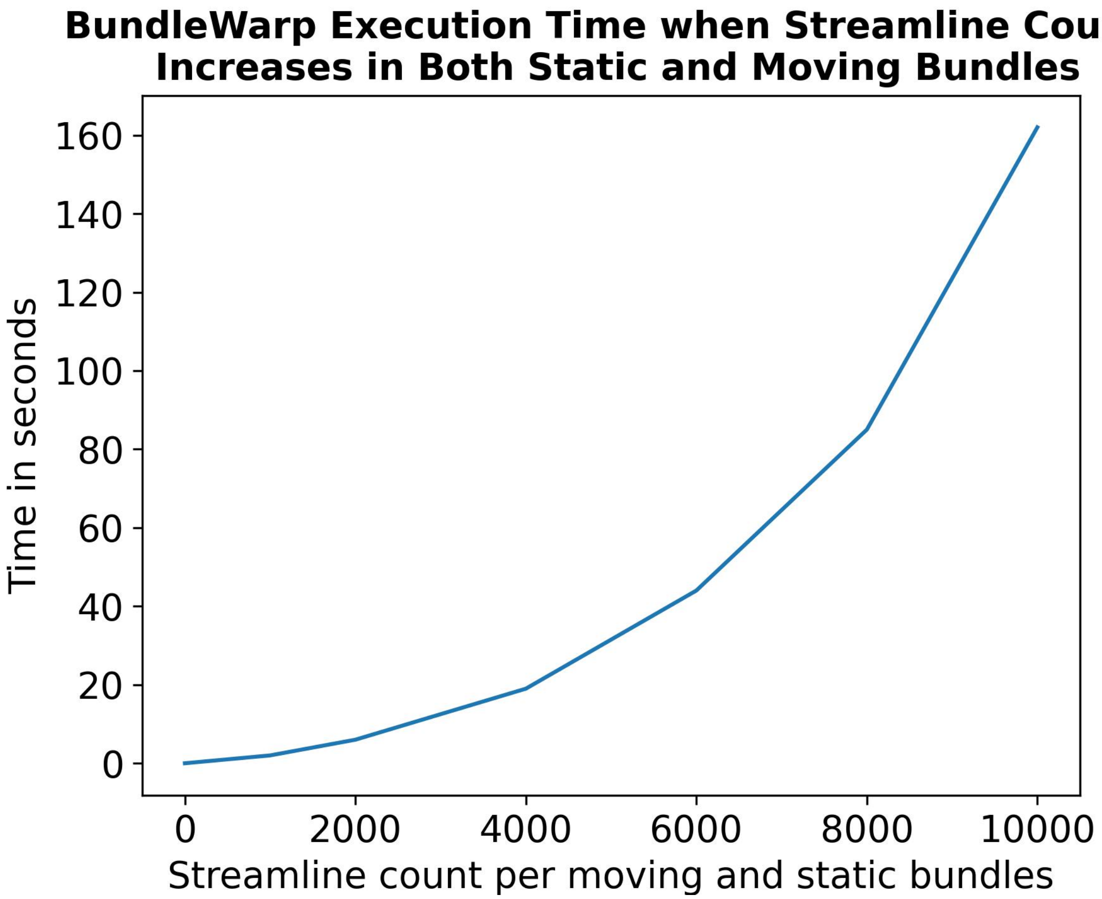


**Figure S2.** Execution time of BundleWarp in minutes when streamline count increases in both static and moving bundles. The x-axis shows the count of streamlines in each bundle, and the y-axis shows BundleWarp execution time for registration of one pair of bundles.

**White Matter Tracts Used in the Paper**

**Projection Pathways**

| Abbreviation | Full Name |
|---|---|
| CST_L/R | Corticospinal Tract - Left / Right |
| ORL_/R | Optic Radiation - Left / Right |
| FPT_L/R | Frontopontine Tract - Left / Right |
| OPT_L/R | Occipito Pontine Tract - Left / Right |

**Commissural Pathways**

| Abbreviation | Full Name |
|---|---|
| CCMid | Corpus Callosum Middle |
| CCForcepsMajor | Corpus Callosum Major |
| CCForcepsMinor | Corpus Callosum Minor |

**Cerebellum Pathways**

| Abbreviation | Full Name |
|---|---|
| V | Verma |

**Association Pathways**

| Abbreviation | Full Name |
|---|---|
| AF_L/R | Arcuate Fasciculus - Left / Right |
| EMC_L/R | Extreme Capsule - Left / Right |
| IFOF_L/R | Inferior Fronto-occipital Fasciculus - Left / Right |
| ILF_L/R | Inferior Longitudinal Fasciculus- Left / Right |
| MdLF_L/R | Middle Longitudinal Fasciculus - Left / Right |
| UF_L/R | Uncinate Fasciculus - Left / Right |

**Brainstem Pathways**

| Abbreviation | Full Name |
|---|---|
| ML_L/R | Medial Lemniscus - Left / Right |
| MLF_L/R | Medial Longitudinal fasciculus - Left / Right |
| STT_L/R | Spinothalamic Tract - Left / Right |

**Figure S3.** Full names of white matter tracts used in the paper.

#### S1.3 BundleWarp Method Comparison

For quantitative results, we used two streamline-based metrics, bundle shape similarity metric (SM)Garyfallidis et al. (2012); Chandio et al. (2020) and bundle-based minimum distance (BMD)Garyfallidis et al. (2015). We also used the volumetric dice similarity metric (DICE) to compare the overlap of fiber density images of tracts. Fig. S4, Fig. S5, and Fig. S6 present a quantitative comparison of methods SM, BMD, and DICE, respectively. The top panels in all three figures show three matrices of the respective metric for three registration methods, ANTs, SLR, and BW. Twenty-seven bundles extracted from 64 subjects (1,728 total bundles) were registered to 27 atlas bundles of the same type. Three metrics were calculated between static and moved (registered) bundles. Here, rows represent subjects, and columns represent bundle types. Each pixel in the matrices indicates the respective metric's score among the atlas bundle and registered bundle (static and moved).

In Fig. S4, results are presented for the SM metric, which is a bounded metric and ranges between 0 and 1. Darker blue (closer to 1) means higher shape similarity, implying better registration performance. Conversely, white or light blue color (closer to 0) implies a lower value of SM and lower shape similarity between bundles. The bottom panel shows the mean SM score per bundle for each method. Twenty-seven bundle names are listed on the left of the bottom panel. BW method has the highest SM score for all bundles. SLR performs second best for all bundles except for the Verma (V) bundle, while ANTs registration has the lowest bundle shape similarity.

In Fig. S5, results are presented for the BMD metric, which represents the distance between two bundles. A lower distance indicates two bundles are closely aligned, and a higher distance means two bundles are far from each other. In the BMD matrices, a darker brown color signifies a lower distance, hence better registration performance. In contrast, a lighter brown color signifies a higher BMD distance between bundles. The bottom panel shows the mean BMD score per bundle for each method. BW method has the lowest BMD score for all bundles (which is desirable). SLR performs second best for all bundles and has the second lowest BMD score. ANTs registration has the highest BMD score, implying comparatively suboptimal alignment of bundles.

In Fig. S6, results are presented for the DICE metric, which is a bounded metric and ranges between 0 and 1. A darker purple color (closer to 1) means a higher DICE score, hence better registration performance. White or light purple color implies a lower value of DICE and lower volumetric overlap between two bundles. The bottom panel shows the mean DICE score per bundle for each method. BW method has the highest DICE score for all bundles. SLR performs second best. ANTs registration has the lowest dice similarity.

#### S1.4 BUAN Tractometry Results with BundleWarp

This paper primarily focuses on evaluating the impact of BundleWarp within the BUAN tractometry framework, with the majority of results comparing BUAN outcomes across four experiments, both with and without BundleWarp. However, we also highlight specific tracts significantly affected by amyloid and tau positivity. Amyloid and tau are critical biomarkers in neurodegenerative diseases, and their presence is closely associated with cognitive decline. These biomarkers represent the earliest pathological changes in Alzheimer's disease, often appearing years before the onset of clinical symptoms. Detecting subtle white matter changes associated with amyloid and tau is crucial for understanding preclinical disease progression and developing interventions aimed at halting or slowing disease advancement in its earliest stages. The application of BundleWarp in BUAN allows for more precise detection of tract-specific microstructural alterations linked to these biomarkers, revealing associations that may be overlooked with conventional methods.

We present BUAN results that highlight the effect of amyloid positivity on white matter tracts, based on data from 329 amyloid-negative participants (CN: 235, MCI: 86, Dementia: 8) (156M, 173F) and 277 amyloid-positive participants (CN: 139, MCI: 87, Dementia: 51) (131M, 146F) from the ADNI3 dataset. Fig.S7 shows the significant impact of amyloid positivity on MD and RD in the corpus callosum major. Alterations in AD and MD in the left cingulum tract, as well as

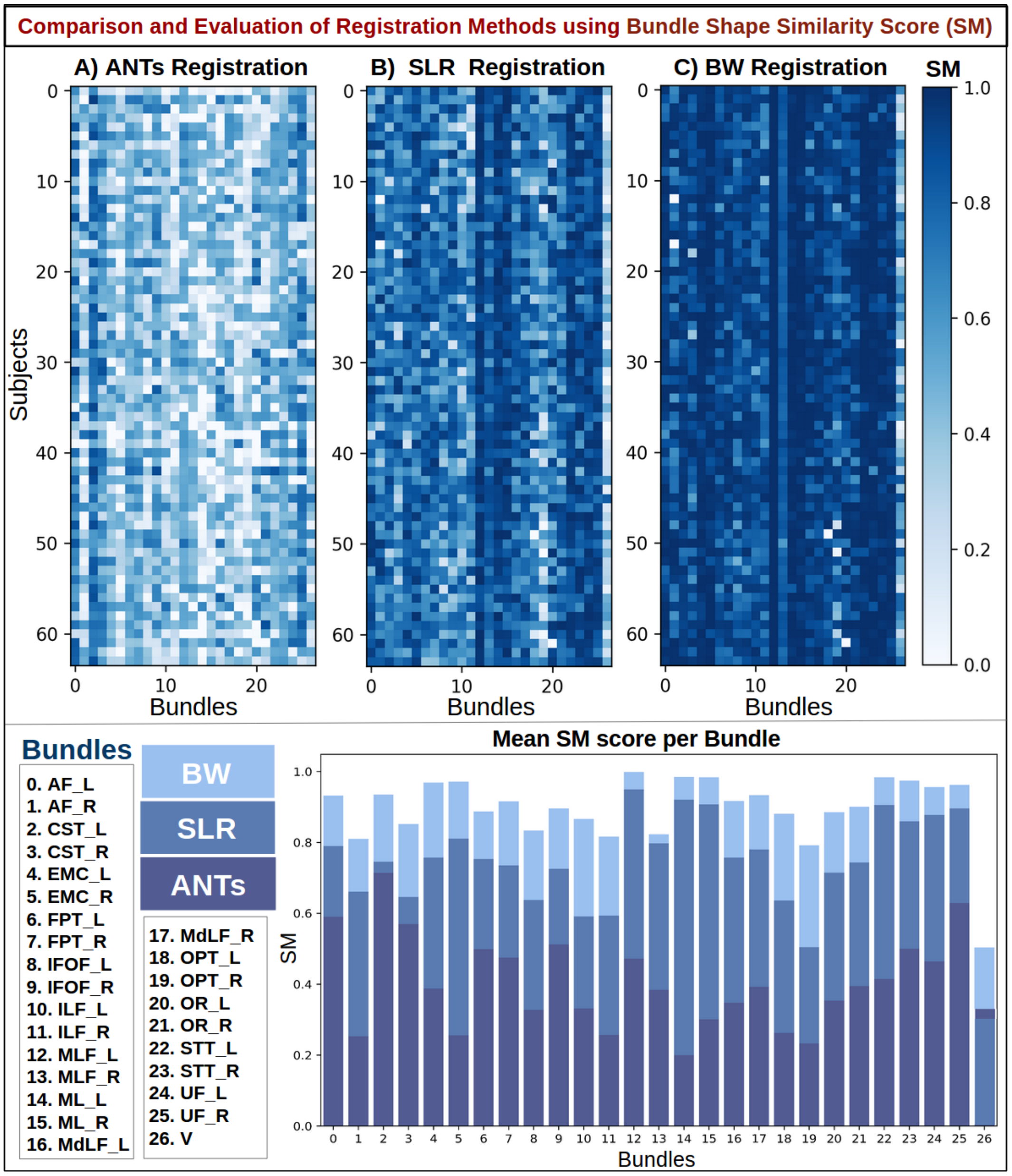


**Figure S4. Quantitative results using bundle shape similarity metric (SM) to compare Ants, SLR, and BundleWarp (BW) methods**. The top panel shows three SM matrices for three registration methods. 27 bundles extracted from 64 subjects (1,728 total bundles) were registered to 27 atlas bundles of the same type. Here, rows represent the subject, and columns represent the bundle type. Each pixel in the matrices indicates the SM score among the atlas bundle and registered bundle (static and moved). Darker blue means higher shape similarity, hence better registration performance. The bottom panel shows the mean SM score per bundle for each method and bundle names. BW method has the highest SM score for all bundles.

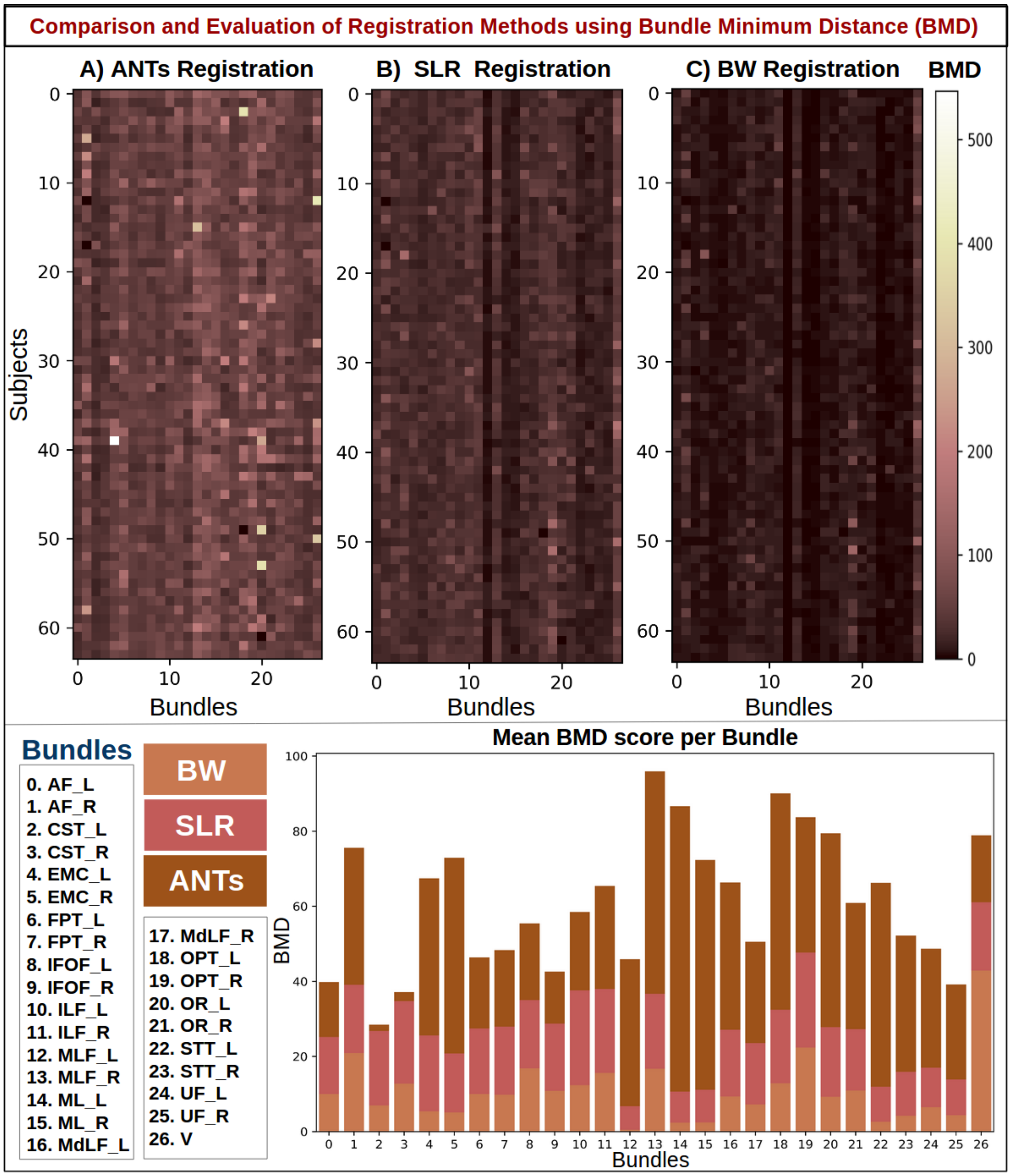


**Figure S5. Quantitative results using bundle-based minimum distance (BMD) to compare Ants, SLR, and BundleWarp (BW) methods**. The top panel shows three BMD matrices for three registration methods. 27 bundles extracted from 64 subjects (1,728 total bundles) were registered to 27 atlas bundles of the same type. Here, rows represent the subject, and columns represent the bundle type. Each pixel in the matrices indicates the BMD distance between the atlas bundle and the registered bundle (static and moved). Darker brown means lower bundle distances, hence better registration performance. The bottom panel shows the mean BMD distance per bundle for each method and bundle names. BW method has the smallest BMD for all bundles.

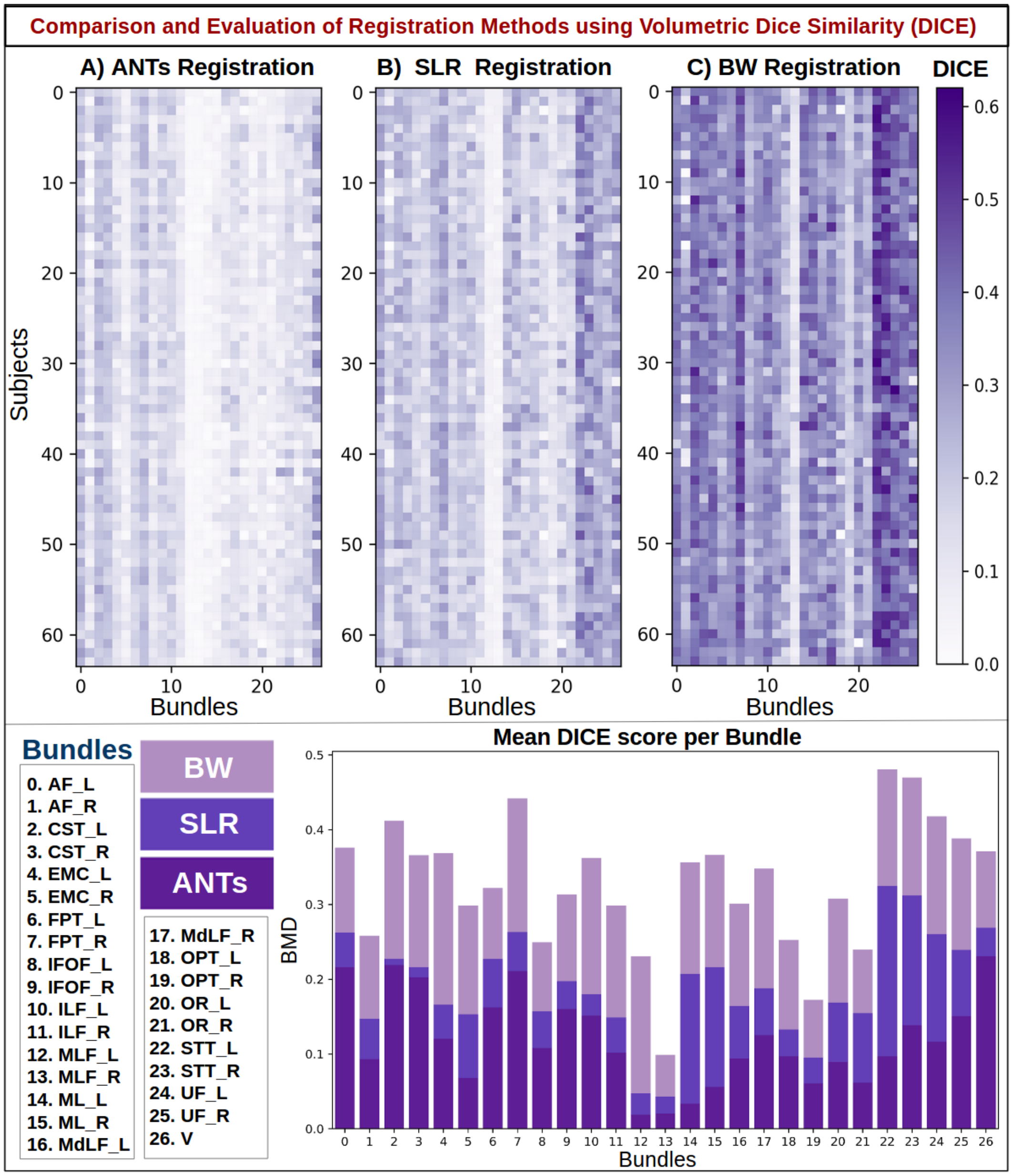


**Figure S6. Quantitative results using volumetric dice similarity score (DICE) to compare Ants, SLR, and BundleWarp (BW) methods**. The top panel shows three DICE matrices for three registration methods. 27 bundles extracted from 64 subjects (1,728 total bundles) were registered to 27 atlas bundles of the same type. Density maps of atlas bundles and registered bundles were created and used for calculating the volumetric DICE score. Here, rows represent the subject and columns represent bundle type. Each pixel in the matrices indicates the DICE score between the atlas bundle and the registered bundle (static and moved). Darker purple means a higher DICE score, hence better registration performance. The bottom panel shows the mean DICE score per bundle for each method and bundle names. BW method has the highest DICE for all bundles.

MD changes in the corpus callosum mid and extreme capsule left, are also presented in Figure S7. Dark purple colors indicate lower $p$-values and a stronger association. Segments with $p$-values $<$ 0.05 are highlighted by a red outline on the tracts. In significant tracts, diffusivity metrics increase while fractional anisotropy decreases in those with higher levels of amyloid pathology, aligning with the established effects of dementia on these metrics Nir, Villalón-Reina, Salminen, Haddad, Zheng, Thomopoulos, Jack Jr, Weiner, Thompson, Jahanshad et al. (2023); Chandio, Villalon-Reina, Nir, Thomopoulos, Feng, Benavidez, Jahanshad, Harezlak, Garyfallidis, Thompson et al. (2024b); Chandio, Villalon-Reina, Nir, Thomopoulos, Feng, Benavidez, Jahanshad, Harezlak, Garyfallidis and Thompson (2024a)

We ran BUAN tractometry to assess the effect of tau positivity on white matter tracts, using data from 401 tau-negative participants (CN: 293, MCI: 95, Dementia: 13) (192M, 209F) and 168 tau-positive participants (CN: 60, MCI: 68, Dementia: 40) (75M, 93F) in the ADNI3 dataset. Fig.S8 illustrates significant microstructural alterations due to tau positivity in MD and RD for the left and right cingulum tracts, as well as in FA and RD for the left arcuate fasciculus, and AD and RD for the left medial longitudinal fasciculus. Dark blue colors imply lower $p$-values and greater strength of association. The red outlines on the 3D tracts mark the segments that were significant, with $p$-values $< 0.05$.

In significant tracts, most diffusivity metrics show an increase, while fractional anisotropy decreases, consistent with expected patterns of microstructural abnormalities previously reported in dementia Nir et al. (2023); Chandio et al. (2024b). However, unlike prior work that primarily examined cortical measures or ROI-based diffusion changes, our analysis focuses directly on tract-level and along-tract microstructural and structural (shape) variation. This enables the detection of localized abnormalities within individual white-matter pathways. Notably, in certain tracts, axial diffusivity exhibits nonuniform alterations along the tract length, underscoring the value of tract-specific and spatially resolved analysis for revealing subtle, regionally varying disease effects.

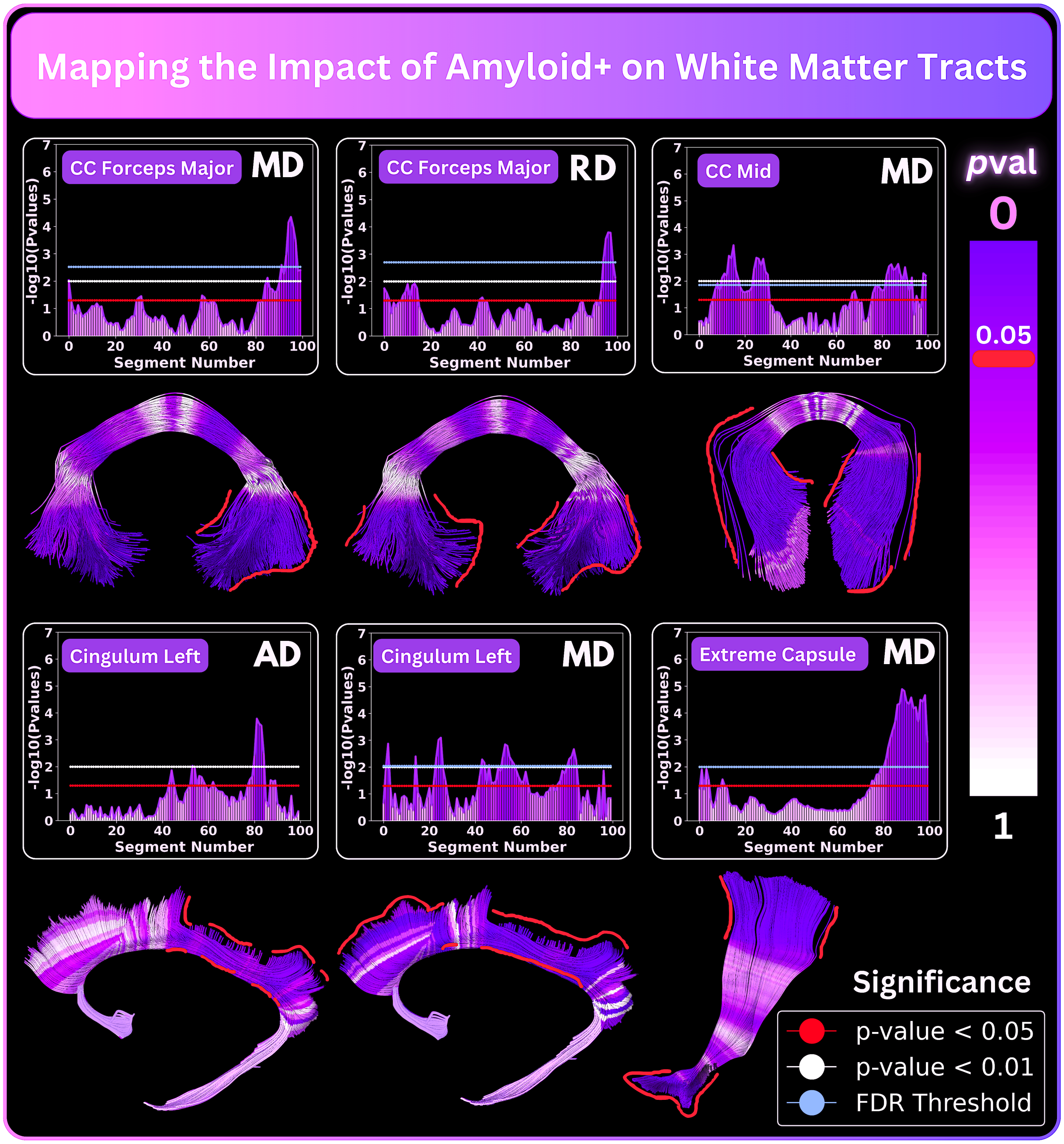


**Figure S7.** BundleWarp integrated into BUAN reveals alterations in white matter tracts due to amyloid-positive load. The first and third rows show $p$-value plots for each tract, where the x-axis represents the segment number along the tract and the y-axis shows a negative logarithm of $p$-values. The blue horizontal line in the plots represents the FDR-corrected threshold. Segments that pass the FDR-corrected threshold are considered significant. The second and fourth rows visualize $p$-values mapped onto the 3D tracts. Dark purple colors imply lower $p$-values and greater strength of association. Segments with $p$-values$<0.05$ are highlighted by the red outline on tracts.

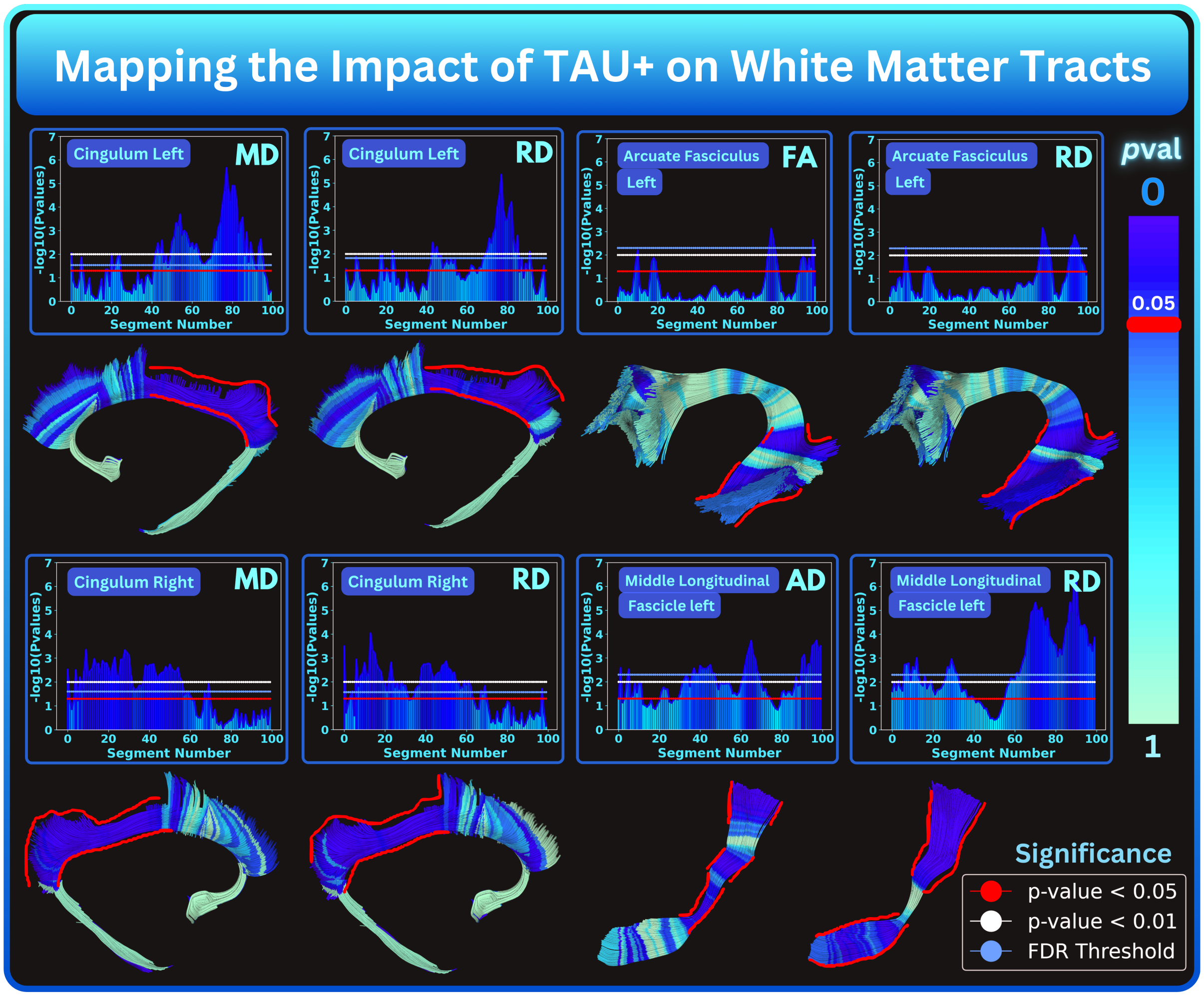


**Figure S8.** BundleWarp integrated into BUAN reveals white matter tract alteration due to tau positive plaque. The first and third rows show $p$-value plots for each tract, where the x-axis represents the segment number along the tract and the y-axis shows a negative logarithm of $p$-values. The blue horizontal line in the plots represents the FDR-corrected threshold. Segments that pass the FDR-corrected threshold are considered significant. The second and fourth rows visualize $p$-values mapped onto the 3D tracts. Dark blue colors imply lower $p$-values and greater strength of association. Segments with $p$-values$<$0.05 are highlighted by the red outline on tracts.

### References


Chandio, B.Q., Risacher, S.L., Pestilli, F., Bullock, D., Yeh, F.C., Koudoro, S., Rokem, A., Harezlak, J., Garyfallidis, E., 2020. Bundle analytics, a computational framework for investigating the shapes and profiles of brain pathways across populations. Scientific Reports 10, 17149.

Chandio, B.Q., Villalon-Reina, J.E., Nir, T.M., Thomopoulos, S.I., Feng, Y., Benavidez, S., Jahanshad, N., Harezlak, J., Garyfallidis, E., Thompson, P.M., 2024a. Bundle analytics based data harmonization for multi-site diffusion mri tractometry. bioRxiv , 2024–02.

Chandio, B.Q., Villalon-Reina, J.E., Nir, T.M., Thomopoulos, S.I., Feng, Y., Benavidez, S., Jahanshad, N., Harezlak, J., Garyfallidis, E., Thompson, P.M., et al., 2024b. Amyloid, tau, and apoe in alzheimer's disease: Impact on white matter tracts. bioRxiv .

Garyfallidis, E., Brett, M., Correia, M.M., Williams, G.B., Nimmo-Smith, I., 2012. Quickbundles, a method for tractography simplification. Frontiers in Neuroscience 6, 175.

Garyfallidis, E., Ocegueda, O., Wassermann, D., Descoteaux, M., 2015. Robust and efficient linear registration of white-matter fascicles in the space of streamlines. NeuroImage 117, 124–140.

Nir, T.M., Villalón-Reina, J.E., Salminen, L.E., Haddad, E., Zheng, H., Thomopoulos, S.I., Jack Jr, C.R., Weiner, M.W., Thompson, P.M., Jahanshad, N., et al., 2023. Cortical microstructural associations with csf amyloid and ptau. Molecular Psychiatry , 1–12.
